# Uncovering the ligandome of low-density lipoprotein receptor-related protein 1 in cartilage: a top-down approach to identify therapeutic targets

**DOI:** 10.1101/2022.03.02.482546

**Authors:** Kazuhiro Yamamoto, Carsten Scavenius, Maria M Meschis, Emilie H Mogensen, Abdulrahman M E Gremida, Ida B Thøgersen, Simone Bonelli, Simone D Scilabra, Salvatore Santamaria, Josefin Ahnström, George Bou-Gharios, Jan J Enghild, Hideaki Nagase

## Abstract

The low-density lipoprotein receptor-related protein 1 (LRP1) is a cell-surface receptor ubiquitously expressed in adult tissues. It plays tissue-specific physiological roles by mediating endocytosis of a diverse range of extracellular molecules. Dysregulation of LRP1 is involved in multiple conditions including Alzheimer’s disease, atherosclerosis and osteoarthritis (OA). However, little information is available about the specific ligand profile (ligandome) for each tissue, which would lead to better understanding of its role in disease states. Here, we investigated adult articular cartilage where impaired LRP1-mediated endocytosis leads to tissue destruction. We used a top-down approach involving analysis of human chondrocyte secretome, direct binding assays and validation in LRP1-deficient fibroblasts, as well as a novel *Lrp1* conditional knockout (KO) mouse model. We found that inhibition of LRP1-mediated endocytosis results in cell death, alteration of the entire secretome and transcriptional modulations in human chondrocytes. We have identified more than 50 novel ligand candidates and confirmed direct LRP1 binding of *HGFAC*, *HMGB1*, *HMGB2*, *CEMIP*, *SLIT2*, *ADAMTS1*, *IGFBP7*, *SPARC* and *LIF*. Our *in vitro* endocytosis assay revealed the correlation of their affinity for LRP1 and the rate of endocytosis. Moreover, a conditional LRP1 KO mouse model demonstrated a critical role of LRP1 in regulating the high-affinity ligands in cartilage *in vivo*. This systematic approach revealed the extent of the chondrocyte LRP1 ligandome and identified potential novel therapeutic targets for OA.

## Introduction

Endocytosis is the process by which cells control the composition of their cell surface and extracellular environment and is thus essential for cellular signaling and metabolism. The low-density lipoprotein receptor-related protein 1 (LRP1, CD91) is a type 1 transmembrane protein consisting of a 515 kDa heavy-chain containing the extracellular ligand-binding domains, and a non-covalently associated 85 kDa light-chain containing a transmembrane and cytoplasmic domain. LRP1 is ubiquitously expressed in different tissues and cell types [1, 2], and mediates clathrin-dependent endocytosis of structurally and functionally diverse array of molecules including lipoproteins, extracellular matrix (ECM) proteins, growth factors, cell surface receptors, proteinases, proteinase inhibitors and secreted intracellular proteins [3, 4]. LRP1 is not only involved in ligand uptake but also in modulation of cellular signalling pathways by binding extracellular ligands including growth factors and subsequent intracellular interaction with scaffolding and adaptor proteins. The LRP1 intracellular domain contains two tyrosine phosphorylation sites, which can directly transmit the signals [5].

Ubiquitous deletion of the *Lrp1* gene in mice results in early embryonic lethality at E13.5 [6] due to extensive haemorrhaging, resulting from a failure to recruit and maintain smooth muscle cells (SMCs) and pericytes in the vasculature [7]. The selective deletion of LRP1 in neurons [8], macrophages [9, 10], hepatocytes [11, 12], SMCs [13, 14], endothelial cells [15] or adipocytes [16] all lead to significant phenotypic alterations. These studies revealed various biological roles of LRP1 including lipoprotein metabolism, insulin signalling, vascular wall integrity, inflammation, and the turnover of ECM components. Unsurprisingly, its dysregulation is linked to several diseases such as Alzheimer’s disease, atherosclerosis, cancer, and osteoarthritis (OA) [5, 17–19].

OA is a major cause of pain and disability and is the most prevalent age-related degenerative joint disease. With an expanding elderly population, OA imposes a major socio-economic burden on society [20]. Despite its prevalence, there is currently no disease-modifying OA treatment available, except joint replacement surgery at the late stage of the disease. The hallmark of OA is loss of articular cartilage, which results in impairment of joint function. Chondrocytes, as the only cell type present in articular cartilage, are essential for balancing in anabolic and catabolic processes. Therefore, chondrocytes are key regulators of cartilage homeostasis and joint health. Our previous study has shown that LRP1 plays a key major role in maintaining the ECM turnover of healthy adult cartilage by controlling extracellular levels of matrix-degrading metalloproteinases such as aggrecanases [21, 22], collagenases [23, 24], and their endogenous inhibitor tissue inhibitor of metalloproteinases (TIMP)3 [25, 26]. However, this endocytic process is impaired in cartilage with OA, resulting in prolonged presence of LRP1 ligands at extracellular milieu shifting homeostatic cartilage matrix turnover to a catabolic state [19]. The impairment of LRP1 is due to an increase in the proteolytic shedding of the ectodomain of the receptor from the cell surface and we have previously identified matrix metalloproteinase 14 (MMP14) and a disintegrin and metalloproteinase 17 (ADAM17) as the major LRP1 sheddases in human OA cartilage. Importantly, inhibition of LRP1 shedding recovered the endocytic capacity of the cells and reduced ECM degradation in OA cartilage [19]. Dysregulated LRP1 shedding is also observed in various pathological conditions such as rheumatoid arthritis, systemic lupus erythematosus [27], and in cancer [28, 29].

To date, more than 80 LRP1 ligands have been reported in the literature [4, 30] and the list is still growing. However, very limited information is available on how LRP1 ligand profiles, i.e., its ligandome, differ in specific tissues including cartilage. This is partly due to that tightly regulated LRP1 ligands are rarely detectable in the tissue unless LRP1-mediated endocytosis is blocked [22, 23, 26, 31]. Uncovering the depth of this “hidden” ligandome is essential to identify novel molecular interactions, and possibly therapeutic opportunities. In this study, we took a systematic top-down approach to uncover the LRP1 ligandome in healthy cartilage. We have identified 38 novel LRP1 interaction partners whose excessive activities exerts detrimental consequences for chondrocytes and cartilage tissue integrity.

## Results

### Inhibition of LRP1-mediated endocytosis induces chondrocyte cell death

Inhibition of LRP1-mediated endocytosis is essential to enable the identification of tightly regulated LRP1 ligands. We thus decided to block LRP1-mediated endocytosis in human chondrocytes using either 10 nM soluble form of full-length LRP1 (sLRP1) or 500 nM receptor-associated protein (RAP). sLRP1 binds to LRP1 ligands and acts as a soluble decoy receptor competing with the endogenous LRP1, whereas RAP antagonises ligand binding by competitively occupying the ligand binding region of LRP1 (**Fig 1A**). Initially, we evaluated the effect of sLRP1 and RAP on chondrocytes morphology and viability. We found that 72-h incubation with either the inhibitor or soluble receptor resulted in altered chondrocyte morphology and a significant number of the cells detaching from the plate surface (**Fig 1B**). The MTS cell metabolic assay showed that compared to 0-h, approx. 53.7% and 57.7% of the cells were dead after 72-h incubation with sLRP1 or RAP, respectively (**Fig 1C**). The time-dependent effect of sLRP1 and RAP on the cell death (**Fig 1D**) indicates that the accumulation of LRP1 ligands gradually leads to chondrocyte cell death.

**Figure 1.**
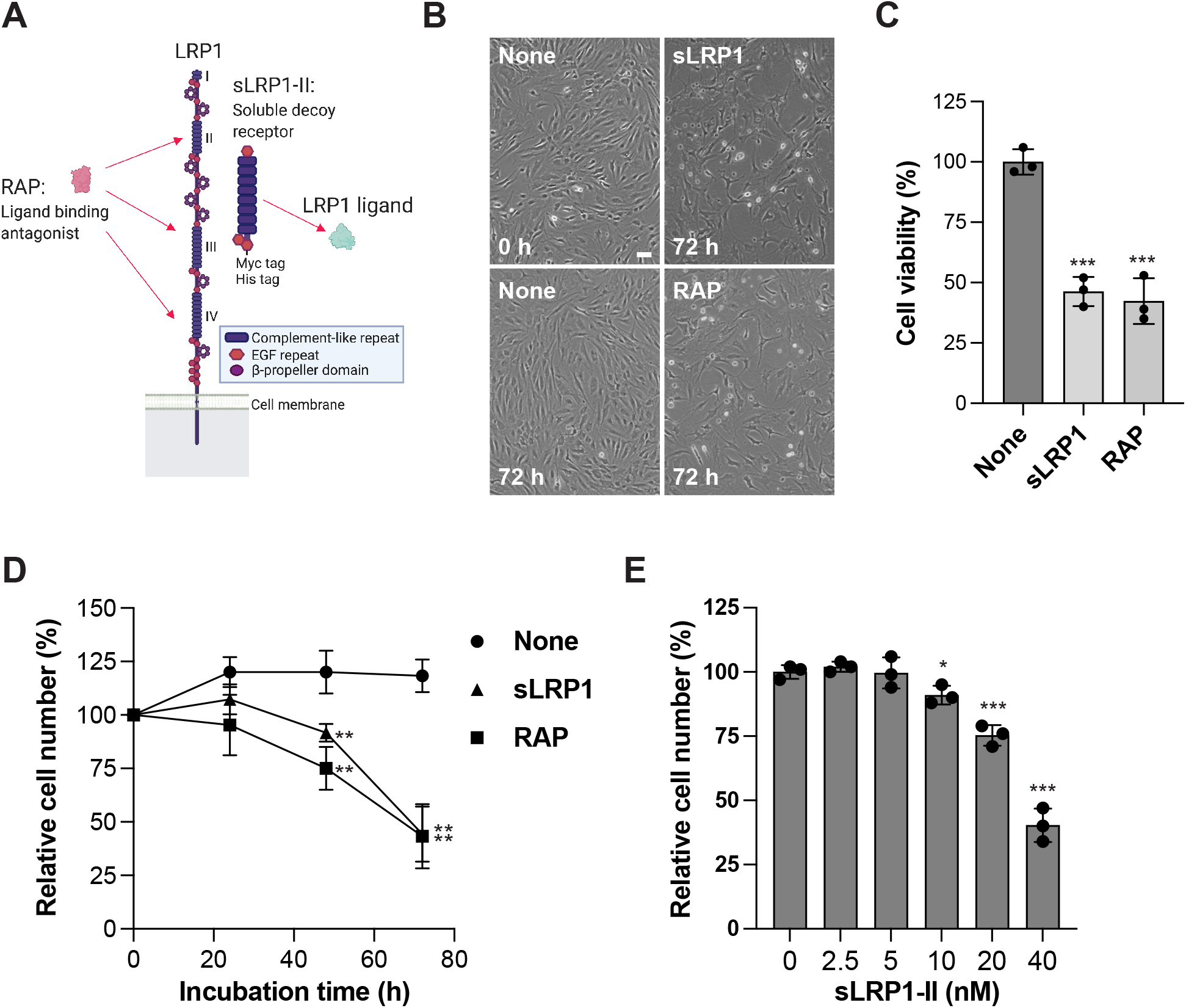
Inhibition of LRP1-mediated endocytosis induces chondrocyte cell death. *A*, Schematic representation for membrane-bound LRP1, receptor-associated protein (RAP), and the soluble form of LRP1 ligand binding cluster II (sLRP1-II). sLRP1-II containing a C-terminal 6xHis/c-Myc tag and c-Myc antibody. RAP antagonises LRP1 ligands by binding to the ligand binding clusters II-IV whereas sLRP1 binds to LRP1 ligands and acts as a soluble decoy receptor. *B-D*, Human chondrocytes from three different donors were cultured with serum-free DMEM in the absence or presence of 10 nM full-length sLRP1 or 500 nM RAP for 0-72 h. Cell morphology was observed and imaged after 72-h incubation. Scale bar, 30 μm (*B*). Cell viability was measured using the MTS assay and the cell viability at each condition was expressed as a % of the cell number at 0 h (*C*). Data are expressed as the mean ± *SD*. Cell number was estimated using hemocytometer and analysed as in (*D*). *E*, Human chondrocytes from three different donors were cultured with serum-free DMEM in the absence or presence of 0-40 nM sLRP1-II for 24 h. Cell number was counted and expressed as in *D*. Circles represent individual cartilage donors and bars show the mean ± *SD*. *, *p* < 0.05, **, *p* < 0.01, ***, *p* < 0.001 by 2-tailed Student’s t test.

The ligand binding regions in LRP1 lie in four clusters (clusters I-IV), containing 2 to 11 individual ligand binding cysteine-rich complement-like repeats. Clusters II and IV are considered to be responsible for most ligand binding [32–34] and our previous studies have demonstrated the importance of cluster II for the binding of a disintegrin and metalloproteinase with thrombospondin motifs (ADAMTS)4, ADAMTS5 [21], MMP13 [23] and TIMP3 [35]. Thus, we next tested the effect of the soluble form of LRP1 ligand-binding cluster II (sLRP1-II) on chondrocyte cell death. After 24-h incubation with 40 nM sLRP1-II, approx. 60% of the cells were dead compared to 0-h (**Fig 1E**). We further examined the dose-dependent effect of sLRP1-II and found that 24-h incubation with 20 nM and 10 nM sLRP1-II resulted in approx. 25% and 9% of cell death, respectively, whereas 5 nM and 2.5 nM sLRP1-II showed negligible effect (**Fig 1E**). The highest concentration of sLRP1-II that does not induce the cell death up to 24-h incubation (5 nM) was thus employed for LRP1 ligand identification.

### Alteration of the entire chondrocyte secretome by inhibition of LRP1-mediated endocytosis

We performed a complete chondrocyte secretome analysis to determine the effect of inhibition of LRP1-mediated endocytosis and identify LRP1 ligands whose excess activity exerts detrimental consequences for chondrocytes. This analysis identified 635 proteins including 197 secreted, 415 intracellular and 23 transmembrane molecules according to the Uniprot annotation. Twenty-four hour-treatment of 5 nM sLRP1-II resulted in significant alteration (*p*-value <0.05) of 417 molecules with an increase in 34 secreted, 253 intracellular, and 10 transmembrane proteins and a decrease in 104 secreted, 9 intracellular and 7 transmembrane proteins (**Fig 2A-C** and **Suppl Table I - III**). The secreted proteins significantly (>2.5-fold difference and p-value <0.05) increased or decreased in the presence of sLRP1-II were listed on **Table I** or **Table II**, respectively. Among the 34 secreted proteins increased by the sLRP1-II treatment, only 10 molecules have previously been reported as LRP1 ligands (**Fig 2C**). Unexpectedly, a total of 21 previously reported LRP1 ligands were decreased (12) or unaltered (9) by the sLRP1-II treatment (**Fig 2D**). These results suggest that the identified chondrocyte LRP1 ligandome consists of mostly novel ligands and that the 21 previously reported LRP1 ligands are not regulated by LRP1-mediated endocytosis at least in human chondrocytes.

**Figure 2.**
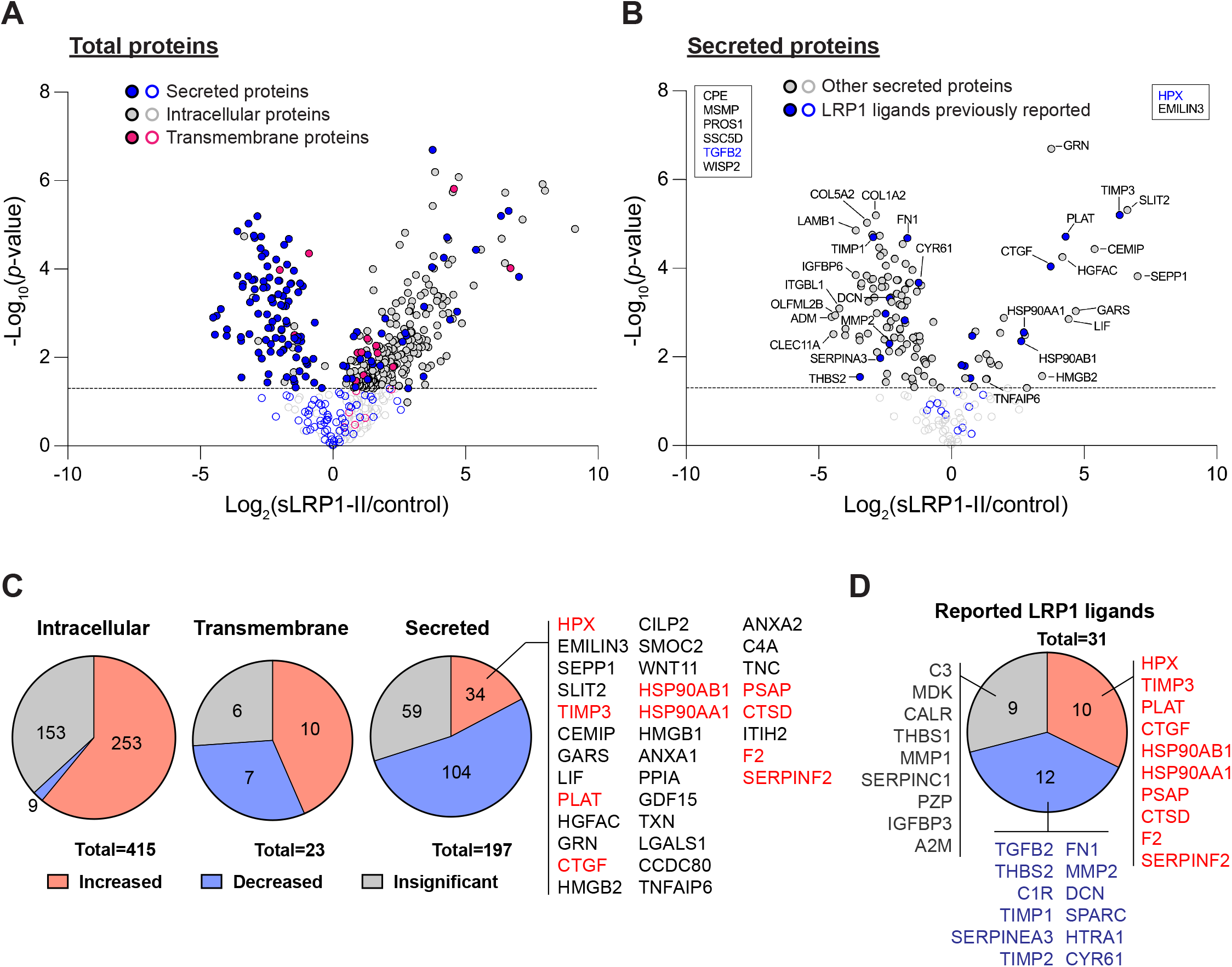
Alteration of the entire chondrocyte secretome by inhibition of LRP1-mediated endocytosis. Human chondrocytes from four different donors were incubated with serum-free DMEM in the absence or presence of 5 nM sLRP1-II for 24 h. The conditioned medium was collected and then subjected to mass spectrometry-based protein identification. *A*, Volcano plot showing the −log_10_ of *p*-values *versus* the log2 of protein ratio for all proteins identified in the medium of chondrocytes in the presence and absence of sLRP1-II. The proteins only found in each condition were not shown. Blue, grey and pink circles represent secreted, intracellular and transmembrane proteins, respectively. *B*, Volcano plot showing the −log_10_ of *p*-values *versus* the log2 of protein ratio for only secreted proteins. Left top panel shows proteins found in the medium of chondrocytes only when sLRP1-II was not added. Right top panel shows proteins found in the medium of chondrocytes only when sLRP1-II was added. Previously reported LRP1 ligands in these panels are highlighted with blue. Blue and grey circles represent previously reported LRP1 ligands and non-ligands, respectively. Closed and open circles represents significantly (p-value <0.05) and non-significantly regulated molecules, respectively. *C*, Pie chart showing number of secreted, intracellular and transmembrane proteins that were significantly increased or decreased, or not significantly changed in the presence and absence of sLRP1-II. Gene name for the significantly increased secreted proteins were listed and previously reported LRP1 ligands highlighted with red. *D*, Pie chart showing previously reported LRP1 ligands with their gene name that were significantly increased or decreased, or not significantly changed in the presence and absence of sLRP1-II.

**Table I:**
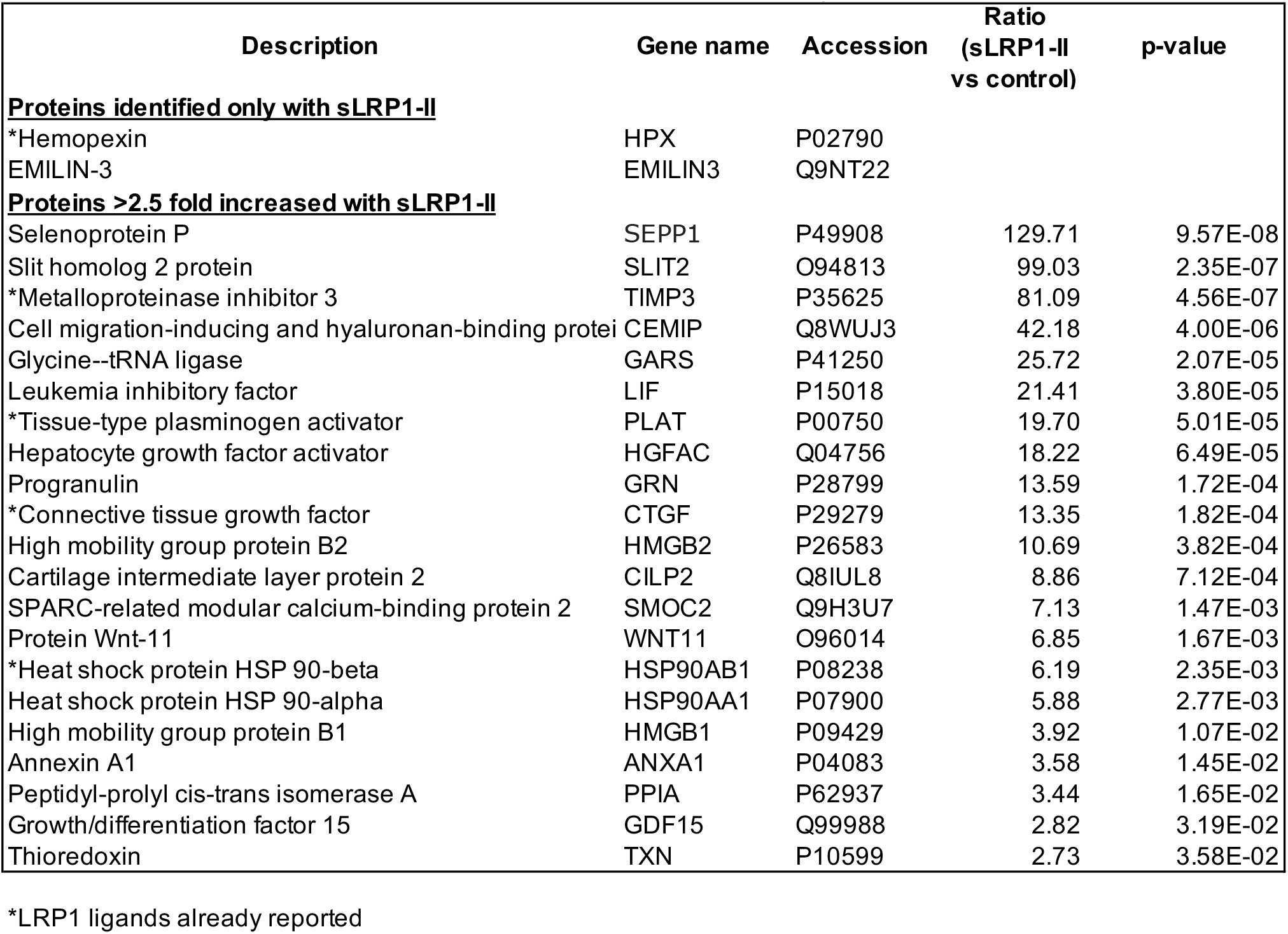
Secreted molecules increased in medium of human chondrocytes after 24-h incubation with sLRP1-II.

| Description | Gene name | Accession | Ratio<br>(sLRP1-II<br>vs control) | p-value |
| --- | --- | --- | --- | --- |
| **Proteins identified only with sLRP1-II** |  |  |  |  |
| *Hemopexin | HPX | P02790 |  |  |
| EMILIN-3 | EMILIN3 | Q9NT22 |  |  |
| **Proteins >2.5 fold increased with sLRP1-II** |  |  |  |  |
| Selenoprotein P | SEPP1 | P49908 | 129.71 | 9.57E-08 |
| Slit homolog 2 protein | SLIT2 | O94813 | 99.03 | 2.35E-07 |
| *Metalloproteinase inhibitor 3 | TIMP3 | P35625 | 81.09 | 4.56E-07 |
| Cell migration-inducing and hyaluronan-binding protein | CEMIP | Q8WUJ3 | 42.18 | 4.00E-06 |
| Glycine--tRNA ligase | GARS | P41250 | 25.72 | 2.07E-05 |
| Leukemia inhibitory factor | LIF | P15018 | 21.41 | 3.80E-05 |
| *Tissue-type plasminogen activator | PLAT | P00750 | 19.70 | 5.01E-05 |
| Hepatocyte growth factor activator | HGFAC | Q04756 | 18.22 | 6.49E-05 |
| Progranulin | GRN | P28799 | 13.59 | 1.72E-04 |
| *Connective tissue growth factor | CTGF | P29279 | 13.35 | 1.82E-04 |
| High mobility group protein B2 | HMGB2 | P26583 | 10.69 | 3.82E-04 |
| Cartilage intermediate layer protein 2 | CILP2 | Q8IUL8 | 8.86 | 7.12E-04 |
| SPARC-related modular calcium-binding protein 2 | SMOC2 | Q9H3U7 | 7.13 | 1.47E-03 |
| Protein Wnt-11 | WNT11 | O96014 | 6.85 | 1.67E-03 |
| *Heat shock protein HSP 90-beta | HSP90AB1 | P08238 | 6.19 | 2.35E-03 |
| Heat shock protein HSP 90-alpha | HSP90AA1 | P07900 | 5.88 | 2.77E-03 |
| High mobility group protein B1 | HMGB1 | P09429 | 3.92 | 1.07E-02 |
| Annexin A1 | ANXA1 | P04083 | 3.58 | 1.45E-02 |
| Peptidyl-prolyl cis-trans isomerase A | PPIA | P62937 | 3.44 | 1.65E-02 |
| Growth/differentiation factor 15 | GDF15 | Q99988 | 2.82 | 3.19E-02 |
| Thioredoxin | TXN | P10599 | 2.73 | 3.58E-02 |

\*LRP1 ligands already reported

**Table II:**
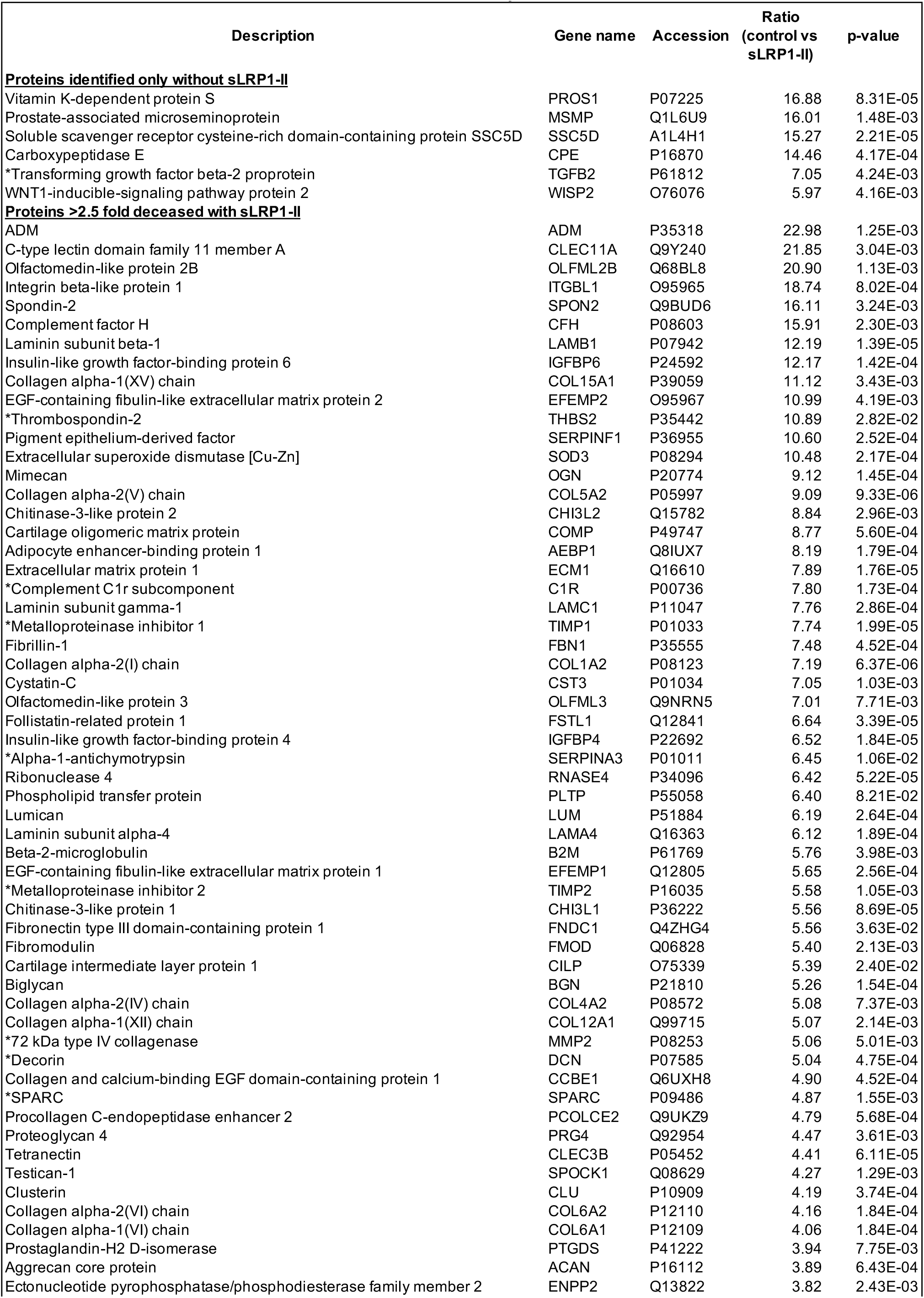
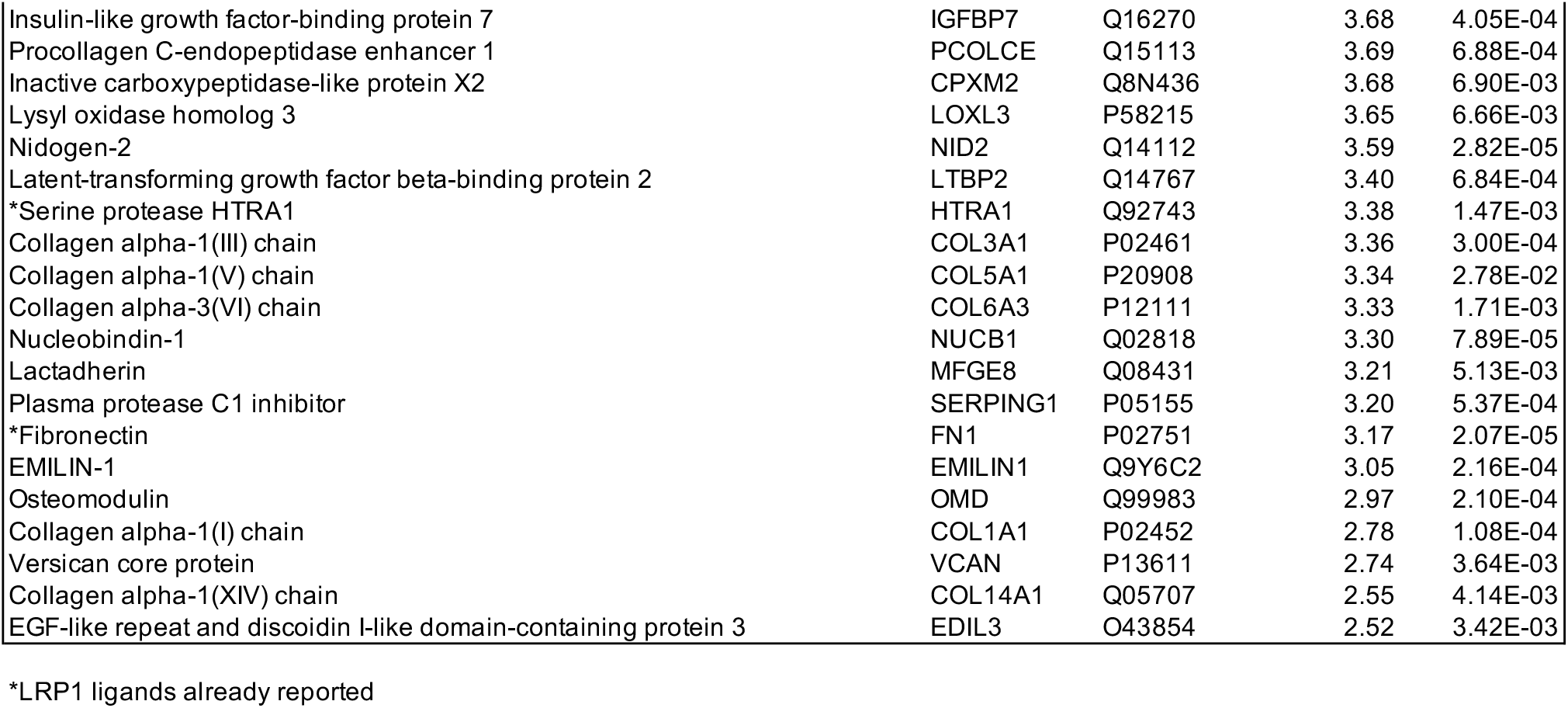
Secreted molecules decreased in medium of human chondrocytes after 24-h incubation with sLRP1-II.

To validate the chondrocyte secretome analysis, hyaluronan-binding protein (CEMIP, also known as KIAA1199 and HYBID), slit homolog 2 protein (SLIT2), high-mobility group protein (HMG)B2 and tumour necrosis factor (TNF)-inducible gene 6 protein (TSG6, TNFAIP6 in the plot) were further investigated by Western blot analysis. Endogenously produced CEMIP and HMGB2 were rarely detectable in the conditioned medium of human chondrocytes after 24-h incubation without sLRP1-II but markedly increased in the presence of 5 nM sLRP1-II (**Suppl Fig 1A and B**). The sLRP1-II treatment increased the levels of SLIT2 and TSG6 in the medium of chondrocytes approx. 2.5- and 2.8-fold, respectively, compared to the control (**Suppl Fig 1C and D**).

### Inhibition of LRP1-mediated endocytosis induces transcriptional modulations

It has been reported that RAP increases mRNA levels of pro-inflammatory mediators such as TNFα, interleukin-6 and C–C motif chemokine ligand 2 in macrophages [36]. We next investigated effect of inhibition of LRP1-mediated endocytosis on transcriptional modulation of several LRP1 ligands and a non-ligand. Our recent [24] study identified MMP1, a collagenase highly elevated in knee joint and synovial fluid from patients with arthritis [37, 38], as an LRP1 ligand. We found that endogenously produced MMP1 was not detectable in the medium of chondrocytes in the absence of RAP, whereas it was increased after 24 h-incubation with 200 nM RAP (**Fig 3A**). Similar results were obtained for MMP3 (**Fig 3B**), which is not a LRP1 ligand but relevant to arthritis pathogenesis [37, 38]. Notably, the quantitative PCR analysis revealed that mRNA levels of MMP1 and MMP3 in human chondrocytes are increased time-dependently in the presence of 200 nM RAP or 10 nM sLRP1 (**Fig 3C and D**). In contrast, denatured sLRP1 did not affect their mRNA expression, suggesting that functional sLRP1 is required for upregulation of mRNA levels of these proteinases. Similar results were obtained for other LRP1 ligands, MMP13 and ADAMTS4 (**Fig 3E and F**), whereas mRNA levels of ADAMTS5 or TIMP3 were not affected by inhibition of LRP1-mediated endocytosis (**Fig 3G and H**). These studies suggest that excessive activity of LRP1 ligands alters transcription of several molecules. In addition, a fraction of the molecules increased upon sLRP1-II treatment in our secretome analysis (**Fig 2**) might not be LRP1 ligands but due to transcriptional upregulation.

**Figure 3.**
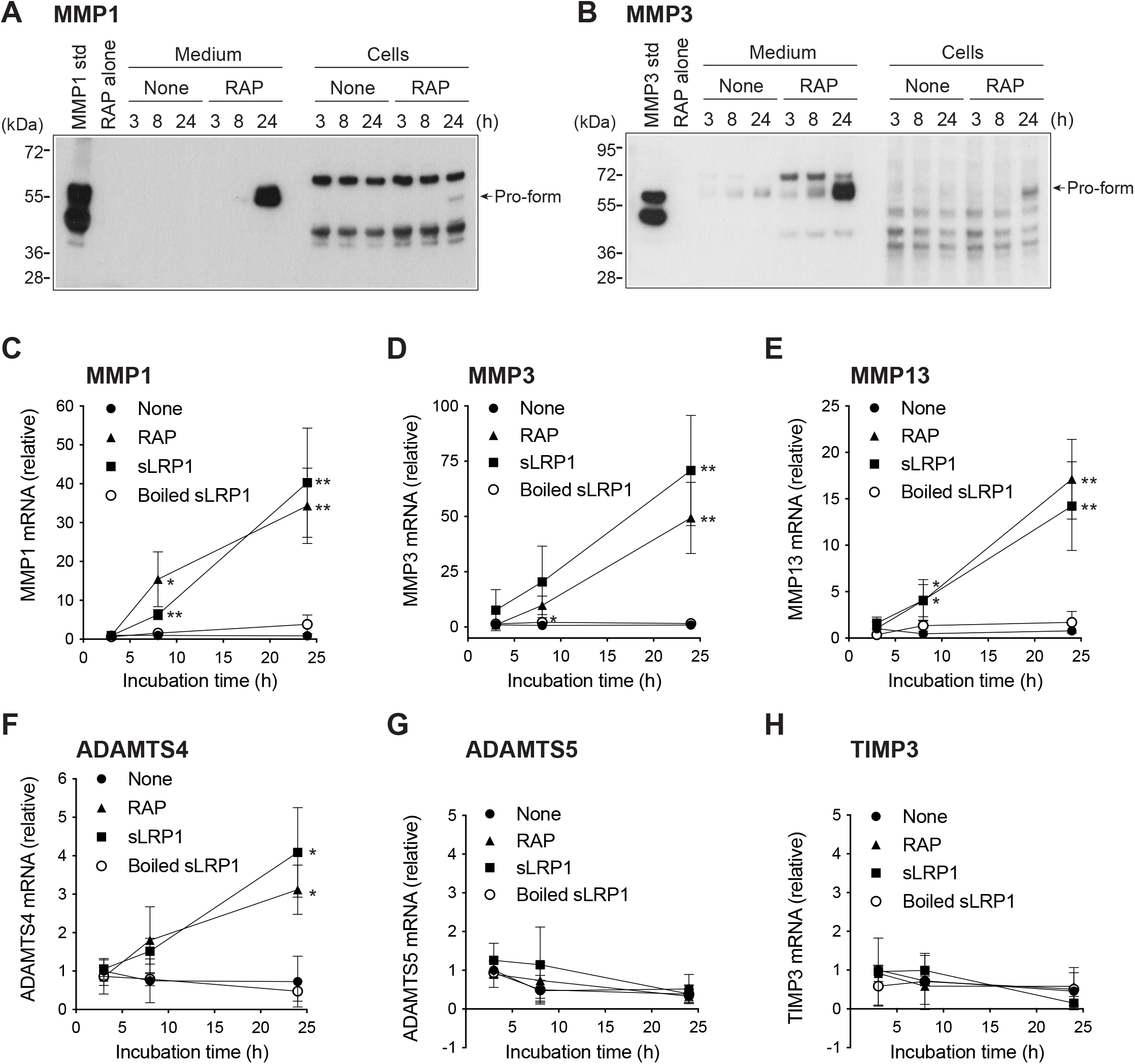
Inhibition of LRP1-mediated endocytosis induces transcriptional modulation. Human normal chondrocytes from three different donors were incubated with serum-free DMEM containing CT1746 (100 μM) and the protease inhibitor cocktail (1/500) in the absence or presence of 200 nM RAP (*A and B*), 10 nM sLRP1 or boiled and denatured sLRP1 (*C-H*) for 0-24 h. Representative Western blotting analysis of MMP1 (*A*) and MMP3 (*B*) in the media and cell lysate detected using the anti-MMP1 and anti-MMP3 antibodies. TaqMan real-time PCR showing relative levels of mRNA for MMP1 (*C*), MMP3 (*D*), MMP13 (*E*), ADAMTS4 (*F*), ADAMTS5 (*G*) and TIMP3 (*H*) in human chondrocytes. Points represent the means ± *SD*. *, *p* < 0.05, **, *p* < 0.01, by 2-tailed Student’s t test. *Std*: 10 nM purified protein for pro- and activated forms of MMP1 and MMP3.

### The chondrocyte LRP1 ligandome comprises a variety of novel ligands

To enrich and identify molecules that directly interact with LRP1, we have employed co-immunoprecipitation assay for sLRP1-II containing a C-terminal 6xHis/c-Myc tag and c-Myc antibody. The isolation of LRP1 ligands using this method was first validated using ADAMTS5 (**Suppl Fig 2**). Purified ADAMTS5 (5 nM) was mixed with 5 nM sLRP1-II and anti-c-Myc antibody-coupled paramagnetic beads, recognising sLRP1-II. Complexes of sLRP1-II, ADAMTS5 and the magnetic beads were then isolated using a magnetic separator. Most of ADAMTS5 and sLRP1-II were detected in the bound fraction (**Suppl Fig 2A**). As expected, RAP competitively inhibited ADAMTS5 binding to sLRP1-II. Since LRP1 ligands dissociate from LRP1 in early endosome due to acidic conditions, we have tested whether acidic conditions (pH 3.0) can dissociate molecules bound to sLRP1-II/anti-c-Myc antibody complexes. As expected, ADAMTS5 were eluted under acidic conditions as effectively as 1% sodium dodecyl sulphate (SDS) or 100 nM RAP (**Suppl Fig 2B**). To identify LRP1 ligands produced by human chondrocytes, cells were incubated without (control) or with 5 nM sLRP1-II for 24 h and the molecules bound to sLRP1-II were isolated as described above (**Fig 4A**). The molecules isolated by the same procedure without sLRP1-II were considered as a negative control and used to estimate a fold-difference for the ligand identification. Western blot analysis of the isolated proteins using anti-His tag antibodies has confirmed the isolation of sLRP1-II added to the culture (**Suppl Fig 3**).

**Figure 4.**
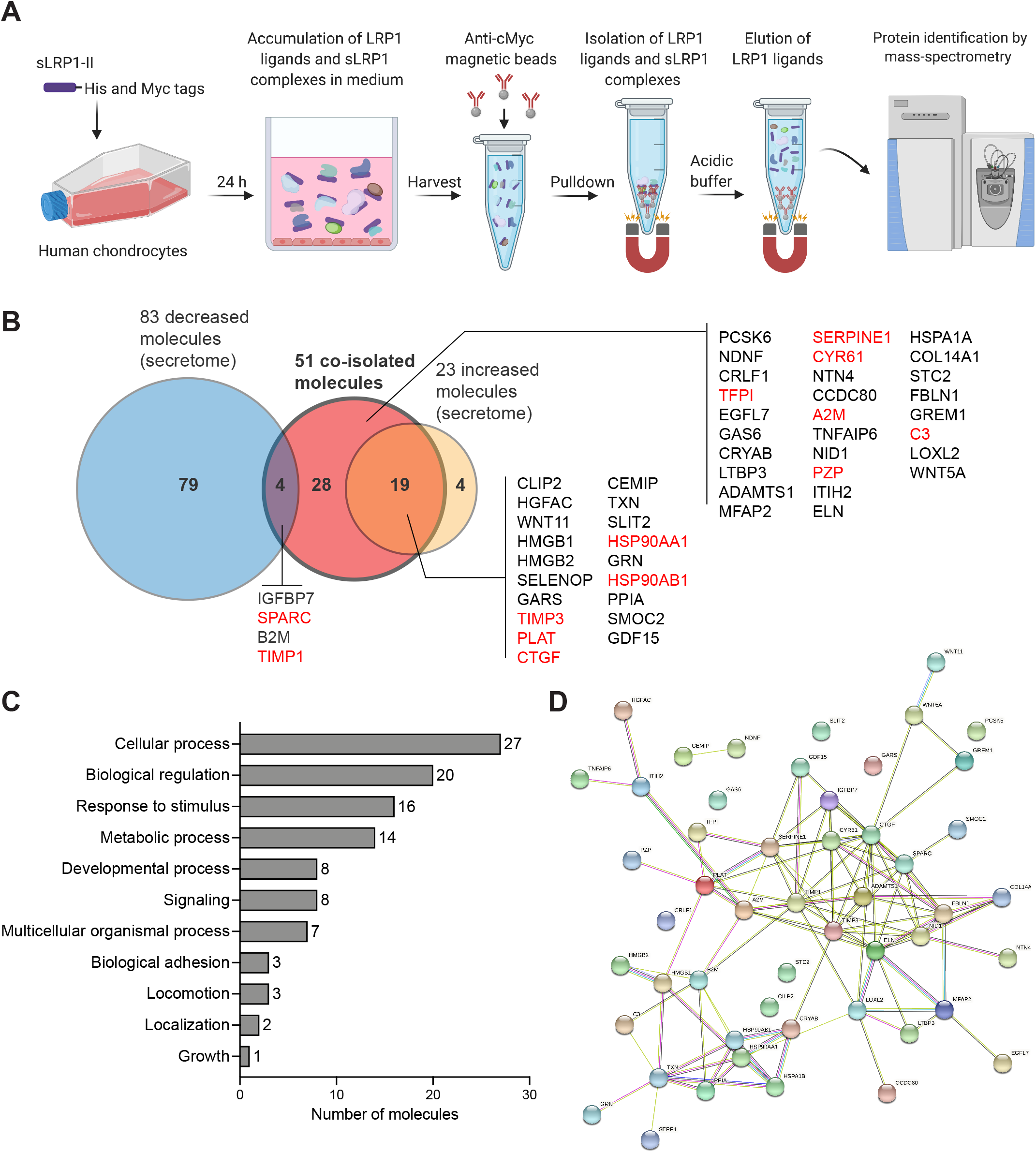
Chondrocyte LRP1 interactome consists of a variety of novel ligand candidates. *A*, Workflow of isolation and identification of LRP1 ligands in human chondrocytes from four different donors. *B*, Venn diagrams showing secreted proteins that were co-isolated with sLRP1-II (red), secreted proteins increased (orange) or decreased (blue) in medium of chondrocytes treated with sLRP1-II (**Fig 2**). *C*, The PANTHER gene ontology analysis for biological processes of 51 secreted proteins co-isolated with sLRP1-II from the medium of human chondrocytes. *D*, The STRING functional and physical interaction map of 51 secreted proteins co-isolated with sLRP1-II from the medium of human chondrocytes.

Mass spectrometry analysis identified 276 molecules co-immunoprecipitated with sLRP1-II from the medium of human chondrocytes. These molecules include 51 secreted proteins, whereas the remainder of the molecules are either intracellular or transmembrane proteins according to the Uniprot annotation (**Suppl Table IV**), supporting a notion that LRP1 plays a role in the clearance of cellular debris [30]. Among the 51 secreted proteins identified, 13 molecules were previously reported LRP1 ligands, while 38 molecules were novel LRP1 ligand candidates (**Fig 4B and Table III**). These 38 ligand candidates included 23 proteins that were significantly increased (19) or decreased (4) (>2.5-fold difference and p-value <0.05) by sLRP1-II treatment in the secretome analysis, respectively. The Protein ANalysis THrough Evolutionary Relationships (PANTHER) [39] gene ontology analysis on biological processes of these molecules that interact with LRP1 showed that 27, 20, 16 and 14 of them are involved in “Cellular processes”, “Biological regulation”, “Response to stimulus” and “Metabolic processes”, respectively (**Fig 4C**). The Search Tool for Retrieval of Interacting Genes/Proteins (STRING) analysis [40] of protein-protein interactions further revealed functional and physical interactions of these molecules illustrated in (**Fig 4D**).

**Table III:**
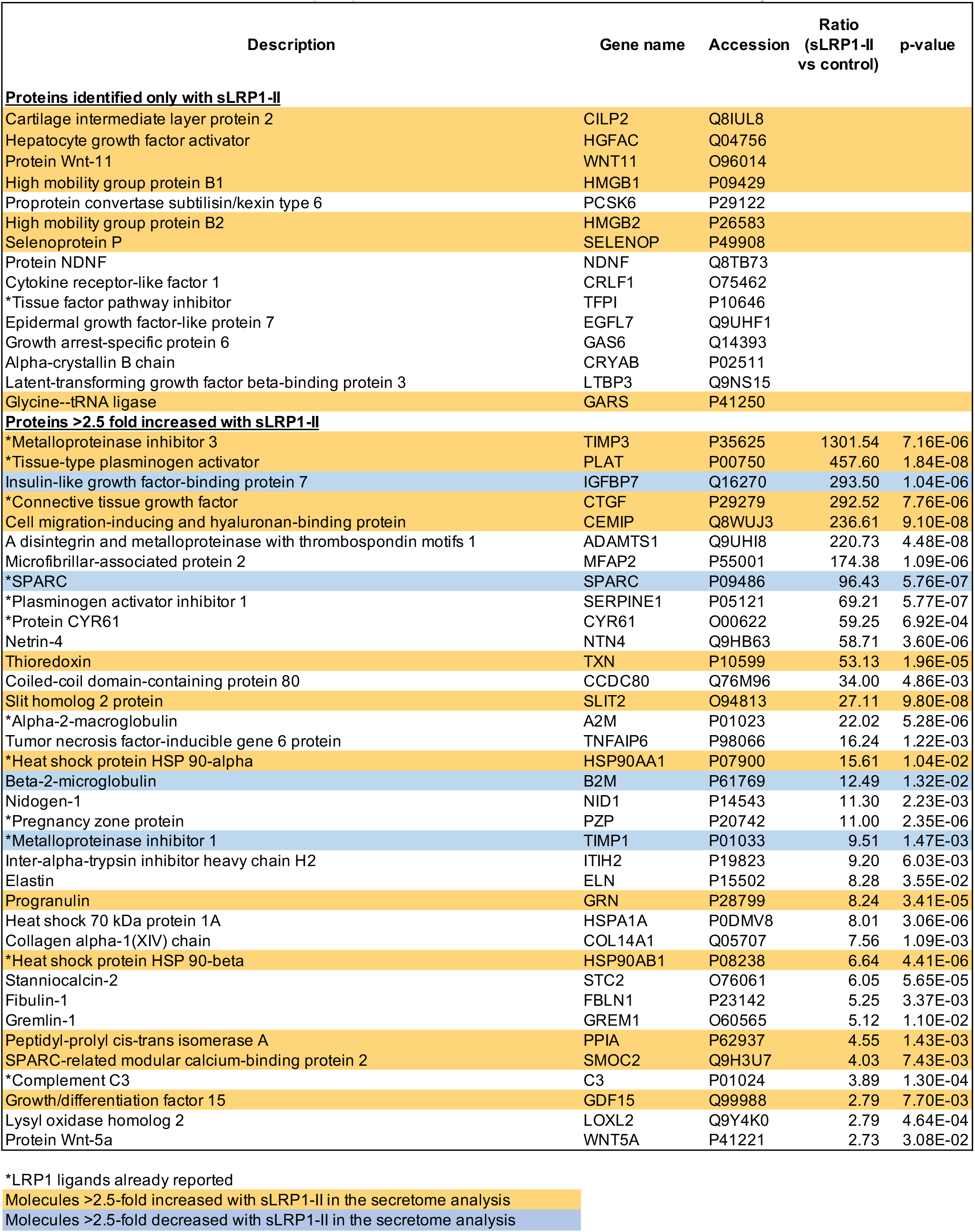
Secreted molecules co-immunoprecipitated with sLRP1-II from medium of human chondrocytes.

| Description | Gene name | Accession | Ratio<br>(sLRP1-II<br>vs control) | p-value |
| --- | --- | --- | --- | --- |
| **Proteins identified only with sLRP1-II** |  |  |  |  |
| Cartilage intermediate layer protein 2 | CILP2 | Q8IUL8 |  |  |
| Hepatocyte growth factor activator | HGFAC | Q04756 |  |  |
| Protein Wnt-11 | WNT11 | O96014 |  |  |
| High mobility group protein B1 | HMGB1 | P09429 |  |  |
| Proprotein convertase subtilisin/kexin type 6 | PCSK6 | P29122 |  |  |
| High mobility group protein B2 | HMGB2 | P26583 |  |  |
| Selenoprotein P | SELENOP | P49908 |  |  |
| Protein NDNF | NDNF | Q8TB73 |  |  |
| Cytokine receptor-like factor 1 | CRLF1 | O75462 |  |  |
| *Tissue factor pathway inhibitor | TFPI | P10646 |  |  |
| Epidermal growth factor-like protein 7 | EGFL7 | Q9UHF1 |  |  |
| Growth arrest-specific protein 6 | GAS6 | Q14393 |  |  |
| Alpha-crystallin B chain | CRYAB | P02511 |  |  |
| Latent-transforming growth factor beta-binding protein 3 | LTBP3 | Q9NS15 |  |  |
| Glycine--tRNA ligase | GARS | P41250 |  |  |
| **Proteins >2.5 fold increased with sLRP1-II** |  |  |  |  |
| *Metalloproteinase inhibitor 3 | TIMP3 | P35625 | 1301.54 | 7.16E-06 |
| *Tissue-type plasminogen activator | PLAT | P00750 | 457.60 | 1.84E-08 |
| Insulin-like growth factor-binding protein 7 | IGFBP7 | Q16270 | 293.50 | 1.04E-06 |
| *Connective tissue growth factor | CTGF | P29279 | 292.52 | 7.76E-06 |
| Cell migration-inducing and hyaluronan-binding protein | CEMIP | Q8WUJ3 | 236.61 | 9.10E-08 |
| A disintegrin and metalloproteinase with thrombospondin motifs 1 | ADAMTS1 | Q9UHI8 | 220.73 | 4.48E-08 |
| Microfibrillar-associated protein 2 | MFAP2 | P55001 | 174.38 | 1.09E-06 |
| *SPARC | SPARC | P09486 | 96.43 | 5.76E-07 |
| *Plasminogen activator inhibitor 1 | SERPINE1 | P05121 | 69.21 | 5.77E-07 |
| *Protein CYR61 | CYR61 | O00622 | 59.25 | 6.92E-04 |
| Netrin-4 | NTN4 | Q9HB63 | 58.71 | 3.60E-06 |
| Thioredoxin | TXN | P10599 | 53.13 | 1.96E-05 |
| Coiled-coil domain-containing protein 80 | CCDC80 | Q76M96 | 34.00 | 4.86E-03 |
| Slit homolog 2 protein | SLIT2 | O94813 | 27.11 | 9.80E-08 |
| *Alpha-2-macroglobulin | A2M | P01023 | 22.02 | 5.28E-06 |
| Tumor necrosis factor-inducible gene 6 protein | TNFAIP6 | P98066 | 16.24 | 1.22E-03 |
| *Heat shock protein HSP 90-alpha | HSP90AA1 | P07900 | 15.61 | 1.04E-02 |
| Beta-2-microglobulin | B2M | P61769 | 12.49 | 1.32E-02 |
| Nidogen-1 | NID1 | P14543 | 11.30 | 2.23E-03 |
| *Pregnancy zone protein | PZP | P20742 | 11.00 | 2.35E-06 |
| *Metalloproteinase inhibitor 1 | TIMP1 | P01033 | 9.51 | 1.47E-03 |
| Inter-alpha-trypsin inhibitor heavy chain H2 | ITIH2 | P19823 | 9.20 | 6.03E-03 |
| Elastin | ELN | P15502 | 8.28 | 3.55E-02 |
| Progranulin | GRN | P28799 | 8.24 | 3.41E-05 |
| Heat shock 70 kDa protein 1A | HSPA1A | P0DMV8 | 8.01 | 3.06E-06 |
| Collagen alpha-1(XIV) chain | COL14A1 | Q05707 | 7.56 | 1.09E-03 |
| *Heat shock protein HSP 90-beta | HSP90AB1 | P08238 | 6.64 | 4.41E-06 |
| Stanniocalcin-2 | STC2 | O76061 | 6.05 | 5.65E-05 |
| Fibulin-1 | FBLN1 | P23142 | 5.25 | 3.37E-03 |
| Gremlin-1 | GREM1 | O60565 | 5.12 | 1.10E-02 |
| Peptidyl-prolyl cis-trans isomerase A | PPIA | P62937 | 4.55 | 1.43E-03 |
| SPARC-related modular calcium-binding protein 2 | SMOC2 | Q9H3U7 | 4.03 | 7.43E-03 |
| *Complement C3 | C3 | P01024 | 3.89 | 1.30E-04 |
| Growth/differentiation factor 15 | GDF15 | Q99988 | 2.79 | 7.70E-03 |
| Lysyl oxidase homolog 2 | LOXL2 | Q9Y4K0 | 2.79 | 4.64E-04 |
| Protein Wnt-5a | WNT5A | P41221 | 2.73 | 3.08E-02 |

\*LRP1 ligands already reported
Molecules >2.5-fold increased with sLRP1-II in the secretome analysis
Molecules >2.5-fold decreased with sLRP1-II in the secretome analysis

### Newly identified LRP1 ligands exhibit a wide range of affinities

Based on the mass-spectrometry scores, reported pathophysiological functions and a feasibility to obtain purified proteins, we selected 11 novel ligand candidates for further characterization: hepatocyte growth factor activator (HGFA, gene name: HGFAC), inflammatory mediators high-mobility group protein (HMG)B1 and HMGB2, CEMIP, slit homolog 2 protein (SLIT2), secreted growth factor progranulin (gene name: GRN), proteoglycanase ADAMTS1, which was increased in medium of HEK293 cells treated with RAP in our previous study [41], tumour necrosis factor (TNF)-inducible gene 6 protein (TSG6, gene name: TNFAIP6), ECM glycoprotein fibulin1C (gene name: FBLN1), insulin-like growth factor-binding protein (IGFBP)7, secreted protein acidic and rich in cysteine (SPARC, also known as osteonectin), which is previously identified as one of the proteins co-isolated with sLRP1 binding clusters II and IV [30] but no evidence of direct LRP1 binding provided, and leukemia inhibitory factor (LIF). To confirm their direct binding and estimate their affinity for LRP1, solid-phase binding assay were performed using purified ligand candidates and full-length sLRP1. We found that HMGB1 (6.4 nM), HMGB2 (3.9 nM), CEMIP (2.6 nM), SLIT2 (2.0 nM), ADAMTS1 (20.2 nM), TSG6 (11.0 nM), IGFBP7 (11.0 nM) and SPARC (41.0 nM) directly bind to immobilised LRP1 with high affinity (apparent binding constant (*K_D,app_*) indicated in parentheses)(**Fig 5B-E, GH and JK**). HGFA (>500 nM) and LIF (>200 nM) also directly bind to immobilised LRP1 but their affinities for LRP1 were much weaker than the high affinity ligands mentioned above (**Fig 5A and L**). In contrast, LRP1 binding to progranulin or fibulin-1C was not observed at all (**Fig 5F and I**). These results indicate that the molecules co-isolated with sLRP1-II include not only direct LRP1 binders but also non LRP1 binders that may interact with LRP1 indirectly, possibly *via* true ligands.

**Figure 5.**
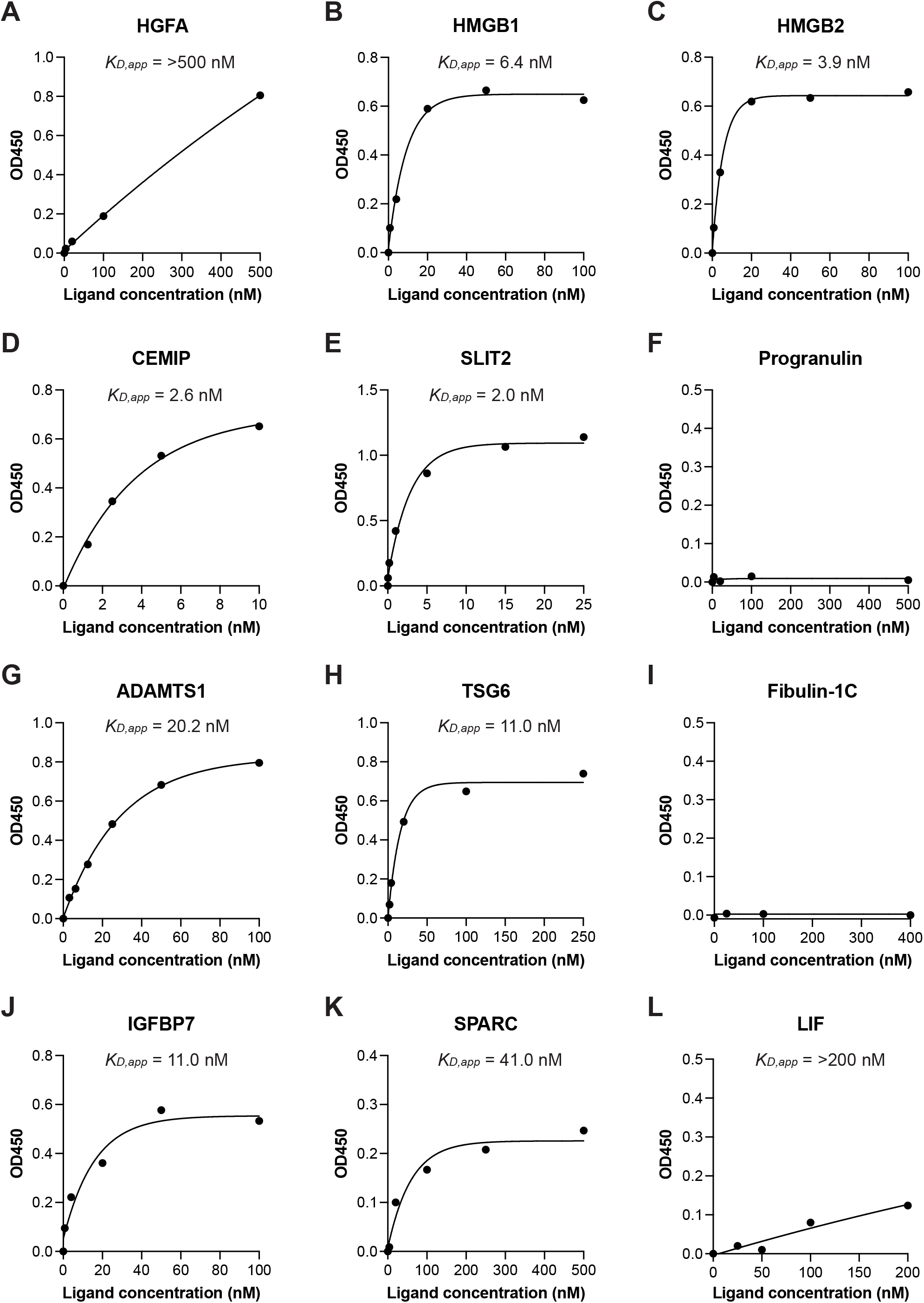
A diverse range of affinities of newly identified ligands for LRP1. Full-length sLRP1 was coated onto microtiter plates and the binding of various concentrations of HGFA (*A*), HMGB1 (*B*), HMGB2 (*C*), CEMIP (*D*), SLIT2 (*E*), progranulin (*F*), ADAMTS1 (*G*), TSG6 (*H*), fibulin-1C (*I*), IGFBP7 (*J*), SPARC (*K*) and LIF (*L*) was measured using anti-FLAG M2 or anti-His tag antibody and a horseradish peroxidase-conjugate secondary antibody as described under “Experimental procedures”. Mean values of technical duplicates for none-coat, LRP1-coat and after normalization by subtracting the none-coat values from the LRP1-coat values were shown as circles. Extrapolated *K_D,app_* values were estimated based on one-phase decay nonlinear fit analysis (black lines).

### LRP1-mediated endocytic clearance of novel LRP1 ligands

Newly identified proteins that directly bind to LRP1 were further tested for their endocytic clearance by cells *via* LRP1. Recombinant HGFA, HMGB2, CEMIP, SLIT2, ADAMTS1, TSG6, IGFBP7 and SPARC (each 10 nM) were incubated with wild-type (WT) mouse embryonic fibroblasts (MEFs) or LRP1-deficient MEFs [42] for 8 h and these molecules in conditioned medium and cell lysate were detected by Western blot analysis using anti-FLAG or His tag antibody. The amounts of exogenously added HMGB2, CEMIP, SLIT2, ADAMTS1, TSG6 and IGFBP7 in the conditioned medium were reduced in WT MEFs by 59%, 92%, 65%, 62%, 80% and 45%, respectively, compared to LRP1-deficient MEFs, whereas no changes were observed for HGFA (**Fig 6A-G**). SPARC was also reduced in WT MEFs by 25% compared to LRP1-deficient cells but the differences appeared to be not statistically significant (**Fig 6H**). Exogenously added SLIT2, ADAMTS1, TSG6 and IGFBP7 were also detected in the cell lysate and the slightly lower amounts of SLIT2, TSG6 and IGFBP7 were detected in WT compared to LRP1-deficient MEFs (**Fig 6D-G**). These results indicate that if the same concentration of LRP1 ligands is present in the extracellular milieu, the ligands with higher affinity are taken up *via* LRP1-mediated endocytosis more rapidly than low affinity binders (**Fig 6I and Table IV**).

**Figure 6.**
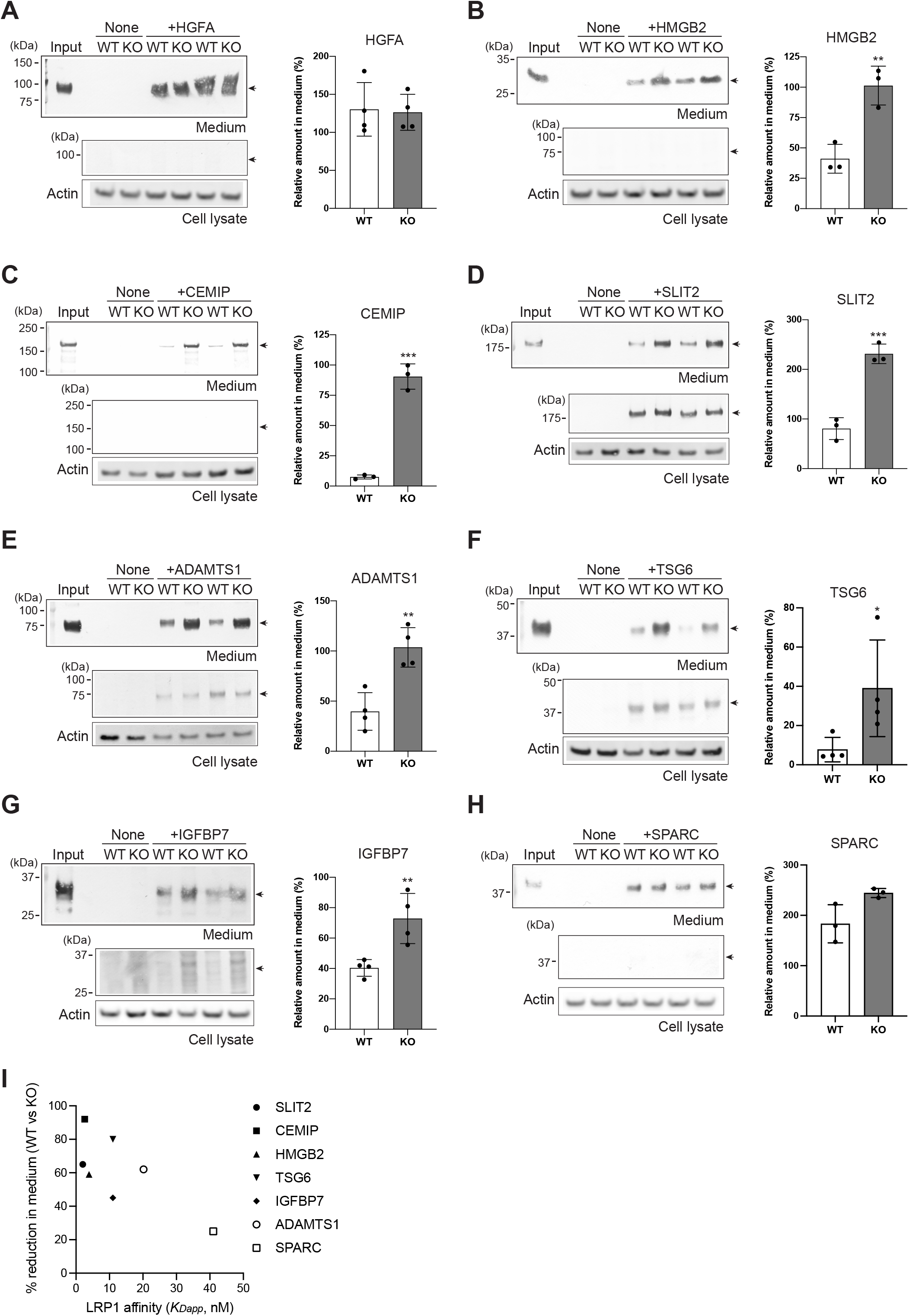
LRP1-mediated endocytic clearance of novel LRP1 ligands. Wild-type (WT) and LRP1 deficient (KO) MEFs (n = 3 or 4 each for *A, D, F* and *H*, or *B*, *C*, *E* and *G*, respectively) were cultured for 8 h in the absence (None) or presence of either 20 nM SLIT2 (*A*), ADAMTS1 (*B*), HGFA (*C*), HMGB2 (*D*), IGFBP7 (*E*), CEMIP (*F*), TSG6 (*G*) or SPARC (*H*) for 8 h. Representative Western blotting analysis of these proteins in the media and cell lysate detected using the anti-His tag antibody (*A*, *C*, *D*, *E*, *G* and *H*) or anti-FLAG M2 antibody (*B* and *F*)(*left panels*). Beta-actin in the cell lysate was detected by Western blotting using an anti-actin antibody. Band intensity was quantified by densitometry and the amount of each protein was expressed as a percentage of the amount of each protein at 0 h (Input), which was taken as 100% (*righ tpanels*). Circles represent a replicate and bars show the mean ± *SD*. *, *p* < 0.05, **, *p* < 0.01, ***, *p* < 0.001 by 2-tailed Student’s t test. *p* values were evaluated by 2-tailed Student’s t test. *I*, The estimated affinity of SLIT2, CEMIP, HMGB2, TSG6, IGFBP7, ADAMTS1 and SPARC for LRP1 (*K_D,app_*) was plotted against the percentage reduction of their amounts in the conditioned medium of WT compared to LRP1 KO MEFs.

**Table IV.** Characterization summary of newly identified LRP1 ligands

| | $K_{D,app}$ values<br>for LRP1 binding<br>(nM) | Exogenous ligands<br>in MEFs medium<br>(% vs LRP1 KO MEFs) | Endogenous levels in chondrocyte medium<br>with sLRP1-II (fold change vs control) | |
| --- | --- | --- | --- | --- |
|  |  |  | Secretome | Western blotting |
| SLIT2 | 2.0 | ↓ 65*** | ↑ 99.0**** | ↑ 2.5 |
| CEMIP | 2.6 | ↓ 92*** | ↑ 42.2**** | ↑ >15681 |
| HMGB2 | 3.9 | ↓ 59** | ↑ 10.7*** | ↑ >25.1 |
| HMGB1 | 6.4 | ND | ↑ 3.9* | ND |
| TSG6 | 11.0 | ↓ 80* | ↑ 2.5* | ↑ 2.8 |
| IGFBP7 | 11.0 | ↓ 45** | ↓ 3.7*** | NS |
| ADAMTS1 | 20.2 | ↓ 62** | ND | ↑ 10 |
| SPARC | 41.0 | NS (↓ 25) | ↓ 4.9** | ND |
| LIF | >200 | ND | ↑ 21.4**** | ND |
| HGFA | >500 | No change | ↑ 18.2**** | ND |
| Progranulin | No binding | ND | ↑ 13.6** | ND |
| Fibulin-1 | No binding | ND | ↓ 7.5*** | ND |

LRP1-mediated endocytic regulation of endogenously produced ADAMTS1 and IGFBP7 was further investigated by Western blot analysis of conditioned medium of human chondrocytes treated with 5 nM sLRP1-II for 24 h. The sLRP1-II treatment increased the levels of ADAMTS1 in the medium of chondrocytes approx. 10-fold compared to the control (**Suppl Fig 4A**), whereas no significant changes were observed for IGFBP7 (**Suppl Fig 4B**).

### High-affinity LRP1 ligands are tightly regulated by LRP1 in knee articular cartilage tissue *in vivo*

Candidate ligands were further validated *in vivo*. Given the complexity of extracellular environment and a possible competition between LRP1 and sulphated glycosaminoglycans (GAGs) in articular cartilage [43, 44], we investigated newly identified high-affinity LRP1 ligands SLIT2 and CEMIP in knee articular cartilage of LRP1 conditional KO mice. Due to the early embryonic lethality of *LRP1* deletion [6], we have established a transgenic mouse line harbouring floxed LRP1 and the ROSA26^(cre/ERT2)^ promoter (*LRP1^(flox/flox)^*/*ROSA26^(cre/ERT2)^*)(**Fig 7A**). The control (LRP1^flox/flox^) and homozygote (LRP1^flox/flox^Rosa^Cre/ERT2^) conditional LRP1 KO mice were given tamoxifen by intraperitoneal (IP) injection and knee joint tissue were collected 3 days after the third dose of tamoxifen. qPCR analysis of total RNA extracted from the mouse skin tissue showed >79% reduction of *Lrp1* mRNA in the KO compared to wild-type tissue (**Fig 7B**). Histological investigation of LRP1 in knee joint tissue revealed that LRP1 is abundantly expressed in the cell membrane of chondrocytes at the superficial area of cartilage and of the cells in the medial meniscus but these immunosignals were markedly diminished in the KO mice (**Fig 7C**). SLIT2 and CEMIP were detected in some chondrocytes in the control whereas stronger immunosignal of these ligands was observed in LRP1 KO tissue (**Fig 7D**). Like SLIT2 and CEMIP, connective tissue growth factor (CTGF), an important LRP1 ligand [45, 46], showed a negative correlation between absence of LRP1 and its level in cartilage *in vivo*.

**Figure 7.**
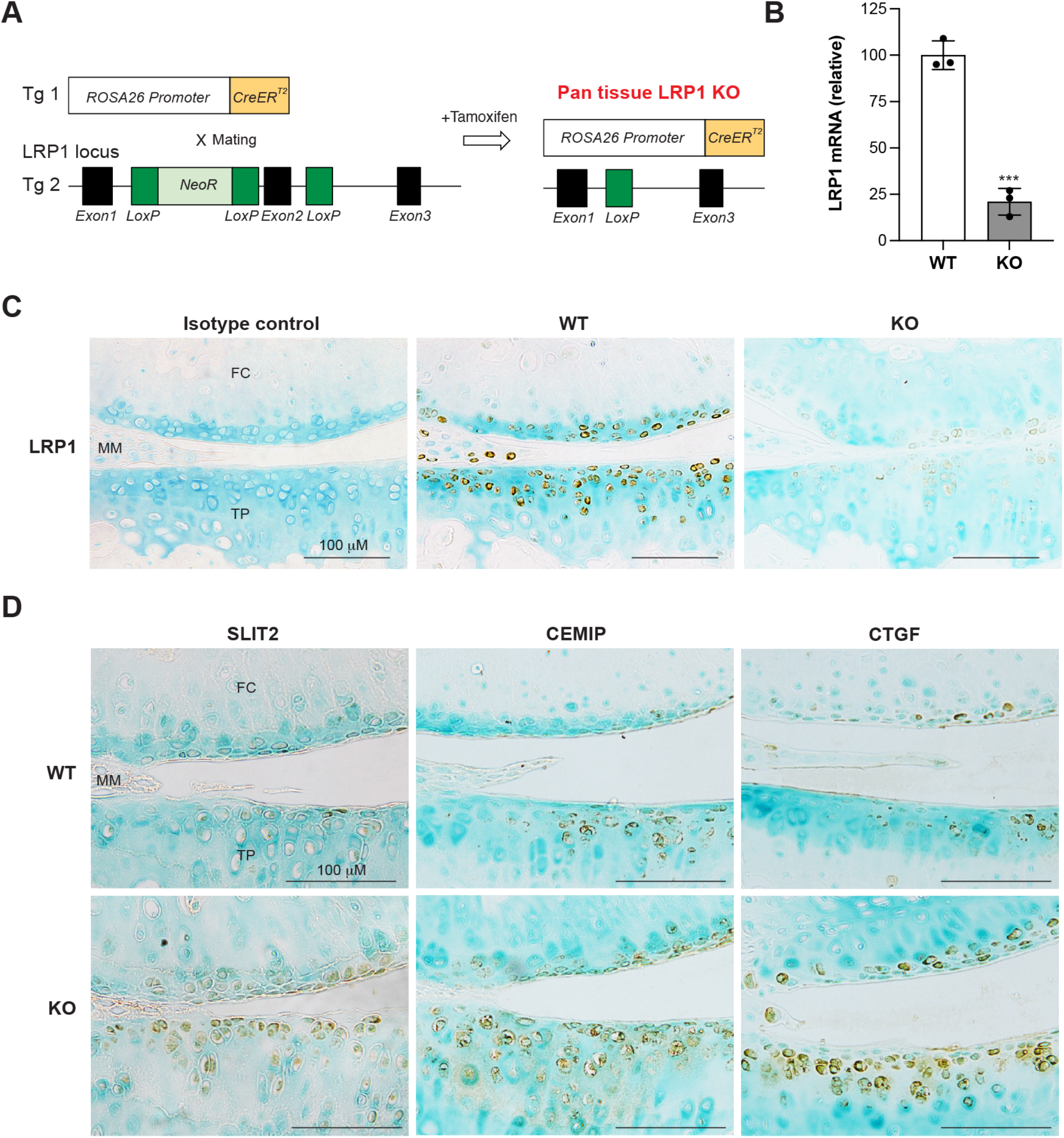
High-affinity LRP1 ligands are tightly regulated by LRP1 in knee articular cartilage tissue in vivo. *A*, Schematic diagram showing the constructs used to generate pan-tissue LRP1 conditional knockout mice. Transgenic mouse lines harbouring ROSA26^(cre/ERT2)^ promoter (Tg 1) and floxed *LRP1* (Tg 2) were used to establish a pan-tissue LRP1 conditional KO mice (*LRP1^(flox/flox)^/ROSA26^(cre/ERT2)^*). Tamoxifen activates *CreER^T2^*, which translocates into nucleus to excise *LRP1* gene. *B-D*, *LRP1^(flox/flox)^*/*ROSA26^(cre/ERT2)^*(KO) and *LRP1^(flox/flox)^* (WT) mice (n = 3 each) were administrated by IP injections for three times over a week with 2 and 4 days interval. The mice were culled 3 days after the final dose of tamoxifen. *B*, TaqMan real-time PCR showing relative levels of mRNA for LRP1 in the skin tissue. Circles represent individual cartilage donors and bars show the mean ± *SD*. ***, *p* = 0.0002 by 2-tailed Student’s t test. *C and D*, Representative immunohistochemical staining images of LRP1 (*C*), SLIT2, CEMIP and CTGF (*D*) in knee articular cartilage. Scale bar, 100 μm. *FC*, femoral condyle. *MM*, medial meniscus. *TP*, tibial plateau.

## Discussion

This study found that inhibition of LRP1-mediated endocytosis in human chondrocytes results in cell death, alteration of the entire secretome and transcriptional modulation. Our systematic approach revealed the extent of the chondrocyte LRP1 ligandome that illustrates novel biological roles of LRP1 in cartilage and potentially other tissues, and the pathological conditions when LRP1 ligands are present in the tissue for extended periods.

We showed that both sLRP1, which acts as a soluble decoy receptor and competes with endogenously expressed LRP1, and RAP, which binds to LRP1 and competes with endogenous ligand binding, induce chondrocyte cell death (**Fig 1**). Previous studies have indicated a role of LRP1 in cell survival and death [47–49]. In agreement with these studies, we found pro-cell survival role of LRP1 in human articular chondrocytes. Furthermore, several studies demonstrated cell survival and death functions for both previously reported and newly identified LRP1 ligands such as TIMP3 [50–55], ADAMTS5 [56], CTGF [57, 58], CEMIP [59], SLIT2 [60, 61], IGFBP7 [62–64] and HMGB2 [65]. The mechanisms by which LRP1 ligands regulate cell survival in chondrocytes remain unclear but the prolonged presence of one or several LRP1 ligands in the extracellular milieu may gradually alter cellular environment and metabolism leading to cell death.

The correlation between the affinity of LRP1 and the rate of endocytosis was previously demonstrated for ADAMTS4 and 5 isoforms [21]. The data presented here also supported this notion for various LRP1 ligands. Importantly, several of LRP1 ligands bind to both LRP1 and sulphated GAGs, with their extracellular availability determined by their relative affinity for each [43, 44, 66]. The extracellular environment in articular cartilage is a complex matrix where chondrocytes are surrounded by a thin layer of pericellular ECM followed by an abundant ECM mainly consisting of aggrecan proteoglycan and type II collagen fibrils. Nonetheless, markedly increased SLIT2, CEMIP and CTGF in knee articular cartilage from conditional LRP1 knockout mice (**Fig 7**), indicate that LRP1 is a master regulator for extracellular availability of high-affinity ligands in the tissue *in vivo*.

The chondrocyte secretome analysis revealed a huge impact of LRP1 blockade with significant changes in 368 molecules. LRP1 ligands can affect each other, as well as molecules that do not interact with LRP1, in several ways including transcriptional modulation, complex formation and proteolytic degradation. It has been reported that RAP increases mRNA levels of pro-inflammatory mediators such as TNFα, interleukin-6 and C–C motif chemokine ligand 2 in macrophages [36]. Furthermore, considering functional diversity of LRP1 ligands, transcriptional modulation of several molecules shown in this study could be the tip of the iceberg. Negligible binding of progranulin and fibulin-1C to LRP1 suggests that these molecules indirectly interact with LRP1 *via* LRP1 ligands (**Fig 5**). This notion is supported by the interaction networks formed by co-isolated proteins (**Fig 4D**). To date, several extracellular proteolytic enzymes, which cleave a broad range of substrates, have been identified as LRP1 ligands [4, 5, 44]. Proteolytic degradation by LRP1 ligands is thus likely to contribute to reduction of the number of secreted proteins in particular ECM molecules. Indeed, levels of versican, nidogen-2 and biglycan, all previously reported substrates for ADAMTS1[67–69], were reduced in the medium of chondrocytes treated with sLRP1-II (**Table II**).

This study generated new information about the pathobiological functions of the key molecules that are regulated by LRP1-mediated endocytosis and their connection to the development of OA. Extracellular HMGB molecules function as alarmins, which are endogenous molecules released upon tissue damage to activate the immune system. HMGB2 KO mice exhibit earlier onset of OA and a more severe progression, which is associated with a profound reduction in cartilage cellularity [65]. By contrast, HMGB1 acts as a late mediator of inflammation and contributes to prolonged and sustained systemic inflammation in subjects with rheumatoid arthritis (see [70] for review). Importantly, hyaluronan degradation is highly associated with the risk of OA progression [71, 72]. CEMIP functions as a hyaluronan depolymerase [73] and is highly relevant to OA pathogenesis [74–76]. Expression levels of CEMIP and CTGF are correlated with disease severity in OA patients [77–79], and we found that their levels are increased in cartilage from LRP1 KO mice *in vivo* (**Fig 7**). Notably, hyaluronan is endocytosed *via* clathrin-coated pits and degraded in vesicles in HEK293 cells overexpressing CEMIP [80]. LRP1 not only regulates extracellular availability of CEMIP but may also play a role in transporting both CEMIP and hyaluronan to the coated pits where hyaluronan is degraded. TSG6 also binds to hyaluronan [81] and its protein and mRNA levels are increased in OA cartilage [82, 83]. In the presence of inter-α-inhibitor (IαI), which was identified as a LRP1 ligand candidate (**Table III**), TSG6 catalyzes formation of anti-inflammatory stabilised hyaluronan-aggrecan assembly [84–86]. However, it blocks hyaluronan-aggrecan assembly in the absence of IαI [87]. Furthermore, our previous study demonstrated that IαI is proteolytically cleaved by ADAMTS5, MMP3, MMP7 and MMP13 [88]. Considering that CEMIP, TSG6, ADAMTS5 and MMP13 are all regulated by LRP1, increased LRP1 shedding in OA effectively leads to cartilage destruction by both increasing extracellular activity of cartilage-degrading proteinases [19] and destabilizing cartilage ECM assembly.

Accumulating evidence suggests a role of LRP1 in the regulation of Wnt signalling, which governs a myriad of biological processes underlying the development and maintenance of adult tissue homeostasis. Macrophage LRP1 increases Wnt/β-catenin pathway by directly binding to and effectively removing secreted frizzled-related protein 5, which prevents Wnt binding to its receptor [86]. LRP1 also stimulates Wnt/β-catenin pathway and prevents intracellular cholesterol accumulation in fibroblasts [87]. A recent study also demonstrated that LRP1 regulates expression of Wnt family member 4 in chondrocytes through transforming growth factor-β1 signalling in chondrocytes [88]. On the other hand, it has been reported that LRP1 interacts with Frizzled1 and downregulates Wnt/ β-catenin pathway in HEK293T cells [89]. Newly identified LRP1 ligands in this study are also involved in the Wnt pathway. For example, HMGB2 enhances the binding of lymphoid enhancer-binding factor 1 (Lef-1), a crucial transcription factor for Wnt/β-catenin pathway, to its target sequence and potentiates transcriptional activation of the Lef-1-β-catenin complex [89]. The HMG domain within HMGB2 is crucial for interaction with Lef-1, suggesting that both HMGB2 and HMGB1 may be involved in this function. CEMIP also enhances Wnt/β-catenin signalling and bone formation of osteoblastic stem cells [90]. Moreover, we have identified WNT5a and WNT11 as LRP1 ligand candidates (**Table III**). These studies emphasise a role of LRP1 in the Wnt pathway and we are currently investigating how the interaction between LRP1 and these WNTs affects Wnt signalling pathways.

This study provided important information about possible roles of LRP1 in other tissue contexts and diseases. SLIT2 binds to Roundabout 1 and 2 receptors [91] and plays a crucial role in neuronal outgrowth in embryogenesis [92, 93]. HGFA, which directly binds to LRP1 and was increased in the medium of chondrocytes by sLRP1-II treatment, is a serine protease responsible for converting HGF into the active two-chain form [94–96]. HGF is considered to play a major role in the repair and regeneration of various tissues, including the liver, kidney, lung, and stomach [97]. Notably, both HGF activation and tissue regeneration were markedly impaired in injured intestinal tissue of HGFA-deficient mice [95, 96], indicating the important function of HGFA during tissue repair process. Although HGFA is the major HGF activator, previously reported LRP1 ligands including urokinase-type plasminogen activator, tissue plasminogen activator and coagulation factor XI were also able to activate HGF *in vitro* [98, 99]. Another newly identified LRP1 ligand, ADAMTS1, is broadly expressed and essential for normal growth, structure, and function of the kidney, adrenal gland, female reproductive-organ, and myocardial trabeculation during heart development (see [100] for review), and limb joint development [101]. Interestingly, recent studies revealed emerging roles of CEMIP in bone development [90, 102], inflammation and antimicrobial activity [103]. A study using CEMIP-deficient mice revealed that hyaluronan metabolism by CEMIP is involved in endochondral ossification during postnatal development by modulation of angiogenesis and osteoclast recruitment at the chondro-osseous junction [102]. In another study, CEMIP-deficient mice showed no hyaluronan digestion and significantly less evidence of skin infection after an intradermal bacterial challenge by *Staphylococcus aureus* [103]. This information about the physiological functions of newly identified ligands implies LRP1 in organ morphogenesis, tissue regeneration and infection.

The methodology established in this study could be useful for other tissue contexts and diseases. One of the limitations of such an approach is that several LRP1 ligands were discovered with conflicting functions. For example, both proteases and their specific inhibitors were identified as LRP1 ligands. It is thus difficult to predict any net outcome of LRP1-mediated endocytosis and its dysregulation from the results of this study. However, our elucidation of a cartilage-specific ligandome opens potential new avenues of therapeutic intervention in diseases characterized by dysregulated LRP1-mediated endocytosis. By designing agents able to modulate the endocytosis of cartilage specific LRP1 ligands, it will be possible to counteract the negative effects of LRP1 dysregulation. For example, variants of TIMP3 resistant to LRP1-mediated endocytosis [104] and soluble LRP1 subcluster that selectively block TIMP3 endocytosis [35] have been already engineered to combat excessive ADAMTS5 activity. Combination of functional studies using cells or animal models and more complete tissue specific LRP1 ligandome identification is required for better understanding of the physiological functions of LRP1, and the detrimental effects of its dysregulation.

## Experimental Procedures

### Reagents and antibodies

The sources of materials used were as follows: Pierce^™^ Anti-c-Myc magnetic beads, RNase Inhibitor, 4-12% Bis-Tris NuPage Gels and fetal bovine serum (FBS) from Thermo Fisher Scientific (Waltham, MA); purified human full-length LRP1 from BioMac (Leipzig, Germany); recombinant human progranulin (C-terminal His tag, 2420-PG), SLIT2 (C-terminal His tag, 8616-SL), HGFA (C-terminal His tag, 1514-SE), TSG6 (C-terminal His tag, 2104-TS), anti-His tag mouse monoclonal antibody (MAB050) and the basic-based antigen unmasking solution (CTS013) from R&D Systems (Minneapolis, MN); anti-FLAG M2 mouse monoclonal antibody (F1804), protease Inhibitor cocktail consisting of aprotinin, bestatin, E-64, leupeptin and pepstatin A (P1860), tamoxifen free base, corn oil, lysis buffer, REDtaq ReadyMix PCR reaction mix, DPX mounting media, Dulbecco’s Modified Eagle’s Medium/Nutrient Mixture F-12 Ham (DMEM/F12) and bovine serum albumin (BSA) from Sigma (Dorset, UK); recombinant human HMGB1 (C-terminal His tag, 10326-H08H), IGFBP7 (C-terminal His tag, 13100-H08H), SPARC (C-terminal His tag, 10929-H08H) and LIF (C-terminal His tag, 14890-H08H) from Sino Biological (Beijing, China); recombinant human HMGB2 (C-terminal His tag, TP720732) from ORIGENE (Rockville, MD); anti-rabbit alkaline phosphatase-linked antibody, anti-mouse alkaline phosphatase-linked antibody, anti-goat alkaline phosphatase-linked antibody and alkaline phosphatase substrate (5-bromo-4-choloro-3-indolyl 1-phosphate and nitroblue tetrazolium) from Promega (Southampton, UK); a hydroxamate-based MMP inhibitor CT-1746 (N1-[2-(S)-(3,3-dimethylbutanamidyl)]-N4-hydroxy-2-(R)-[3-(4-chlorophenyl)-propyl]-succinamide) from UCB Celltech (Slough, UK); anti-TSG6 rat monoclonal antibody (sc-65886), goat polyclonal anti-actin antibody (I-19) from Santa Cruz Biotechnology (Dallas, TX); anti-LRP1 β-chain rabbit monoclonal (ab92544), anti-CEMIP rabbit polyclonal (ab98947), anti-HMGB2 rabbit monoclonal (ab124670), anti-SLIT2 rabbit monoclonal (ab134166), anti-ADAMTS1 rabbit polyclonal (ab216977), anti-CTGF rabbit polyclonal (ab6992) and anti-MMP1 mouse monoclonal (ab25483), anti-MMP3 mouse monoclonal (ab38907) antibodies from Abcam (Cambridge, UK); control rabbit IgG (I-1000-5), ImmPRESS peroxidase-micropolymer conjugated horse anti-rabbit IgG, Goat serum, Vectastain solution, methyl green, 3,3′-diaminobenzidine from Vector Laboratories (Burlingame, CA); anti-CEMIP rabbit polyclonal antibody (21129-1-AP) from Proteintech (Rosemont, IL). Recombinant human RAP [22], sLRP1-II [21], ADAMTS1 [67], ProMMP1 (E200A) [105], ProMMP3 (E202A) [105] were prepared as previously reported. All other reagents used were of the highest analytical grade available.

### Purification of full-length sLRP1

LRP1 was purified from human placenta by RAP affinity chromatography (Mogensen *et al*, under review). Briefly, for column preparation, C-terminally his-tagged RAP [22] was expressed in *Escherichia coli*, harvested by centrifugation, and resuspended in 50 mM HEPES (pH 7.4) and 150 mM NaCl (HBS) supplemented with complete EDTA-free Protease Inhibitor Cocktail (Roche) prior to lysis by sonication on ice. For RAP purification, the lysate was applied to a HisTrap column (Cytiva Life Sciences), and bound proteins were eluted using a stepwise increase of imidazole concentrations from 12.5-250 mM. A total of 3 mg RAP was coupled to a 1-ml HiTrap NHS-activated HP column (Cytivia Life Sciences) according to the manufacturer’s protocol. LRP1 was extracted from fresh human placenta as previously described [106], and the placental mixture was applied to the RAP affinity column equilibrated in HBS with 5 mM CaCl_2_ and 0.05 % Tween. Bound proteins were eluted by 100 mM CH3COONa (pH 5.0) and 0.5 M NaCl and further purified using a Sephacryl S-300 HR (Cytiva Life Sciences) size exclusion column equilibrated in 100 mM CH3COONa (pH 5.5) and 150 mM NaCl.

### Purification of CEMIP

The expression construct for full-length human CEMIP (Uniprot ID: Q8WUJ3) containing a FLAG tag at the C-terminus in pReceiver-M39 was purchased from GeneCopoeia. HEK293 expressing the SV40 large T antigen were cultured in DMEM/F12 with 10% FBS, 1 U/mL Penicillin and 0.1 mg/mL Streptomycin (Pen/Strep)(Sigma) at 37°C, with 5% CO_2_. Constructs were transiently transfected using TransIT2020 (Mirus Bio) following the manufacturer’s protocol. After the transfection, cells were incubated for 3 days in the presence of 100 μg/ml heparin from porcine intestinal mucosa (Sigma, H3393) to prevent LRP1-mediated endocytic clearance before harvesting. Conditioned medium (2 L) was harvested and centrifuged for 20 mins at 1500 x g, followed by filtration (0.45 μm) to remove cell debris. The conditioned medium was passed over a 2-ml anti-FLAG M2 Affinity Gel (Sigma, A2220) at 4°C. The column was first washed extensively with 50mM Tris-HCl (pH 7.5) containing 1 M NaCl, 5 mM CaCl_2_, and 0.02% Brij-35. The bound material was eluted with 200 μg/mL FLAG peptide (Sigma, F3290). The purified samples were then passed through a PD-10 column (GE Healthcare, UK) pre-equilibrated in 50 mM HEPES, 150 mM NaCl, 5 mM CaCl_2_, and 0.02% Brij-35. Fractions containing pure CEMIP were pooled, concentrated and stored at −80°C before activity assays. Protein concentration was measured using Nanodrop and calculated according to Beer–Lambert law, using extinction coefficient 226,115 M-1cm-1 using the Expasy ProtParam web tool.

### Purification of Fibulin1C

The cDNA coding for fibulin1C (Uniprot number P23142-4) with a C-terminal 6x His tag was custom-synthesized by Invitrogen and transiently transfected in HEK293 T cells using polyethylenimine in OPTIMEM (Invitrogen) containing 2 mM CaCl_2_. After 72h, fibulin1C was concentrated 15-fold using a Lab scale TFF system (Merck) and purified using a Ni-sepharose column (GE Healthcare) equilibrated with 3 column volumes TBS (20 mM Tris-HCl (pH 7.4), 150 mM NaCl). Following binding, the column was washed with TBS containing 10 mM imidazole and bound proteins were eluted using a linear gradient (10-300 mM) of imidazole. Eluted fractions containing recombinant proteins were subjected to SDS-polyacrylamide gel electrophoresis (PAGE), pooled, concentrated on Amicon Ultra spin columns (100 kDa cut-off) and dialyzed extensively against TBS. Purified fibulin1C was stored at −80°C. Protein concentration was measured using Nanodrop and calculated according to Beer–Lambert law, using extinction coefficient 35,750 M-1cm-1 using the Expasy ProtParam web tool.

### Isolation and culture of human articular chondrocytes

Healthy (normal) articular cartilage was obtained from the Stanmore BioBank, Institute of Orthopaedics, Royal National Orthopaedic Hospital, Stanmore from patients following informed consent and approval by the Royal Veterinary College Ethics and Welfare Committee (Institutional approval URN 2010 0004H). Articular cartilage was obtained from the femoral condyles of the knee following amputation due to soft tissue sarcoma and osteosarcoma with no involvement of the cartilage. Tissues were obtained from 5 patients (3 males aged 18, 23 and 57 years; 2 females aged 19 and 68 years) and chondrocytes were isolated as described previously [26]. The cells were grown and maintained in DMEM/F12 media supplemented with 10% FBS at 37 °C with 5% CO_2_. Both primary and passaged (< 3 times) human cells were used in the study.

### Cell-survival assay

Human chondrocytes were grown on 24 well plate with DMEM/F12 containing 10% FBS. Once the cells reached 90% confluency, the medium was removed and the cells were rested in 0.4 ml of serum-free (SF) DMEM/F12 for 1 day. The cells were then incubated with 0.4 ml of fresh SF DMEM/F12 in the absence or presence of 10 nM full-length sLRP1 (BioMac) or 500 nM RAP for 0-72 h. Cell images after 72-h incubation were acquired by Nikon Eclipse Ti-E and Diaphot with ToupCam camera (Nikon, Tokyo, Japan). Cell viability at 72-h incubation was measured by MTS cell proliferation colorimetric assay (CellTiter 96^®^ AQueous One Solution Cell Proliferation Assay system, Promega, G3582, Italy). Cell numbers at 0, 24, 48 and 72-n incubation were counted by the hemocytometer (Hausser Scientific, PA, USA). For the experiment using sLRP1-II, cells were maintained on 48 well plate and incubated with 0.2 ml of SF DMEM/F12 in the absence or presence of 0-40 nM sLRP1-II for 24 h.

### Liquid Chromatography with tandem mass spectrometry (LC-MS/MS)

Samples were denatured and reduced in 8 M Urea containing 5 mM Dithiothreitol for 1 h, alkylated with 15 mM Iodoacetamide for 1 h and diluted 10-fold with 50 mM NH_4_HCO_3_. Sequencing grade trypsin was added (1:50 w/w) and the sample was digested overnight 37 °C. The sample was desalted using self-packed reverse phase micro-columns containing C18 column material (EmporeTM). LC-MS/MS was performed using an EASY-nLC 1000 (Thermo Scientific) connected to an Q Exactive Plus mass spectrometer (Thermo Scientific). The peptides were separated on a 15-cm analytical column (75 μm inner diameter). The column was packed in-house with ReproSil-Pur C18-AQ 3 μm resin (Dr. Maisch GmbH, Ammerbuch-Entringen, Germany). The flow rate were 250 nl/min and a 50 min gradient from 5% to 35% phase B (0.1% formic acid, 90% acetonitrile). The acquired MS data were processed in Proteome Discoverer 2.4 (Thermo Scientific). The data was searched against the human proteome (UniProt) using the Sequest HT search engine. Search parameters were trypsin as protease; minimum peptide length of 6 residues; Precursor Mass Tolerance of 10 ppm; Fragment Mass Tolerance of 0.02 Da and dynamic modifications with oxidation on Met. The search results were used for precursor ion quantification without normalization. Statistical analysis using the search results and intensities from Proteome discover was made with Perseus (v1.6.5.0) [107]. Initially the results were filtered based on the Protein FDR to remove low confidence IDs. The raw intensities were log2 transformed and missing values were replaced based on the normal distribution of the entire dataset. A two-sample t-test was performed to identify significant changes and calculate the fold change. Volcano plot analysis was performed using GraphPad Prism 9. Several proteins were identified in the sLRP1-II preparation thereby all intensities were normalized to the intensity of sLRP-II preparation in the sample. Exogenously added sLRP1-II and associated RAP were excluded from the data set of secretome and co-immunoprecipitation analysis.

### Western blot analysis of secreted proteins produced by human chondrocytes

Human chondrocytes were incubated for 24 h with SF DMEM/F12 containing CT1746 (100 μM) and the protease inhibitor cocktail (1/500) in the presence and the absence of 5 nM sLRP1-II. After incubation, media were collected, and the protein was precipitated with trichloroacetic acid and dissolved in 20 μl of 1x SDS-sample buffer (50 mM Tris-HCl pH 6.8, 5% 2-mercaptoethanol (2ME), 2% SDS and 10% glycerol). All samples were analyzed by SDS-PAGE and Western blotting using a specific antibody against each protein. Immunoreactive bands were quantified using NIH ImageJ, and the relative amount of each ligand in the medium in the presence and absence of sLRP1-II was calculated.

### Quantitative reverse transcriptase-polymerase chain reaction (qRT-PCR)

Human chondrocytes were incubated for 24 h with SF DMEM/F12 containing CT1746 (100 μM) and the protease inhibitor cocktail (1/500) in the presence and the absence of 200 nM RAP, 10 nM full-length sLRP1 or 10 nM denatured full-length sLRP1 (preincubated at 98 °C for 10 min) for 0-24 h. cDNA was generated from these cells using a reverse-transcription kit (Applied Biosystems, CA, USA) and random primers from RNA extracted and prepared using the RNeasymini kit (Qiagen, CA, USA) following the manufacturer’s guidelines. cDNA was then used for real time PCR assays using TaqMan technology. The ΔΔthreshold cycle (ΔΔCt) method of relative quantitation was used to calculate relative mRNA levels for each transcript examined. The 60S acidic ribosomal protein P0 (RPLP0) gene was used to normalize the data. Pre-developed primer/probe sets for MMP1, MMP3, MMP13, ADAMTS4, ADAMTS5, TIMP3 and RPLP0 were purchased from Applied Biosystems.

### Isolation of LRP1 ligands in human chondrocytes

Human chondrocytes were grown on 12 well plate in DMEM/F12 containing 10% FBS. Once the cells reached 90% confluency, the medium was removed and the cells were rested in 1 ml of SF DMEM/F12 for 1 day. The cells were then incubated with 1 ml of SF DMEM containing the broad-spectrum hydroxamate metalloproteinases inhibitor CT1746 (100 μM) and the protease inhibitor cocktail (a mixture of aprotinin, bestatin, E-64, leupeptin and pepstatin A)(SIGMA, P1860) in the presence or absence (control) of 5 nM sLRP1-II for 24 h. The conditioned medium was collected into BSA pre-coated 1.5 ml eppendorf tubes and centrifuged briefly by table-top centrifuge to remove cell debris. 900 μl of the supernatant was collected into new BSA pre-coated 1.5 ml tubes and 100 μl of the supernatant was used for the secretome analysis. 800 μl of the supernatant was further incubated with 50 μl of anti-c-Myc paramagnetic Dynabeads at 4°C for 1 h. sLRP1-II and the magnetic beads complexes were then isolated using the magnetic stand, washed with the buffer consisting of 50 mM Tris-HCl (pH 7.5), 150m M NaCl, 5 mM CaCl_2_ (TNC), and 0.02% Brij-35 (TNCB) for three times, and eluted with 200 μl of 0.1 M glycine buffer (pH 3.0) for 10 min at 4°C. The eluents were neutralised with 10 μl of 1 M Tris-HCl (pH 8.0).

### Functional analysis of proteomics data

The PANTHER Classification System [39] was used to determine gene ontology and biological processes of the identified proteins. The functional and physical networks analysis for the identified proteins were performed using the STRING version 11.0 [40].

### Enzyme-linked immunosorbent assay (ELISA) for the binding of recombinant LRP1 ligand candidate proteins to full-length sLRP1

Purified human full-length LRP1 (10 nM in 100 μl of TNC) was coated overnight at 4 °C onto microtiter plates (Corning, NY). Wells were blocked with 3% BSA in TNC (1 h; 37 °C) and washed in TNC containing 0.05% Brij-35 after this and each subsequent step. Wells were then incubated with various concentrations of recombinant proteins in blocking solution for 3 h at room temperature. Bound proteins were detected using anti-FLAG M2 antibody or anti-His tag antibody (1 h; room temperature) and then with a secondary antibody coupled to horseradish peroxidase (1 h; room temperature). Hydrolysis of tetramethylbenzidine substrate (KPL, Gaithersburg, MA) was measured at 450 nm using a FLUOstar Omega (BMG Labtech). Mean values of technical duplicate were normalized by subtracting the amount of recombinant protein bound to control well that was not coated with LRP1. Extrapolated *K_D,app_* values were estimated based on one-phase decay nonlinear fit analysis using GraphPad Prism 9.

### Endocytosis assay using WT and LRP1 deficient MEFs

WT and LRP1 KO MEFs were generated as described previously [42] and kindly provided by Professor Dudley Strickland (University of Maryland School of Medicine). Cells (1×10^4^/well) cultured in 96-well plates (pre-coated with 0.1% gelatin for 3 h) were rested in 100 μl of DMEM for 1 day. The medium was replaced with 50 μl of fresh DMEM containing recombinant proteins (20 nM). After 8 h, 30 μl of medium were collected and mixed with 10 μl of 4x SDS-sampling buffer (200 mM Tris-HCl (pH 6.8), 8% SDS and 40% glycerol) containing 10% 2ME. The medium was then completely removed and the cells were lysed with 50 μl of 2x SDS-sampling buffer containing 5% 2ME. All samples were analyzed by SDS-PAGE under reducing conditions and immunoblotting using anti-His tag or anti-FLAG M2 antibody. Immune signals for exogenously added proteins in the medium were quantified using ImageJ and the amount of the protein remaining in the medium after 8 h incubation was calculated as a percentage of the amount before incubation.

### Establishment of pan-tissue LRP1 conditional knockout mice

LRP^flox/flox^ (129S7-Lrp1^tm2Her^/J) mutant mice carrying a floxed Neo cassette and a single loxP site downstream of exon2 of the targeted *LRP1* gene, were crossed with tamoxifen-inducible *Cre* recombinase transgene R26CreERT2 (129-Gt (ROSA)26Sor^tm1(cre/ERT2)Tyj^/J) mice to generate LRP1 conditional pan tissue KO mice. Both were provided by The Jackson Laboratory (012604 and 008463). Fifty mg tamoxifen free base (Sigma, T5648) was suspended in 500 μl 100% ethanol, diluted and fully dissolved in 4,500 μl of corn oil (Sigma, C8267) by sonication (final 10 mg/ml). 5-6 months-old mice were administrated 40 mg tamoxifen/Kg body weight by intraperitoneal injections for three times over a week with 2- and 4-days interval.

### Genotyping

Four weeks after birth, ear notches were collected from mouse pups to extract genomic DNA using the RED Extract-N-Amp Tissue PCR Kit (Sigma-Aldrich). Ear notches were incubated in the tissue preparation (25 μl/sample) and extraction solutions (100 μl/sample) at room temperature for 10 minutes. Ear notches were heated to 95 °C for 3 minutes and later mixed with the neutralization solution (100 μl/sample) to offset inhibitory substances before PCR. Samples were kept at 4°C until PCR. For R26-Cre ERT2 mouse genotyping, 300 ng of genomic DNA was mixed with 0.2 μl of Go Taq^®^G2 flexi DNA polymerase (PROMEGA), 5.0 μl of 5X Green Go Taq ^®^ Flexi buffer (PROMEGA), 3.0 μl MgCl2 (25mM) (PROMEGA), 1.0 μl dNTPs (100 mM, Meridian), 0.5 μl (20 μM) of each primer and nuclease-free water was added to make the reaction mixture up to 25 μl. Primer pairs for genotyping were as follows; ROSA Cre WT Forward (21306) 5′ - CTG GCT TCT GAG GAC CG - 3′, ROSA Cre WT Reverse (oIMR9021) 5′-CCG AAA ATC TGT GGG AAGTC - 3′, ROSA Cre Mut. Forward (oIMR3621) 5′-CGT GAT CTG CAA CTC CAG TC-3′, ROSA Cre Mut. Reverse (oIMR9074) 5′-AGG CAA ATT TTG GTG TAC GG - 3′. Cycle conditions were as follows: Genotyping – 1 cycle of 94°C for 2 min, 10 cycles of 94°C for 20 s; 65°C for 15 s (−0.5°C per cycle) (touchdown); 68°C for 10 s, and 28 cycles of 94°C for 15 s; 60°C for 15 s; 72°C for 10 s, followed by a final cycle of 72°C for 2 mins. PCR products were separated by gel electrophoresis and imaged using a BioRad Gel Doc XR+ System. For LRP1 floxed mice mouse genotyping, 100 ng of genomic DNA was mixed with 10 μl of REDExtract-N-Amp^™^ PCR ReadyMix^™^, 0.5 μl (20 μM) of each primer and nuclease-free water was added to make the reaction mixture up to 20 μl. Primer pairs for genotyping were as follows; LRP1 Flox forward 5′-CATACCCTCTTCAAACCCCTTCCTG - 3′, LRP1 Flox Reverse 5′-GCAAGCTCTCCTGCTCAGACCTGGA - 3′. Cycle conditions were as follows: Genotyping - 1 cycle of 94°C for 3 min, 35 cycles of 94°C for 30 s; 65°C for 30 s; 72°C for 30 s, followed by a final cycle of 72°C for 2 min. PCR products were separated by gel electrophoresis and imaged using a BioRad Gel Doc XR+ System.

### Tissue processing for histology

Animals were killed by CO_2_ and knee joints were removed by using sanitised scissors. The tissues were fixed in 4%paraformaldehyde for 24 h, stored in 70% ethanol followed by decalcification with 10% EDTA (pH 8.0) for 2 weeks. The decalcified tissues were then processed through graded ethanol and xylene before being embedded in paraffin wax. 4.5 μm sections were cut using a rotary microtome RM2235 (Leica), adhered to microscope slides, then dried overnight at 37°C. Sections were dewaxed and rehydrated with xylene followed by a series of decreasing ethanol concentrations prior to the histological analysis.

### Immunohistochemical staining of LRP1 and LRP1 ligands

Slides were incubated with the basic-based antigen unmasking solution by placing a polypropylene Coplin staining jar filled with the respective retrieval solution into a water bath at 92-95 °C for 3-8 minutes. Slides were washed with distilled water followed by PBS and blocked with 0.3% hydrogen peroxide for 15 min at 37 °C. Slides were then incubated with Avidin solution for 15 minutes followed by the Biotin solution for 15 minutes at room temperature. After washing with PBS, slides were incubated with PBS containing 10% goat serum and 0.1% BSA for 3 hours at room temperature. Slides were then incubated with anti-LRP1 β-chain rabbit monoclonal antibody (ab92544), anti-SLIT2 rabbit monoclonal antibody (ab134166), anti-CEMIP rabbit polyclonal antibody (21129-1-AP) or anti-CCN2 rabbit polyclonal antibody (ab6992) for overnight at 4 °C. Rabbit IgG was used as an isotype control. Slides were washed with PBS containing 0.1% Tween 20 (PBST) three times for 10 minutes each. Slides were then incubated with secondary antibody ImmPRESS peroxidase-micropolymer conjugated horse anti-rabbit IgG for 30 minutes at room temperature followed by incubation with Vectastain solution (Vector Labs, PK-6100) for 30 minutes at room temperature. After wash with PBST three times for 10 minutes each slide was stained with 3,3′-diaminobenzidine and counterstained with Methyl green solution for 5 minutes followed by dehydration and mounted with DPX mounting media. Images were acquired using a Nikon Eclipse Ti-E microscope (Nikon, Tokyo, Japan).

## Supporting information

Supplementary table III

## Acknowledgements

We are grateful to the staff at the University of Liverpool Biomedical Services and Histology Units. We thank Dr Jaysh Dudhia (Royal Veterinary College, United Kingdom) for providing human normal cartilage. This work was supported by the Versus Arthritis (21447 to K.Y.) and the University of Liverpool Crossley Barnes Bequest fund (K.Y. and M.M.M.), the British Heart Foundation (FS/IBSRF/20/25032 to S.S.), a “Fondazione con il Sud” grant within the Brains to South program (2018-PDR-0799 to S.B. and S.D.S.), the VELUX FONDEN (00014557 to J.J.E.), the Danish Council for Independent Research-Medical Science (DFF-4004-00471 to J.J.E.), and the LEO Foundation and the Novo Nordisk Foundation (BIO-MS)(NNF18OC0032724 to J.J.E.).

## Author contributions

K.Y. designed and performed the experiments, acquired the data, wrote and edit the manuscript. C.S., M.M.M., E.H.M., A.M.E.G. and I.B.T. performed the experiments and acquired the data. S.B. and S.D.S. analysed the secretome data. S.S. and J.A. performed the experiments. G.B.G., J.J.E. and H.N. designed the experiments and edit the manuscript. All authors reviewed and revised the manuscript.

## Abbreviations

LRP1: low-density lipoprotein receptor-related protein 1
SLIT2: slit homolog 2 protein
CEMIP: cell migration-inducing and hyaluronan-binding protein
ECM: extracellular matrix
SMCs: smooth muscle cells
OA: osteoarthritis
TIMP: tissue inhibitor of metalloproteinase
MMP: matrix metalloproteinase
ADAM: a disintegrin and metalloproteinase
ADAMTS: ADAM with thrombospondin motifs
RAP: receptor-associated protein
HMG: high mobility group protein
HGFA: hepatocyte growth factor activator
IGFBP: insulin-like growth factorbinding protein
TNF: tumor necrosis factor
TSG6: TNF-inducible gene 6 protein
SPARC: secreted protein acidic and rich in cysteine
WT: wild-type
KO: knockout
MEFs: mouse embryonic fibroblasts
CTGF: connective tissue growth factor
GAGs: glycosaminoglycans
IL-1β: interleukine-1β
Lef-1: lymphoid enhancer-binding factor 1
IαI: inter-α-inhibitor
FBS: fetal bovine serum
BSA: bovine serum albumin
DMEM/F12: Dulbecco’s Modified Eagle’s Medium/Nutrient Mixture F-12 Ham
SF: serum-free
LC-MS/MS: liquid Chromatography with tandem mass spectrometry
SDS: sodium dodecyl sulphate
PAGE: polyacrylamide gel electrophoresis
2ME: 2-mercaptoethanol
qRT-PCR: quantitative reverse transcriptase-polymerase chain reaction

