## Supplementary materials for "Uncovering the ligandome of low-density lipoprotein receptor-related protein 1 in cartilage: a top-down approach to identify therapeutic targets"

**Supplementary Information**

**<Methods>**

*Reagents and antibodies*

The sources of materials used were as follows: the silver staining kit from Thermo Fisher Scientific (Waltham, MA); anti-ADAMTS1 mouse monoclonal antibody (TA804585) from ORIGENE (Rockville, MD); and anti-IGFBP7 rabbit polyclonal (ab74169) from Abcam (Cambridge, UK). The anti-human ADAMTS-5 catalytic domain rabbit polyclonal antibody was raised in rabbits and characterized [1].

*Validation of LRP1 ligand isolation using sLRP1-II and anti-c-Myc magnetic beads*

Purified recombinant ADAMTS5 lacking C-terminal TS domain (10 nM) and sLRP1-II (5 nM) was incubated in 500 µl of buffer consisting of 50 mM Tris-HCl (pH 7.5), 150 mM NaCl, 10 mM CaCl_2_, 0.01% BSA (TNCB buffer) and 20 µM CT1746 in the absence or presence of RAP (100 nM) for 1 h at 4°C. 30 µl of anti-c Myc antibody-conjugated magnetic Dynabeads was then added to the mixture and rotated for 30 min at 4°C. sLRP1-II and the magnetic beads complexes were then isolated using the magnetic stand and washed with TNCB buffer for three times. 40 µl of 2x SDS sampling buffer containing 5% mercaptoethanol, 30 µl of 0.1 M Glycine buffer (pH 3.0) or 30 µl of 500 nM RAP in TNCB buffer were added to the magnetic beads complexes and incubated at 95 °C for 5 min (for SDS) or at 4°C for 10 min (for Glycine and RAP). 10 µl of 4x SDS sampling buffer was further added to Glycine and RAP eluents followed by incubation at 95 °C for 5 min.

*Silver staining*

Purified proteins were separated under reducing conditions on a 4-12% Bis-Tris NuPage Gels (Thermo Fisher) and stained with a commercially available silver staining kit (Thermo scientific™, 24612). The gel was washed in ultrapure water and fixed in 30% ethanol:10% acetic acid solution twice for 15 minutes. The gel was washed in 10% ethanol solution and then sensitized in a Sensitizer Working Solution containing 50µL Sensitizer and 25mL of ultrapure water for 1 minute. Next, the gel was incubated with a previously prepared Stain Working Solution, containing 0.5mL of Enhancer and 25mL Stain solution for 30 minutes. Then a Developer Working Solution was prepared adding 0.5mL Enhancer to 25mL Developer solution. Finally, the gel was developed for 3-5 minutes until bands appeared and the reaction was stopped adding a 5% acetic acid solution for 10 minutes.

**<Figure Legends>**

**Suppl Fig 1. LRP1-mediated endocytic clearance of the endogenously produced CEMIP, SLIT2, HMGB2 and TSG6 in human chondrocytes.**

Human normal chondrocytes from three different donors were incubated with serum-free DMEM containing CT1746 (100 µM) and the protease inhibitor cocktail (1/500) in the absence or presence of 5 nM sLRP1-II for 24 h. The conditioned medium was collected and then subjected to Western blot analysis using anti-CEMIP (*A*), SLIT2 (*B*), HMGB2 (*C*), and TSG6 (*D*) antibodies. Band intensity was quantified by densitometry and the fold increase in the presence of sLRP1-II compared to the absence of sLRP1-II in each patient shown under the blot image. *Std*: 5 nM purified protein of CEMIP and 20 nM purified proteins of SLIT2, HMGB2 and TSG6.

**Suppl Fig 2. Validation of LRP1 ligand isolation using sLRP1-II and anti-c-Myc magnetic beads.**

*A*, 10 nM ADAMTS5 (ATS5) and 5 nM sLRP1-II were incubated in the presence or absence of 100 nM RAP for 1 h at 4°C followed by incubation with anti-cMyc antibody-conjugated magnetic Dynabeads for 30 min at 4°C. ADAMTS5 and sLRP1-II complexes were isolated using the magnetic stand, washed, eluted with SDS sampling buffer, and subjected to Western blot analysis using anti-ADAMTS5 antibody (*upper panel*) and anti-6x His antibody (*lower panel*). *B*, 10 nM ADAMTS5 and 5 nM sLRP1-II were incubated for 1 h at 4°C followed by incubation with anti-cMyc antibody-conjugated magnetic Dynabeads for 30 min at 4°C. ADAMTS5 and sLRP1-II complexes were isolated using the magnetic stand, washed, eluted with SDS sampling buffer, RAP (100 nM) or 0.1 M glycine buffer (pH 3.0) and subjected to Western blot analysis using anti-ADAMTS5 antibody. Asterisks indicate possible IgG proteins extracted from anti-cMyc antibody-conjugated magnetic Dynabeads by SDS and 2ME.

**Suppl Fig 3. SDS-PAGE analysis of proteins isolated with sLRP1-II from medium of human chondrocytes.**

Human normal chondrocytes from four different donors (HC1-4) were incubated with serum-free DMEM containing sLRP1-II (5 nM), metalloproteinase inhibitor CT1746 (100 µM) and the protease inhibitor cocktail (1/500) for 24 h. The conditioned medium was collected, further incubated with anti-cMyc magnetic Dynabeads, and sLRP1-II and magnetic beads complexes were isolated using the magnetic stand. Glycine buffer elution was then subjected to SDS-PAGE followed by silver staining (*upper panel*) or Western blot analysis using anti-6xHis antibody (*lower panel*). Asterisk indicates bovine serum albumin coated on the test tubes used to collect and store the samples.

**Suppl Fig 4. LRP1-mediated endocytic clearance of the endogenously produced ADAMTS1 and IGFBP7 in human chondrocytes.**

Human normal chondrocytes from three different donors were incubated with serum-free DMEM containing CT1746 (100 µM) and the protease inhibitor cocktail (1/500) in the absence or presence of 5 nM sLRP1-II for 24 h. The conditioned medium was collected and then subjected to Western blot analysis using anti-ADAMTS1 (*A*) and IGFBP7 (*B*) antibodies. Band intensity was quantified by densitometry and the fold increase in the presence of sLRP1-II compared to the absence of sLRP1-II in each patient shown under the blot image. *Std*: 20 nM purified proteins of ADAMTS1 and IGFBP7.
