## Supplementary figures for "Uncovering the ligandome of low-density lipoprotein receptor-related protein 1 in cartilage: a top-down approach to identify therapeutic targets"

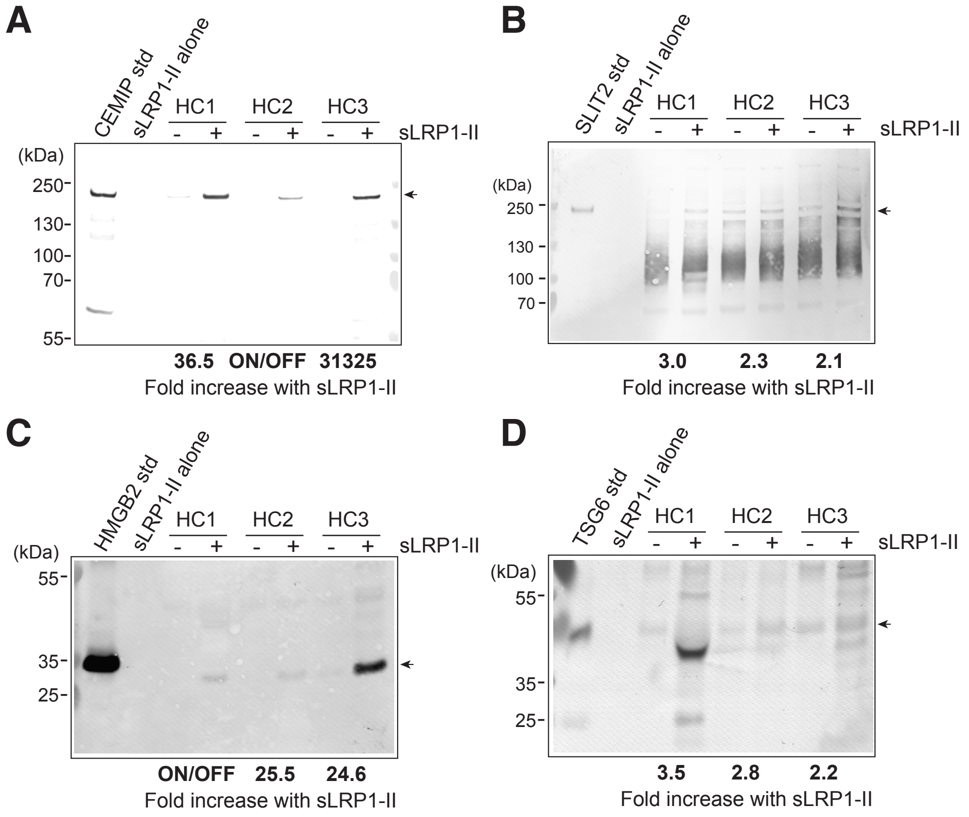


**Suppl Fig 1. LRP1-mediated endocytic clearance of endogenously produced CEMIP, SLIT2, HMGB2 and TSG6 in human chondrocytes**

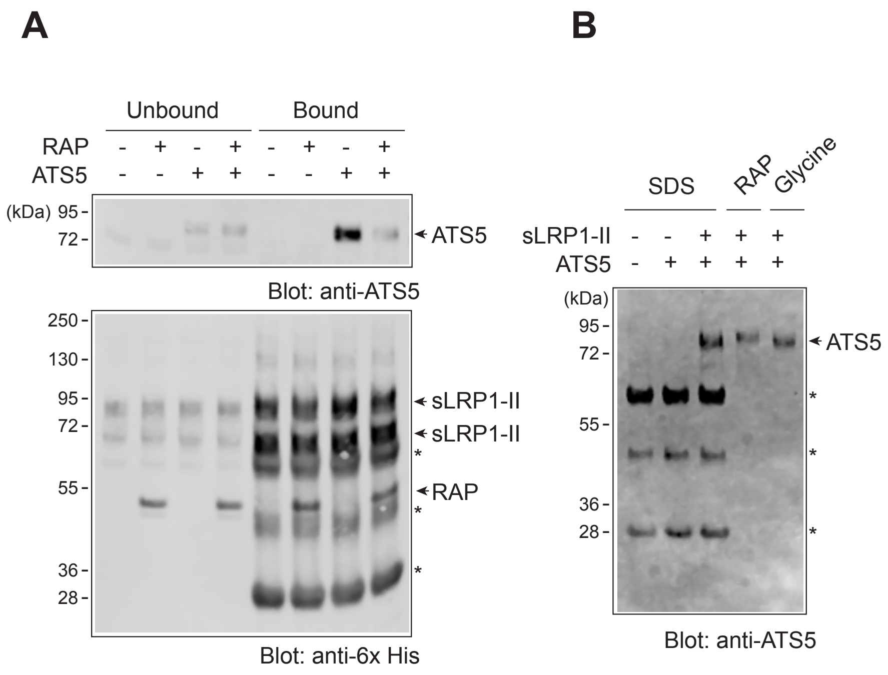


**Suppl Fig 2. Validation of LRP1 ligand isolation using sLRP1-II and anti-c-Myc magnetic beads.**

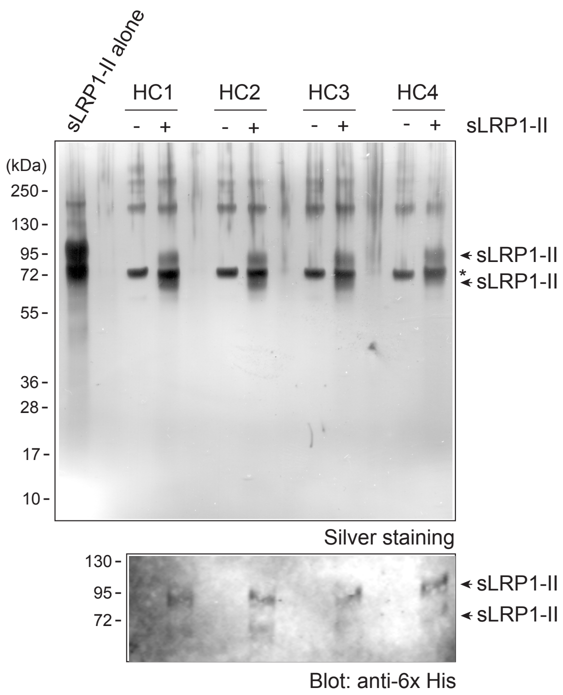


**Suppl Fig 3. SDS-PAGE analysis of proteins isolated with sLRP1-II from medium of human chondrocytes**

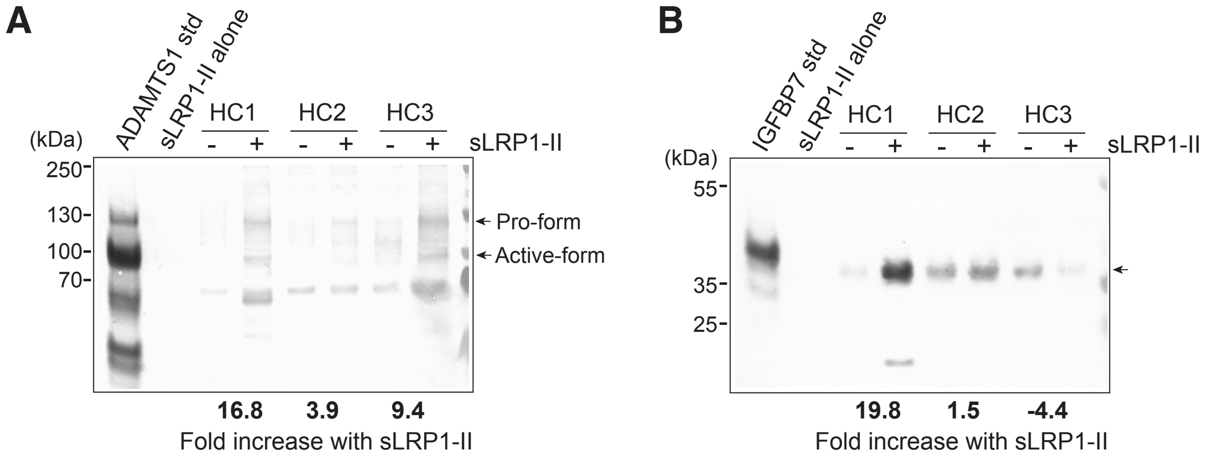


**Suppl Fig 4. LRP1-mediated endocytic clearance of endogenously produced ADAMTS1 and IGFBP7LRP1 ligands in human chondrocytes**
