## Supplementary table I for "Uncovering the ligandome of low-density lipoprotein receptor-related protein 1 in cartilage: a top-down approach to identify therapeutic targets"

**Suppl Table I: Molecules used for the volcano plot analysis**

| Gene name | $log_2$(LRP1-II/Control) | $-log_{10}$(P-value) |
|---|---|---|
| **Intracellular molecules_not significant** | | |
| HNRNPC | 2.80969453 | 0.992480887 |
| WDR1 | 2.32521605 | 1.298911451 |
| MAT2A | 2.123871565 | 1.132952184 |
| CRIP2 | 2.01084542 | 1.147718723 |
| UAP1 | 1.964226246 | 0.705985477 |
| SUMO3 | 1.91251159 | 1.021092502 |
| RPS24 | 1.874982595 | 1.06623677 |
| PSMA5 | 1.77738452 | 1.155051517 |
| SARS | 1.72631741 | 1.19098862 |
| PYGB | 1.71383381 | 1.264570979 |
| ARPC1B | 1.68907309 | 0.846618286 |
| PSMA3 | 1.66420937 | 0.714758542 |
| SPTBN1 | 1.65202141 | 1.138537876 |
| RPLP2 | 1.63459587 | 1.2731649 |
| LXN | 1.62528825 | 0.8026753 |
| ARPC2 | 1.61013079 | 1.119250053 |
| CSRP1 | 1.56855106 | 0.85550287 |
| ACTR2 | 1.51557636 | 1.058955758 |
| CCT6A | 1.46544099 | 0.992376263 |
| HNRNPA1 | 1.45958948 | 1.169096235 |
| XPO1 | 1.44817853 | 0.731887603 |
| EIF3CL | 1.43735647 | 0.576919184 |
| CAP1 | 1.43016171 | 0.778877983 |
| LSM3 | 1.41973114 | 1.186125386 |
| P4HA1 | 1.39885712 | 0.686911275 |
| TAGLN | 1.37353706 | 0.865651802 |
| GSTO1 | 1.35659266 | 0.866788073 |
| TGM2 | 1.32493973 | 0.997757512 |
| RBBP4 | 1.32029319 | 0.850529521 |
| HNRNPH1 | 1.29740906 | 0.441275629 |
| SUMF2 | 1.29432297 | 0.858254462 |
| RPS19 | 1.27973723 | 0.537133195 |
| ACTN4 | 1.26033497 | 1.232083516 |
| RBBP7 | 1.2528801 | 1.055334851 |
| PSMA7 | 1.24451923 | 1.087988991 |
| CS | 1.23286915 | 0.866946583 |
| RPL13A | 1.22792053 | 0.413922343 |
| PPP3CB | 1.22718859 | 0.550297934 |
| DDX17 | 1.22608423 | 0.990449761 |
| HMGA1 | 1.22076035 | 0.802201725 |
| GNPDA1 | 1.19988298 | 0.855219892 |
| XRCC6 | 1.17775369 | 0.398202626 |
| SEC31A | 1.17460394 | 0.479347794 |
| TUBB2A | 1.16128826 | 0.513522751 |
| YWHAG | 1.11852312 | 1.143789395 |
| TPT1 | 1.11699438 | 1.086909039 |
| CFL1 | 1.10340023 | 1.236970815 |
| NAP1L1 | 1.08601761 | 0.6016521 |
| TALDO1 | 1.08547735 | 1.175714235 |
| SERPINH1 | 1.07742739 | 1.025702017 |
| YWHAE | 1.06705284 | 1.253062421 |
| FKBP10 | 1.06585217 | 1.234021902 |
| AKR1B1 | 1.04372215 | 1.20225831 |
| COPG1 | 1.04351377 | 0.298544252 |
| SH3BGRL | 1.03431511 | 0.595332466 |
| HEBP2 | 1.0011425 | 1.159479378 |
| NARS | 0.97763991 | 0.471992906 |
| ACTN1 | 0.97657824 | 0.989649483 |
| SEPT7 | 0.97478318 | 0.723483709 |
| GDI2 | 0.9688344 | 0.881603266 |
| FKBP2 | 0.95782542 | 0.769496252 |
| PDLIM5 | 0.95515251 | 1.256456768 |
| LASP1 | 0.9505136 | 1.059940532 |
| EHD2 | 0.93758059 | 0.573009269 |
| ALDOA | 0.93433809 | 1.29960074 |
| TKT | 0.92615271 | 1.297874272 |
| PSMA1 | 0.91041613 | 0.979030098 |
| DYNC1I2 | 0.90594482 | 0.40659551 |
| HIST1H2BK | 0.90084743 | 1.061787105 |
| MYH9 | 0.89879608 | 1.25327992 |
| CALM1 | 0.8912425 | 0.794711712 |
| RPL12 | 0.86444592 | 0.395717108 |
| MDH1 | 0.8605094 | 0.598106355 |
| FKBP1A | 0.85088444 | 1.035247265 |
| PFN1 | 0.84363699 | 0.854493293 |
| CAPG | 0.83296108 | 1.032045644 |
| P4HB | 0.82715893 | 0.595810542 |
| TPM4 | 0.81297064 | 1.051621287 |
| SET | 0.80074215 | 1.061725306 |
| CAST | 0.79693294 | 0.34513143 |
| RPL23A | 0.79003167 | 0.381591034 |
| ANP32A | 0.75566101 | 0.986254329 |
| MSN | 0.73309374 | 1.242896377 |
| S100A11 | 0.7282939 | 0.949578996 |
| PRDX3 | 0.72523928 | 0.223811391 |
| H2AFX | 0.70766783 | 0.84988985 |
| TPI1 | 0.68116808 | 1.031020618 |
| PGAM1 | 0.67320824 | 1.177141567 |
| PSMA6 | 0.67015481 | 0.188028141 |
| TUBB | 0.65284109 | 1.078216679 |
| PEBP1 | 0.64715242 | 0.785007311 |
| GBE1 | 0.63912702 | 0.3237037 |
| HMOX1 | 0.62911344 | 0.330914114 |

| | | |
|---|---|---|
| PTMS | 0.62161398 | 0.664326268 |
| TPM1 | 0.61568332 | 0.160201638 |
| VCP | 0.59127474 | 0.480930277 |
| PEPD | 0.58894849 | 0.653807701 |
| SERPINB6 | 0.58184981 | 0.598609363 |
| LMNA | 0.56919336 | 0.558284423 |
| CPPED1 | 0.55917454 | 0.369788044 |
| PSMB7 | 0.55019569 | 0.673623527 |
| SNRPD3 | 0.5172348 | 0.608206403 |
| PDIA4 | 0.51010084 | 0.370142928 |
| PLBD2 | 0.50417852 | 1.01580449 |
| PDIA3 | 0.48454237 | 0.478565486 |
| PSMB5 | 0.47106862 | 0.190276082 |
| TXNDC5 | 0.45547438 | 0.354053645 |
| CYCS | 0.44709921 | 0.645237369 |
| PARK7 | 0.44539976 | 0.568557486 |
| DDAH2 | 0.43109012 | 0.148638959 |
| TWF1 | 0.38521051 | 0.140400538 |
| PRKCSH | 0.38003421 | 0.141160805 |
| GTF2A1 | 0.37155962 | 1.171419213 |
| BASP1 | 0.34929228 | 0.525172562 |
| RBMX | 0.33334589 | 0.390528335 |
| FKBP9 | 0.32425547 | 0.334847805 |
| PSMA4 | 0.30027199 | 0.385470802 |
| TBCA | 0.29079247 | 0.175572531 |
| RCN1 | 0.27738428 | 0.379922669 |
| PPIB | 0.25062084 | 0.219910936 |
| DDX39B | 0.24772978 | 0.561974781 |
| PSMD11 | 0.16760015 | 0.184864449 |
| S100A16 | 0.13458395 | 0.10724381 |
| ATP5B | 0.13209963 | 0.083126347 |
| VNN1 | 0.10099649 | 0.227181033 |
| COTL1 | 0.09297371 | 0.163318984 |
| STMN1 | 0.08760524 | 0.052978634 |
| DBNL | 0.08711243 | 0.061840785 |
| LPP | 0.07638645 | 0.209819793 |
| PTMA | 0.06119013 | 0.067759513 |
| SH3BGRL3 | -0.0018568 | 0.001548194 |
| TMSB4X | -0.0276475 | 0.012052764 |
| GORASP2 | -0.0721505 | 0.036153537 |
| LMNB2 | -0.0838938 | 0.078081228 |
| CAPZA1 | -0.0954063 | 0.045192167 |
| CTSL | -0.1135616 | 0.10154829 |
| STIP1 | -0.1530828 | 0.149675715 |
| HIST1H1D | -0.1568594 | 1.161853512 |
| PLEC | -0.2682867 | 0.735050445 |
| IMPA1 | -0.3404202 | 0.20929545 |
| KRT6B | -0.356992 | 0.1638152 |
| MT1G | -0.4211369 | 0.287231243 |
| OBSL1 | -0.4326446 | 0.277200418 |
| GDI1 | -0.5214055 | 0.370184224 |
| CTSZ | -0.5224686 | 1.100978974 |
| SDF4 | -0.613225 | 0.742560881 |
| PLOD2 | -0.8730011 | 0.847329069 |
| PSMB6 | -0.8734021 | 0.747737055 |
| KRT6C | -1.106771 | 0.356871758 |
| KRT5 | -1.2567363 | 1.113267862 |
| ARMC5 | -1.3076725 | 1.054574168 |
| CASP14 | -1.4351511 | 0.82916308 |
| KRT14 | -1.6814821 | 0.884119538 |
| **Intracellular molecules_significant** | | |
| RPS4X | 9.129591227 | 4.908750316 |
| RPS11 | 7.999605894 | 5.778012242 |
| LRPAP1 | 7.908970833 | 5.922735828 |
| RPS3A | 7.146104574 | 5.120989763 |
| SRP14 | 6.497942924 | 5.731250348 |
| PRDX5 | 6.48849678 | 4.137816183 |
| RPS26 | 6.35578728 | 4.688542726 |
| NRD1 | 5.588085413 | 4.446272485 |
| RPL7 | 5.31093001 | 3.627124258 |
| TLN1 | 4.85928893 | 2.779513239 |
| RPS25 | 4.74005795 | 6.085625854 |
| RPL27 | 4.54355264 | 3.426040342 |
| PGD | 4.51479697 | 2.876800294 |
| AMPD2 | 4.51473165 | 5.742075521 |
| CCT8 | 4.49792051 | 4.219328538 |
| UBA1 | 4.35392809 | 3.008919909 |
| ADSL | 4.353912115 | 4.671541702 |
| PPP1CB | 4.348917007 | 4.107411027 |
| RPS14 | 4.297822714 | 3.28027168 |
| HSPB1 | 4.27843714 | 2.039269646 |
| RPS12 | 4.13830829 | 2.985506178 |
| DYNC1H1 | 4.13750339 | 3.372069241 |
| SYNCRIP | 4.13178301 | 3.180176263 |
| CALD1 | 4.059571266 | 1.917540167 |
| CCT7 | 3.96440125 | 3.393525173 |
| RPL7A | 3.95437694 | 2.497573186 |
| HNRNPK | 3.95222139 | 3.388338801 |
| PDIA5 | 3.92341375 | 2.73110813 |
| HSPA4 | 3.9208653 | 3.284985947 |
| RSL1D1 | 3.848071814 | 6.198280009 |
| HMGB3 | 3.810292482 | 3.512300221 |
| RPL14 | 3.8089273 | 4.01729748 |
| RPL18 | 3.79409695 | 4.298015362 |
| RPL5 | 3.76040125 | 5.429943078 |
| PCBP2 | 3.6901896 | 1.82272218 |

| | | |
|---|---|---|
| EIF2S3 | 3.642795324 | 3.116570714 |
| PGM3 | 3.618304014 | 3.652423697 |
| VAT1 | 3.59194255 | 2.239015694 |
| ARF3 | 3.56597114 | 1.764151005 |
| RCN3 | 3.53401637 | 2.388213093 |
| PLIN3 | 3.50073123 | 3.785026858 |
| SNRPD2 | 3.500712156 | 2.981419948 |
| EIF2S1 | 3.50049591 | 2.25915385 |
| NQO1 | 3.46722746 | 1.3929291 |
| EEF1G | 3.46646595 | 3.470861217 |
| RPL4 | 3.42960286 | 2.620084539 |
| CLTC | 3.37479329 | 1.803295492 |
| SND1 | 3.35483432 | 2.237418904 |
| PCBP1 | 3.35162067 | 2.421639895 |
| SEPT11 | 3.30916405 | 3.404247574 |
| SRM | 3.23701978 | 2.402430356 |
| UGDH | 3.21711779 | 2.22076347 |
| COPB2 | 3.20025301 | 2.082156564 |
| ADIRF | 3.1899333 | 2.098474434 |
| ARF4 | 3.18224907 | 2.47839712 |
| AP2M1 | 3.15309095 | 2.965356722 |
| RPL29 | 3.12580633 | 3.868857355 |
| RPS18 | 3.05103254 | 2.116684614 |
| TARS | 3.02553606 | 2.239593191 |
| PA2G4 | 2.981220961 | 3.274758406 |
| VPS29 | 2.970266104 | 2.402660005 |
| TUBB4B | 2.96788645 | 2.578401899 |
| EEF1A1 | 2.94224024 | 2.601291308 |
| CAND1 | 2.93852472 | 3.152338143 |
| EIF3B | 2.93830395 | 2.571470975 |
| IARS | 2.91790652 | 2.263266471 |
| PABPC1 | 2.90823436 | 2.758353061 |
| IQGAP1 | 2.89406395 | 3.047323111 |
| CDC42 | 2.884329796 | 1.978026068 |
| CCT3 | 2.86429882 | 3.048756404 |
| FLNB | 2.86420321 | 1.452595639 |
| CNN3 | 2.84121394 | 2.399621791 |
| SRSF3 | 2.83201027 | 3.466313811 |
| SNRPF | 2.81538558 | 3.701738223 |
| HNRNPC | 2.809694529 | 0.992480887 |
| MYL12B | 2.80266976 | 1.849086959 |
| PHGDH | 2.75754666 | 2.160883429 |
| ARHGAP1 | 2.75039506 | 1.711901863 |
| RPL36 | 2.71735549 | 4.824374106 |
| ZYX | 2.69615436 | 1.96400492 |
| ADH5 | 2.68780613 | 2.648888094 |
| YWHAH | 2.66786289 | 1.742426146 |
| ARHGDIA | 2.66709518 | 1.617000154 |

| | | |
|---|---|---|
| PKM | 2.6610918 | 2.472365945 |
| AKR1C3 | 2.60455513 | 1.665970737 |
| GAPDH | 2.59980249 | 3.288999543 |
| YWHAQ | 2.57808924 | 2.076193688 |
| SRI | 2.56570244 | 2.066835353 |
| PRMT5 | 2.561795712 | 2.492918526 |
| RPL17 | 2.51393747 | 4.549725467 |
| RUVBL1 | 2.489670753 | 1.357286397 |
| DBI | 2.48042655 | 1.365005478 |
| ACTC1 | 2.4419384 | 2.037734199 |
| AP2A1 | 2.43135643 | 2.314091622 |
| EEF1D | 2.40869808 | 1.856957135 |
| HNRNPD | 2.38288116 | 2.922713894 |
| CA2 | 2.37807941 | 2.747391581 |
| AHCY | 2.3677578 | 3.232098527 |
| NNMT | 2.33156538 | 2.398968584 |
| RPL24 | 2.31170797 | 1.810684979 |
| EEF2 | 2.30029821 | 2.169206518 |
| U2AF2 | 2.29146791 | 1.806503369 |
| RPL32 | 2.257208109 | 2.195081697 |
| TPD52L2 | 2.253760099 | 1.440362411 |
| RAC1 | 2.25081253 | 2.365410237 |
| OTUB1 | 2.22252345 | 1.556978586 |
| EIF3J | 2.21892905 | 1.993333544 |
| VPS35 | 2.211839437 | 1.717247225 |
| NME1 | 2.20722055 | 3.044474338 |
| KPNB1 | 2.20668745 | 1.616544043 |
| PSME1 | 2.17546582 | 1.8285568 |
| FSCN1 | 2.16030979 | 1.910068035 |
| EPB41L2 | 2.15567303 | 2.771037381 |
| CAPN2 | 2.11180997 | 2.461524903 |
| MYL6 | 2.09302092 | 2.240187782 |
| XRCC5 | 2.08957815 | 2.168710572 |
| SF3B1 | 2.06810427 | 1.586049536 |
| PDIA6 | 2.03509951 | 1.489780719 |
| PRDX6 | 2.02806044 | 2.126654247 |
| EIF5A | 2.0206666 | 1.662388647 |
| FHL2 | 1.997776985 | 1.93626802 |
| CMPK1 | 1.97957611 | 1.733627988 |
| DSTN | 1.95957947 | 1.95653096 |
| SEC23B | 1.94585037 | 1.886752659 |
| ILF2 | 1.91347075 | 2.212992382 |
| SEPT9 | 1.90511131 | 1.630183009 |
| MAP4 | 1.89079428 | 1.487779385 |
| CAPZA2 | 1.86735344 | 2.108224228 |
| RPL9 | 1.850508451 | 3.034822734 |
| THY1 | 1.83840871 | 1.3985491 |
| AKR1C2 | 1.83231783 | 2.196010853 |

| | | |
|---|---|---|
| ILF3 | 1.80430913 | 1.73676817 |
| CAPZB | 1.76861811 | 1.830238217 |
| GNB1 | 1.73690939 | 1.994422766 |
| S100A10 | 1.73364758 | 2.256711111 |
| SELENBP1 | 1.71174097 | 1.813838996 |
| FAM129B | 1.69518304 | 1.82914613 |
| DDB1 | 1.68948269 | 1.785301348 |
| CORO1B | 1.68264771 | 1.861140727 |
| RPSA | 1.659513 | 1.718938374 |
| ARPC3 | 1.65141773 | 2.209280511 |
| RPLP0 | 1.61510706 | 1.668980042 |
| FLNA | 1.60632706 | 1.441643332 |
| UGP2 | 1.59790087 | 1.346987983 |
| HSPA8 | 1.58150959 | 1.772658261 |
| HSP90B1 | 1.57986879 | 1.73541446 |
| GANAB | 1.5795722 | 1.435642136 |
| MAPK1 | 1.5769453 | 1.42197645 |
| PSMD2 | 1.54308224 | 1.513159418 |
| ERP29 | 1.53232479 | 1.477429934 |
| DNASE2 | 1.52484655 | 3.523416275 |
| CNDP2 | 1.51848555 | 1.302548135 |
| HBD | 1.50600958 | 4.001921737 |
| ARPC4 | 1.49639273 | 1.473447312 |
| HNRNPA2B1 | 1.49627972 | 1.700780087 |
| VIM | 1.47045374 | 1.60317158 |
| ARPC5 | 1.44140816 | 1.482485879 |
| DPYSL2 | 1.43840885 | 1.322903998 |
| RNH1 | 1.43600655 | 1.595121132 |
| AK1 | 1.43130684 | 1.785186281 |
| PGK1 | 1.38167334 | 1.412250973 |
| EIF4A1 | 1.37650108 | 1.679432392 |
| MTPN | 1.35641074 | 1.510706282 |
| PCMT1 | 1.35212755 | 2.381764514 |
| DPYSL3 | 1.34984016 | 1.816553377 |
| AHNAK | 1.31570625 | 1.686195067 |
| GNS | 1.29399252 | 2.602711348 |
| PLS3 | 1.26227808 | 2.165859496 |
| PRDX2 | 1.25342941 | 1.649655718 |
| CORO1C | 1.24862957 | 1.311974531 |
| PGM1 | 1.24779367 | 1.854733862 |
| GLOD4 | 1.23865819 | 1.462567326 |
| GSTP1 | 1.23613453 | 1.39906651 |
| TPM3 | 1.23470879 | 1.580024614 |
| VCL | 1.23379993 | 1.78442184 |
| LDHA | 1.19718695 | 1.512500779 |
| PTBP1 | 1.19172907 | 1.357749142 |
| PSMB1 | 1.17531633 | 1.37961988 |
| LDHB | 1.17375469 | 1.51568758 |

| | | |
|---|---|---|
| SRSF1 | 1.17299747 | 1.753045264 |
| SERPIND1 | 1.16194534 | 1.838414455 |
| S100A4 | 1.13932991 | 2.082789584 |
| TSN | 1.13790202 | 1.461227685 |
| YWHAZ | 1.1199851 | 1.444973658 |
| RAD23B | 1.107687 | 1.969442694 |
| GNAS | 1.09704971 | 1.338240233 |
| PSAT1 | 1.07575464 | 1.70525312 |
| TAGLN2 | 1.07429695 | 1.489780809 |
| ACLY | 1.03177404 | 1.396567977 |
| RPS21 | 1.01737928 | 1.364501504 |
| FKBP3 | 1.00232863 | 2.494158392 |
| IDH1 | 0.99054623 | 1.391224956 |
| CSTB | 0.9742322 | 1.491444365 |
| ANXA5 | 0.91858768 | 1.91570531 |
| NCL | 0.90807819 | 1.417616617 |
| YWHAB | 0.9070549 | 1.567232495 |
| EZR | 0.89721441 | 1.456841984 |
| NT5E | 0.86600971 | 1.431880798 |
| MARCKS | 0.84180927 | 1.90800786 |
| MYH14 | 0.81223392 | 1.694828766 |
| AKR1A1 | 0.78878403 | 1.323357533 |
| RPS27A | 0.74321175 | 1.938921629 |
| CD109 | 0.66905451 | 1.940452705 |
| S100A6 | 0.64931107 | 1.669719821 |
| SOD1 | 0.49250126 | 1.419340447 |
| HIST1H1B | -0.3004284 | 1.8118935 |
| ACTR3 | -1.4546614 | 2.484924378 |
| NUFIP2 | -1.4626365 | 2.709688935 |
| WARS | -1.4791234 | 1.673024338 |
| CDH13 | -1.5165706 | 2.231767044 |
| EIF3D | -1.8305261 | 1.450613394 |
| THBS3 | -2.8148789 | 2.375563566 |
| C1S | -3.018652 | 3.358726011 |
| TK2 | -3.3465924 | 4.740151878 |
| **Secreted molecules_not significant** | | |
| MIF | 2.56294513 | 0.819733407 |
| PDCD6IP | 2.13630128 | 1.294052573 |
| YBX1 | 1.53289199 | 0.659914833 |
| SPON1 | 1.49089766 | 1.049522572 |
| RNPEP | 1.42623806 | 1.119860121 |
| PZP | 1.20555782 | 1.150550576 |
| FABP5 | 1.01819038 | 0.652289006 |
| SERPINB1 | 0.89653349 | 0.933291674 |
| MMP1 | 0.77135944 | 0.271853579 |
| MDK | 0.6840086 | 0.876430644 |
| GPI | 0.60200453 | 0.896900744 |

| | | |
|---|---|---|
| CP | 0.46252608 | 0.308313485 |
| KNG1 | 0.46197081 | 1.12716117 |
| LGALS3 | 0.44824314 | 0.521338262 |
| CALR | 0.42964602 | 0.409360103 |
| LTBP3 | 0.34979 | 0.62231 |
| FBLN2 | 0.2871871 | 0.744810344 |
| C7 | 0.27841091 | 1.171221263 |
| GPC6 | 0.25910807 | 0.091093564 |
| THBS1 | 0.24628258 | 0.340198668 |
| C3 | 0.22606802 | 1.219826826 |
| MYDGF | 0.21587086 | 0.150431614 |
| LTF | 0.1326437 | 0.537410023 |
| SERPINC1 | 0.03405285 | 0.11681606 |
| GSN | 0.01911354 | 0.029672094 |
| ITIH1 | -0.0111303 | 0.022423431 |
| ALB | -0.0403 | 0.15046 |
| IGF2 | -0.073925 | 0.218254711 |
| FBN2 | -0.1277215 | 0.073590814 |
| AFP | -0.1768699 | 0.650469364 |
| GC | -0.2041049 | 0.598743087 |
| A2M | -0.2091 | 0.70403 |
| SMOC1 | -0.2219 | 0.363 |
| STC2 | -0.2415 | 0.58225 |
| NID1 | -0.2947578 | 0.953078476 |
| PENK | -0.3021135 | 0.140973001 |
| LOXL2 | -0.3025684 | 0.32343303 |
| CALU | -0.3126559 | 0.532212083 |
| SERPINE1 | -0.3730912 | 0.788704421 |
| CFI | -0.4415646 | 0.922027639 |
| AHSG | -0.4601 | 0.47844 |
| NUCB2 | -0.4669352 | 0.570697352 |
| TGFBI | -0.5152221 | 0.960810028 |
| FST | -0.6964331 | 0.431370998 |
| IGFBP2 | -0.7002754 | 1.045273365 |
| IGFBP3 | -0.7974582 | 0.928190784 |
| GAS6 | -0.8453565 | 0.371446343 |
| PLG | -0.9073668 | 0.802059429 |
| MMP3 | -0.9300685 | 0.714383916 |
| PRELP | -0.9485271 | 0.533632606 |
| POSTN | -0.9937625 | 0.342480071 |
| PTX3 | -1.0522709 | 0.76718785 |
| IGFBP1 | -1.2893465 | 0.577637357 |
| KRT10 | -1.6248 | 1.10885 |
| SVEP1 | -1.755053 | 1.14072007 |
| CYTL1 | -2.1497145 | 0.665193499 |
| PLTP | -2.6789966 | 1.085750216 |
| NELL2 | 2.428266048 | 1.018040553 |
| PDCD6IP | 2.136301279 | 1.294052573 |

| **Secreted molecules_significant** | | |
|---|---|---|
| SEPP1 | 7.019131422 | 3.822261417 |
| CILP2 | 3.435349804 | 3.147217 |
| SLIT2 | 6.629853487 | 5.315784977 |
| TIMP3 | 6.341382504 | 5.200487121 |
| CEMIP | 5.398365736 | 4.437701478 |
| GARS | 4.68474054 | 3.039268526 |
| LIF | 4.42043757 | 2.856769172 |
| PLAT | 4.300270557 | 4.714877385 |
| HGFAC | 4.18743658 | 4.257666525 |
| GRN | 3.764694691 | 6.695692842 |
| CTGF | 3.73909807 | 4.045476178 |
| HMGB2 | 3.41777825 | 1.571150302 |
| SMOC2 | 2.83364606 | 1.302724641 |
| WNT11 | 2.776615858 | 2.499162026 |
| HSP90AA1 | 2.72889353 | 2.5567737 |
| HSP90AB1 | 2.62945604 | 2.356065438 |
| HMGB1 | 1.971817017 | 2.883026952 |
| ANXA1 | 1.83879089 | 2.544246146 |
| PPIA | 1.78183651 | 1.816062416 |
| GDF15 | 1.49591255 | 1.847513276 |
| TXN | 1.44658995 | 1.913372227 |
| LGALS1 | 1.34096146 | 1.502216078 |
| CCDC80 | 1.31165028 | 2.062941871 |
| TNFAIP6 | 1.30735064 | 1.504529976 |
| ANXA2 | 1.03050041 | 1.963492043 |
| C4A | 0.9058013 | 2.578344318 |
| TNC | 0.8460784 | 1.32991083 |
| PSAP | 0.78243732 | 2.473531398 |
| CTSD | 0.7197485 | 1.524914499 |
| ITIH2 | 0.51451254 | 1.522256691 |
| F2 | 0.50144768 | 1.79791027 |
| SERPINF2 | 0.38626719 | 1.816425911 |
| ISLR | -0.4204373 | 1.312531294 |
| HSPG2 | -0.6869 | 2.79065 |
| RBP4 | -0.7105851 | 1.431326124 |
| MFAP2 | -0.7356 | 1.97288 |
| SERPINE2 | -0.9557347 | 1.843927934 |
| NPC2 | -1.0038033 | 1.88225805 |
| MGP | -1.0258937 | 1.571305232 |
| LOX | -1.03406 | 1.93365219 |
| ATRN | -1.1422107 | 3.617323906 |
| COL16A1 | -1.1894107 | 2.372094859 |
| FBLN1 | -1.2065549 | 2.472338519 |
| CYR61 | -1.2397156 | 3.676363506 |
| LTBP1 | -1.2581916 | 1.688431206 |
| CTSB | -1.3064494 | 2.194887394 |
| LAMB2 | -1.3100386 | 3.474430914 |

| | | |
|---|---|---|
| EDIL3 | -1.3329124 | 2.465369109 |
| COL14A1 | -1.3490963 | 2.383076436 |
| ABI3BP | -1.3858314 | 2.003864177 |
| IGFBP5 | -1.4016986 | 1.323004667 |
| AGT | -1.4458253 | 1.441796748 |
| VCAN | -1.4515634 | 2.438470858 |
| COL1A1 | -1.4739795 | 3.96515009 |
| COL2A1 | -1.4879768 | 1.519523008 |
| OMD | -1.5691061 | 3.678400101 |
| EMILIN1 | -1.6084032 | 3.664637625 |
| FN1 | -1.6646461 | 4.6843694 |
| SERPING1 | -1.6766768 | 3.27038342 |
| MFGE8 | -1.6805983 | 2.289862021 |
| NUCB1 | -1.7237601 | 4.102663315 |
| COL6A3 | -1.7342277 | 2.768019072 |
| COL5A1 | -1.73948 | 1.555340621 |
| COL3A1 | -1.7490258 | 3.522188521 |
| HTRA1 | -1.7589993 | 2.832421643 |
| LTBP2 | -1.7668495 | 3.164700287 |
| NID2 | -1.8443375 | 4.550111995 |
| LOXL3 | -1.8666573 | 2.176528895 |
| CPXM2 | -1.8807096 | 2.161010188 |
| IGFBP7 | -1.8811569 | 3.392844363 |
| PCOLCE | -1.8841572 | 3.162712134 |
| DAG1 | -1.9164209 | 2.425077551 |
| ENPP2 | -1.9348598 | 2.614507543 |
| GALNT2 | -1.9384217 | 2.634756108 |
| ACAN | -1.9595366 | 3.19169659 |
| PTGDS | -1.9783857 | 2.110948358 |
| COL6A1 | -2.0213442 | 3.736201585 |
| COL6A2 | -2.0579467 | 3.734825555 |
| CLU | -2.065405369 | 3.427260581 |
| SPOCK1 | -2.0944986 | 2.889721129 |
| CLEC3B | -2.1412215 | 4.214013632 |
| PRG4 | -2.1603823 | 2.442282732 |
| PCOLCE2 | -2.2599587 | 3.245539011 |
| SPARC | -2.2850609 | 2.810249207 |
| CCBE1 | -2.2941813 | 3.344555785 |
| DCN | -2.3328786 | 3.323011139 |
| MMP2 | -2.3384471 | 2.3002382 |
| COL12A1 | -2.3416276 | 2.669860441 |
| COL4A2 | -2.3448942 | 2.132396465 |
| BGN | -2.395956 | 3.811639712 |
| CILP | -2.4315515 | 1.620677093 |
| FMOD | -2.4337559 | 2.670682476 |
| FNDC1 | -2.4755025 | 1.439646467 |
| CHI3L1 | -2.4761577 | 4.060857115 |
| TIMP2 | -2.479279 | 2.978874009 |

| | | |
|---|---|---|
| EFEMP1 | -2.497623 | 3.59218137 |
| B2M | -2.5264273 | 2.3996227 |
| QSOX1 | -2.5535812 | 4.354083557 |
| LAMA4 | -2.613349 | 3.722488866 |
| LUM | -2.6307864 | 3.578153615 |
| RNASE4 | -2.6823101 | 4.28227827 |
| SERPINA3 | -2.6882849 | 1.976575406 |
| IGFBP4 | -2.7043052 | 4.734540693 |
| FSTL1 | -2.7314339 | 4.469951871 |
| OLFML3 | -2.8097043 | 2.112723939 |
| CST3 | -2.8179369 | 2.987016823 |
| COL1A2 | -2.8454065 | 5.195893432 |
| FBN1 | -2.9035082 | 3.344618861 |
| TIMP1 | -2.9517517 | 4.700931535 |
| LAMC1 | -2.9554758 | 3.544161402 |
| C1R | -2.9631248 | 3.76197295 |
| ECM1 | -2.979404 | 4.754393823 |
| AEBP1 | -3.0334845 | 3.747386862 |
| COMP | -3.1317625 | 3.25208472 |
| CHI3L2 | -3.1446109 | 2.528897158 |
| COL5A2 | -3.1839347 | 5.030100896 |
| OGN | -3.1897621 | 3.837423099 |
| SOD3 | -3.3901095 | 3.663245423 |
| SERPINF1 | -3.4059801 | 3.598135857 |
| THBS2 | -3.4449344 | 1.549860218 |
| EFEMP2 | -3.4578483 | 2.377496351 |
| COL15A1 | -3.4753933 | 2.464243964 |
| IGFBP6 | -3.6055961 | 3.847438231 |
| LAMB1 | -3.6071296 | 4.857442361 |
| CFH | -3.9922667 | 2.638341264 |
| SPON2 | -4.0094528 | 2.48942365 |
| ITGBL1 | -4.2277875 | 3.095846321 |
| OLFML2B | -4.3851492 | 2.945430074 |
| CLEC11A | -4.4494994 | 2.517627515 |
| ADM | -4.5219951 | 2.902493195 |
| **Transmembrane molecules_not significant** | | |
| CD44 | 2.20159841 | 1.283502808 |
| RTN4 | 0.87357044 | 1.241479607 |
| LAMP2 | 0.60876656 | 0.763668636 |
| RRBP1 | 1.21383381 | 0.636800029 |
| ITGB1 | 0.8320322 | 0.4816218 |
| YIPF3 | 0.43291068 | 0.430002168 |
| **Transmembrane molecules_significant** | | |
| GLG1 | 6.6954 | 4.02626 |
| SLC39A10 | 4.55961 | 5.81473 |
| CLIC4 | 2.26634288 | 1.789459873 |
| VCAM1 | 1.67995739 | 2.100222949 |

| | | |
|---|---|---|
| ABCA10 | 1.63057756 | 2.266909105 |
| MMP14 | 1.29195642 | 2.427535151 |
| CLIC1 | 1.1505332 | 1.603779951 |
| APP | 1.07238078 | 2.12310456 |
| ANPEP | 0.93050194 | 2.110770484 |
| CD59 | 0.86198139 | 1.478180339 |
| CLSTN1 | -0.903 | 4.35369 |
| ATRN | -1.1422107 | 3.617323906 |
| MAN1A1 | -1.4525275 | 2.508820903 |
| DAG1 | -1.9164209 | 2.425077551 |
| GALNT2 | -1.9384217 | 2.634756108 |
| GOLM1 | -2.0109653 | 3.981397004 |
| QSOX1 | -2.5535812 | 4.354083557 |

* Molecules only identified with or without sLRP1-II were excluded
