## Supplementary table II for "Uncovering the ligandome of low-density lipoprotein receptor-related protein 1 in cartilage: a top-down approach to identify therapeutic targets"

Suppl Table II: Molecules increased in medium of human chondrocytes after 24-h incubation with sLRP1-II

| Description | Gene name | Accession | Ratio (sLRP1-II vs control) | p-value |
|---|---|---|---|---|
| **Proteins identified only with sLRP1-II** | | | | |
| Nardilysin | NRD1 | O43847 | | |
| 60S ribosomal protein L13 | RPL13 | P26373 | | |
| 60S ribosomal protein L36a | RPL36A | P83881 | | |
| 60S ribosomal protein L30 | RPL30 | P62888 | | |
| T-complex protein 1 subunit delta | CCT4 | P50991 | | |
| 40S ribosomal protein S6 | RPS6 | P62753 | | |
| 60S ribosomal protein L8 | RPL8 | P62917 | | |
| Adenylosuccinate lyase | ADSL | P30566 | | |
| Serine/threonine-protein phosphatase PP1-beta catalytic subu | PPP1CB | P62140 | | |
| 40S ribosomal protein S14 | RPS14 | P62263 | | |
| Splicing factor 3B subunit 2 | SF3B2 | Q13435 | | |
| 60S ribosomal protein L18a | RPL18A | Q02543 | | |
| Caldesmon | CALD1 | Q05682 | | |
| Ribosomal L1 domain-containing protein 1 | RSL1D1 | O76021 | | |
| High mobility group protein B3 | HMGB3 | O15347 | | |
| Endoplasmic reticulum resident protein 44 | ERP44 | Q9BS26 | | |
| Eukaryotic translation initiation factor 2 subunit 3 | EIF2S3 | P41091 | | |
| Serine/threonine-protein kinase PRP4 homolog | PRPF4B | Q13523 | | |
| Phosphoacetylglucosamine mutase | PGM3 | O95394 | | |
| Small nuclear ribonucleoprotein Sm D2 | SNRPD2 | P62316 | | |
| 40S ribosomal protein S15 | RPS15 | P62841 | | |
| Multifunctional protein ADE2 | PAICS | P22234 | | |
| Serine/arginine-rich splicing factor 4 | SRSF4 | Q08170 | | |
| 60S ribosomal protein L15 | RPL15 | P61313 | | |
| 60S ribosomal protein L22 | RPL22 | P35268 | | |
| Adipogenesis regulatory factor | ADIRF | Q15847 | | |
| *Hemopexin | HPX | P02790 | | |
| Proliferation-associated protein 2G4 | PA2G4 | Q9UQ80 | | |
| 60S ribosomal protein L21 | RPL21 | P46778 | | |
| Vacuolar protein sorting-associated protein 29 | VPS29 | Q9UBQ0 | | |
| Receptor of activated protein C kinase 1 | GNB2L1 | P63244 | | |
| Cell division control protein 42 homolog | CDC42 | P60953 | | |
| FACT complex subunit SPT16 | SUPT16H | Q9Y5B9 | | |
| Small nuclear ribonucleoprotein F | SNRPF | P62306 | | |
| 60S ribosomal protein L36 | RPL36 | Q9Y3U8 | | |
| EMILIN-3 | EMILIN3 | Q9NT22 | | |
| Aldo-keto reductase family 1 member C3 | AKR1C3 | P42330 | | |
| Protein arginine N-methyltransferase 5 | PRMT5 | O14744 | | |
| RuvB-like 1 | RUVBL1 | Q9Y265 | | |
| 60S ribosomal protein L32 | RPL32 | P62910 | | |
| Tumor protein D54 | TPD52L2 | O43399 | | |
| Vacuolar protein sorting-associated protein 35 | VPS35 | Q96QK1 | | |
| Four and a half LIM domains protein 2 | FHL2 | Q14192 | | |
| UDP-N-acetylhexosamine pyrophosphorylase | UAP1 | Q16222 | | |
| tRNA-splicing ligase RtcB homolog | RTCB | Q9Y3I0 | | |
| 40S ribosomal protein S24 | RPS24 | P62847 | | |
| Sorting nexin-2 | SNX2 | O60749 | | |
| 60S ribosomal protein L9 | RPL9 | P32969 | | |
| Proteasome subunit alpha type-5 | PSMA5 | P28066 | | |
| Prostaglandin E synthase 3 | PTGES3 | Q15185 | | |
| **Proteins >2.5 fold increased with sLRP1-II** | | | | |
| 40S ribosomal protein S4, X isoform | RPS4X | P62701 | 560.12 | 7.42E-10 |
| 40S ribosomal protein S11 | RPS11 | P62280 | 255.93 | 1.00E-08 |

| | | | | |
|---|---|---|---|---|
| 40S ribosomal protein S3a | RPS3A | P61247 | 141.64 | 7.14E-08 |
| Selenoprotein P | SEPP1 | P49908 | 129.71 | 9.57E-08 |
| Golgi apparatus protein 1 | GLG1 | Q92896 | 103.64 | 2.02E-07 |
| Slit homolog 2 protein | SLIT2 | O94813 | 99.03 | 2.35E-07 |
| Signal recognition particle 14 kDa protein | SRP14 | P37108 | 90.38 | 3.18E-07 |
| Peroxiredoxin-5, mitochondrial | PRDX5 | P30044 | 89.79 | 3.25E-07 |
| 40S ribosomal protein S26 | RPS26 | P62854 | 81.90 | 4.41E-07 |
| *Metalloproteinase inhibitor 3 | TIMP3 | P35625 | 81.09 | 4.56E-07 |
| Signal recognition particle 9 kDa protein | SRP9 | P49458 | 76.95 | 5.42E-07 |
| Four and a half LIM domains protein 1 | FHL1 | Q13642 | 76.26 | 5.59E-07 |
| Stromal cell-derived factor 2-like protein 1 | SDF2L1 | Q9HCN8 | 71.05 | 7.07E-07 |
| 40S ribosomal protein S2 | RPS2 | P15880 | 62.42 | 1.09E-06 |
| 40S ribosomal protein S5 | RPS5 | P46782 | 58.81 | 1.32E-06 |
| 40S ribosomal protein S8 | RPS8 | P62241 | 48.87 | 2.45E-06 |
| 60S ribosomal protein L11 | RPL11 | P62913 | 42.81 | 3.80E-06 |
| Cell migration-inducing and hyaluronan-binding protein | CEMIP | Q8WUJ3 | 42.18 | 4.00E-06 |
| 60S ribosomal protein L36a | RPL36A | P83881 | 41.22 | 4.31E-06 |
| 60S ribosomal protein L7 | RPL7 | P18124 | 39.70 | 4.89E-06 |
| Guanosine-3',5'-bis(diphosphate) 3'-pyrophosphohydrolase MI | HDDC3 | Q8N4P3 | 38.89 | 5.23E-06 |
| 40S ribosomal protein S16 | RPS16 | P62249 | 37.02 | 6.17E-06 |
| 60S ribosomal protein L3 | RPL3 | P39023 | 34.67 | 7.66E-06 |
| Talin-1 | TLN1 | Q9Y490 | 29.03 | 1.38E-05 |
| Transcription initiation factor IIA subunit 2 | GTF2A2 | P52657 | 27.94 | 1.57E-05 |
| 60S ribosomal protein L30 | RPL30 | P62888 | 26.94 | 1.77E-05 |
| 40S ribosomal protein S25 | RPS25 | P62851 | 26.72 | 1.82E-05 |
| 60S ribosomal protein L27a | RPL27A | P46776 | 26.71 | 1.82E-05 |
| 60S ribosomal protein L6 | RPL6 | Q02878 | 26.68 | 1.83E-05 |
| Heat shock 70 kDa protein 1A | HSPA1A | P0DMV8 | 26.19 | 1.95E-05 |
| Non-POU domain-containing octamer-binding protein | NONO | Q15233 | 26.13 | 1.96E-05 |
| Glycine--tRNA ligase | GARS | P41250 | 25.72 | 2.07E-05 |
| T-complex protein 1 subunit delta | CCT4 | P50991 | 25.01 | 2.27E-05 |
| Zinc transporter ZIP10 | SLC39A10 | Q9ULF5 | 23.58 | 2.76E-05 |
| 40S ribosomal protein S23 | RPS23 | P62266 | 23.36 | 2.85E-05 |
| 60S ribosomal protein L27 | RPL27 | P61353 | 23.32 | 2.86E-05 |
| AMP deaminase 2 | AMPD2 | Q01433 | 22.86 | 3.06E-05 |
| 6-phosphogluconate dehydrogenase, decarboxylating | PGD | P52209 | 22.86 | 3.06E-05 |
| 40S ribosomal protein S6 | RPS6 | P62753 | 22.82 | 3.07E-05 |
| T-complex protein 1 subunit theta | CCT8 | P50990 | 22.59 | 3.18E-05 |
| 60S ribosomal protein L8 | RPL8 | P62917 | 21.48 | 3.76E-05 |
| Leukemia inhibitory factor | LIF | P15018 | 21.41 | 3.80E-05 |
| 40S ribosomal protein S15a | RPS15A | P62244 | 21.19 | 3.93E-05 |
| 40S ribosomal protein S3 | RPS3 | P23396 | 20.46 | 4.42E-05 |
| Ubiquitin-like modifier-activating enzyme 1 | UBA1 | P22314 | 20.45 | 4.43E-05 |
| *Tissue-type plasminogen activator | PLAT | P00750 | 19.70 | 5.01E-05 |
| *Heat shock protein beta-1 | HSPB1 | P04792 | 19.41 | 5.27E-05 |
| Hepatocyte growth factor activator | HGFAC | Q04756 | 18.22 | 6.49E-05 |
| 40S ribosomal protein S12 | RPS12 | P25398 | 17.61 | 7.27E-05 |
| Cytoplasmic dynein 1 heavy chain 1 | DYNC1H1 | Q14204 | 17.60 | 7.29E-05 |
| Heterogeneous nuclear ribonucleoprotein Q | SYNCRIP | O60506 | 17.53 | 7.38E-05 |
| Splicing factor 3B subunit 2 | SF3B2 | Q13435 | 17.22 | 7.83E-05 |
| 60S ribosomal protein L18a | RPL18A | Q02543 | 16.83 | 8.46E-05 |
| WD repeat-containing protein 5 | WDR5 | P61964 | 16.71 | 8.66E-05 |
| T-complex protein 1 subunit eta | CCT7 | Q99832 | 15.61 | 1.09E-04 |
| 60S ribosomal protein L7a | RPL7A | P62424 | 15.50 | 1.11E-04 |
| Heterogeneous nuclear ribonucleoprotein K | HNRNPK | P61978 | 15.48 | 1.12E-04 |
| Protein disulfide-isomerase A5 | PDIA5 | Q14554 | 15.17 | 1.19E-04 |
| *Heat shock 70 kDa protein 4 | HSPA4 | P34932 | 15.15 | 1.20E-04 |
| 40S ribosomal protein S27 | RPS27 | P42677 | 14.67 | 1.33E-04 |
| 60S ribosomal protein L10a | RPL10A | P62906 | 14.32 | 1.45E-04 |

| | | | | |
|---|---|---|---|---|
| 60S ribosomal protein L14 | RPL14 | P50914 | 14.02 | 1.55E-04 |
| 60S ribosomal protein L18 | RPL18 | Q07020 | 13.87 | 1.61E-04 |
| Progranulin | GRN | P28799 | 13.59 | 1.72E-04 |
| 60S ribosomal protein L5 | RPL5 | P46777 | 13.55 | 1.74E-04 |
| *Connective tissue growth factor | CTGF | P29279 | 13.35 | 1.82E-04 |
| Poly(rC)-binding protein 2 | PCBP2 | Q15366 | 12.91 | 2.04E-04 |
| Endoplasmic reticulum resident protein 44 | ERP44 | Q9BS26 | 12.90 | 2.04E-04 |
| GTP-binding nuclear protein Ran | RAN | P62826 | 12.83 | 2.08E-04 |
| Serine/threonine-protein kinase PRP4 homolog | PRPF4B | Q13523 | 12.48 | 2.28E-04 |
| Synaptic vesicle membrane protein VAT-1 homolog | VAT1 | Q99536 | 12.06 | 2.56E-04 |
| 40S ribosomal protein S15 | RPS15 | P62841 | 11.89 | 2.68E-04 |
| ADP-ribosylation factor 3 | ARF3 | P61204 | 11.84 | 2.72E-04 |
| 40S ribosomal protein S20 | RPS20 | P60866 | 11.63 | 2.89E-04 |
| Reticulocalbin-3 | RCN3 | Q96D15 | 11.58 | 2.92E-04 |
| Perilipin-3 | PLIN3 | O60664 | 11.32 | 3.16E-04 |
| Eukaryotic translation initiation factor 2 subunit 1 | EIF2S1 | P05198 | 11.32 | 3.16E-04 |
| NAD(P)H dehydrogenase [quinone] 1 | NQO1 | P15559 | 11.06 | 3.41E-04 |
| Elongation factor 1-gamma | EEF1G | P26641 | 11.05 | 3.42E-04 |
| Multifunctional protein ADE2 | PAICS | P22234 | 10.97 | 3.51E-04 |
| 60S ribosomal protein L4 | RPL4 | P36578 | 10.77 | 3.72E-04 |
| Transcriptional repressor protein YY1 | YY1 | P25490 | 10.70 | 3.81E-04 |
| High mobility group protein B2 | HMGB2 | P26583 | 10.69 | 3.82E-04 |
| 60S ribosomal protein L15 | RPL15 | P61313 | 10.64 | 3.88E-04 |
| 60S ribosomal protein L22 | RPL22 | P35268 | 10.47 | 4.10E-04 |
| Clathrin heavy chain 1 | CLTC | Q00610 | 10.37 | 4.22E-04 |
| Staphylococcal nuclease domain-containing protein 1 | SND1 | Q7KZF4 | 10.23 | 4.42E-04 |
| Poly(rC)-binding protein 1 | PCBP1 | Q15365 | 10.21 | 4.45E-04 |
| Septin-11 | SEPT11 | Q9NVA2 | 9.91 | 4.91E-04 |
| Splicing factor, proline- and glutamine-rich | SFPQ | P23246 | 9.83 | 5.05E-04 |
| Spermidine synthase | SRM | P19623 | 9.43 | 5.79E-04 |
| UDP-glucose 6-dehydrogenase | UGDH | O60701 | 9.30 | 6.07E-04 |
| Sushi repeat-containing protein SRPX | SRPX | P78539 | 9.20 | 6.28E-04 |
| Coatomer subunit beta' | COPB2 | P35606 | 9.19 | 6.31E-04 |
| ADP-ribosylation factor 4 | ARF4 | P18085 | 9.08 | 6.57E-04 |
| AP-2 complex subunit mu | AP2M1 | Q96CW1 | 8.90 | 7.03E-04 |
| Cartilage intermediate layer protein 2 | CILP2 | Q8IUL8 | 8.86 | 7.12E-04 |
| 40S ribosomal protein S9 | RPS9 | P46781 | 8.80 | 7.28E-04 |
| Nucleophosmin | NPM1 | P06748 | 8.80 | 7.29E-04 |
| 60S ribosomal protein L29 | RPL29 | P47914 | 8.73 | 7.49E-04 |
| Tubulin alpha-1B chain | TUBA1B | P68363 | 8.72 | 7.50E-04 |
| 40S ribosomal protein S18 | RPS18 | P62269 | 8.29 | 8.89E-04 |
| Threonine--tRNA ligase, cytoplasmic | TARS | P26639 | 8.14 | 9.43E-04 |
| 60S ribosomal protein L21 | RPL21 | P46778 | 7.87 | 1.06E-03 |
| Tubulin beta-4B chain | TUBB4B | P68371 | 7.82 | 1.08E-03 |
| Elongation factor 1-alpha 1 | EEF1A1 | P68104 | 7.69 | 1.14E-03 |
| Cullin-associated NEDD8-dissociated protein 1 | CAND1 | Q86VP6 | 7.67 | 1.15E-03 |
| Eukaryotic translation initiation factor 3 subunit B | EIF3B | P55884 | 7.67 | 1.15E-03 |
| Peroxiredoxin-4 | PRDX4 | Q13162 | 7.61 | 1.18E-03 |
| Isoleucine--tRNA ligase, cytoplasmic | IARS | P41252 | 7.56 | 1.21E-03 |
| Polyadenylate-binding protein 1 | PABPC1 | P11940 | 7.51 | 1.24E-03 |
| Receptor of activated protein C kinase 1 | RACK1 | P63244 | 7.46 | 1.26E-03 |
| Ras GTPase-activating-like protein IQGAP1 | IQGAP1 | P46940 | 7.43 | 1.28E-03 |
| T-complex protein 1 subunit gamma | CCT3 | P49368 | 7.28 | 1.37E-03 |
| Filamin-B | FLNB | O75369 | 7.28 | 1.37E-03 |
| Calponin-3 | CNN3 | Q15417 | 7.17 | 1.44E-03 |
| SPARC-related modular calcium-binding protein 2 | SMOC2 | Q9H3U7 | 7.13 | 1.47E-03 |
| Serine/arginine-rich splicing factor 3 | SRSF3 | P84103 | 7.12 | 1.47E-03 |
| FACT complex subunit SPT16 | SUPT16H | Q9Y5B9 | 7.10 | 1.48E-03 |
| Myosin regulatory light chain 12B | MYL12B | O14950 | 6.98 | 1.58E-03 |

| | | | | |
|---|---|---|---|---|
| Protein Wnt-11 | WNT11 | O96014 | 6.85 | 1.67E-03 |
| D-3-phosphoglycerate dehydrogenase | PHGDH | O43175 | 6.76 | 1.75E-03 |
| Rho GTPase-activating protein 1 | ARHGAP1 | Q07960 | 6.73 | 1.78E-03 |
| T-complex protein 1 subunit alpha | TCP1 | P17987 | 6.50 | 1.99E-03 |
| Zyxin | ZYX | Q15942 | 6.48 | 2.01E-03 |
| Alcohol dehydrogenase class-3 | ADH5 | P11766 | 6.44 | 2.05E-03 |
| 14-3-3 protein eta | YWHAH | Q04917 | 6.35 | 2.15E-03 |
| Rho GDP-dissociation inhibitor 1 | ARHGDIA | P52565 | 6.35 | 2.15E-03 |
| Pyruvate kinase PKM | PKM | P14618 | 6.33 | 2.18E-03 |
| *Heat shock protein HSP 90-beta | HSP90AB1 | P08238 | 6.19 | 2.35E-03 |
| Glyceraldehyde-3-phosphate dehydrogenase | GAPDH | P04406 | 6.06 | 2.51E-03 |
| 14-3-3 protein theta | YWHAQ | P27348 | 5.97 | 2.64E-03 |
| Sorcin | SRI | P30626 | 5.92 | 2.72E-03 |
| Protein arginine N-methyltransferase 5 | PRMT5 | O14744 | 5.90 | 2.74E-03 |
| Heat shock protein HSP 90-alpha | HSP90AA1 | P07900 | 5.88 | 2.77E-03 |
| 60S ribosomal protein L17 | RPL17 | P18621 | 5.71 | 3.06E-03 |
| RuvB-like 1 | RUVBL1 | Q9Y265 | 5.62 | 3.24E-03 |
| Acyl-CoA-binding protein | DBI | P07108 | 5.58 | 3.31E-03 |
| Actin, alpha cardiac muscle 1 | ACTC1 | P68032 | 5.43 | 3.61E-03 |
| AP-2 complex subunit alpha-1 | AP2A1 | O95782 | 5.39 | 3.70E-03 |
| Elongation factor 1-delta | EEF1D | P29692 | 5.31 | 3.90E-03 |
| Heterogeneous nuclear ribonucleoprotein D0 | HNRNPD | Q14103 | 5.22 | 4.14E-03 |
| Carbonic anhydrase 2 | CA2 | P00918 | 5.20 | 4.19E-03 |
| Adenosylhomocysteinase | AHCY | P23526 | 5.16 | 4.29E-03 |
| Nicotinamide N-methyltransferase | NNMT | P40261 | 5.03 | 4.66E-03 |
| 60S ribosomal protein L24 | RPL24 | P83731 | 4.96 | 4.88E-03 |
| Elongation factor 2 | EEF2 | P13639 | 4.93 | 5.01E-03 |
| Splicing factor U2AF 65 kDa subunit | U2AF2 | P26368 | 4.90 | 5.11E-03 |
| Chloride intracellular channel protein 4 | CLIC4 | Q9Y696 | 4.81 | 5.42E-03 |
| Ras-related C3 botulinum toxin substrate 1 | RAC1 | P63000 | 4.76 | 5.61E-03 |
| Ubiquitin thioesterase OTUB1 | OTUB1 | Q96FW1 | 4.67 | 5.99E-03 |
| Eukaryotic translation initiation factor 3 subunit J | EIF3J | O75822 | 4.66 | 6.04E-03 |
| Nucleoside diphosphate kinase A | NME1 | P15531 | 4.62 | 6.21E-03 |
| Importin subunit beta-1 | KPNB1 | Q14974 | 4.62 | 6.21E-03 |
| Proteasome activator complex subunit 1 | PSME1 | Q06323 | 4.52 | 6.68E-03 |
| Fascin | FSCN1 | Q16658 | 4.47 | 6.91E-03 |
| Band 4.1-like protein 2 | EPB41L2 | O43491 | 4.46 | 6.99E-03 |
| Calpain-2 catalytic subunit | CAPN2 | P17655 | 4.32 | 7.73E-03 |
| Myosin light polypeptide 6 | MYL6 | P60660 | 4.27 | 8.07E-03 |
| X-ray repair cross-complementing protein 5 | XRCC5 | P13010 | 4.26 | 8.14E-03 |
| Splicing factor 3B subunit 1 | SF3B1 | O75533 | 4.19 | 8.55E-03 |
| Protein disulfide-isomerase A6 | PDIA6 | Q15084 | 4.10 | 9.22E-03 |
| Peroxiredoxin-6 | PRDX6 | P30041 | 4.08 | 9.37E-03 |
| Eukaryotic translation initiation factor 5A-1 | EIF5A | P63241 | 4.06 | 9.54E-03 |
| UMP-CMP kinase | CMPK1 | P30085 | 3.94 | 1.05E-02 |
| High mobility group protein B1 | HMGB1 | P09429 | 3.92 | 1.07E-02 |
| Destrin | DSTN | P60981 | 3.89 | 1.10E-02 |
| Protein transport protein Sec23B | SEC23B | Q15437 | 3.85 | 1.13E-02 |
| Interleukin enhancer-binding factor 2 | ILF2 | Q12905 | 3.77 | 1.22E-02 |
| Septin-9 | SEPT9 | Q9UHD8 | 3.75 | 1.24E-02 |
| Microtubule-associated protein 4 | MAP4 | P27816 | 3.71 | 1.29E-02 |
| Nucleoside diphosphate kinase B | NME2 | P22392 | 3.69 | 1.31E-02 |
| F-actin-capping protein subunit alpha-2 | CAPZA2 | P47755 | 3.65 | 1.36E-02 |
| Annexin A1 | ANXA1 | P04083 | 3.58 | 1.45E-02 |
| Thy-1 membrane glycoprotein | THY1 | P04216 | 3.58 | 1.45E-02 |
| Aldo-keto reductase family 1 member C2 | AKR1C2 | P52895 | 3.56 | 1.47E-02 |
| Interleukin enhancer-binding factor 3 | ILF3 | Q12906 | 3.49 | 1.57E-02 |
| Peptidyl-prolyl cis-trans isomerase A | PPIA | P62937 | 3.44 | 1.65E-02 |
| F-actin-capping protein subunit beta | CAPZB | P47756 | 3.41 | 1.70E-02 |

| | | | | |
|---|---|---|---|---|
| Guanine nucleotide-binding protein G(I)/G(S)/G(T) subunit beta | GNB1 | P62873 | 3.33 | 1.83E-02 |
| Protein S100-A10 | S100A10 | P60903 | 3.33 | 1.85E-02 |
| Methanethiol oxidase | SELENBP1 | Q13228 | 3.28 | 1.94E-02 |
| Niban-like protein 1 | FAM129B | Q96TA1 | 3.24 | 2.02E-02 |
| DNA damage-binding protein 1 | DDB1 | Q16531 | 3.23 | 2.04E-02 |
| Coronin-1B | CORO1B | Q9BR76 | 3.21 | 2.08E-02 |
| Vascular cell adhesion protein 1 | VCAM1 | P19320 | 3.20 | 2.09E-02 |
| 40S ribosomal protein SA | RPSA | P08865 | 3.16 | 2.19E-02 |
| Actin-related protein 2/3 complex subunit 3 | ARPC3 | O15145 | 3.14 | 2.23E-02 |
| ATP-binding cassette sub-family A member 10 | ABCA10 | Q8WWZ4 | 3.10 | 2.34E-02 |
| 60S acidic ribosomal protein P0 | RPLP0 | P05388 | 3.06 | 2.43E-02 |
| Endoplasmin | HSP90B1 | P14625 | 2.99 | 2.63E-02 |
| Heat shock cognate 71 kDa protein | HSPA8 | P11142 | 2.99 | 2.62E-02 |
| Deoxyribonuclease-2-alpha | DNASE2 | O00115 | 2.88 | 2.99E-02 |
| Actin, cytoplasmic 1 | ACTB | P60709 | 2.87 | 3.02E-02 |
| Hemoglobin subunit delta | HBD | P02042 | 2.84 | 3.12E-02 |
| Growth/differentiation factor 15 | GDF15 | Q99988 | 2.82 | 3.19E-02 |
| Thioredoxin | TXN | P10599 | 2.73 | 3.58E-02 |
| Hemoglobin subunit alpha | HBA1 | P69905 | 2.60 | 4.16E-02 |
| Peroxiredoxin-1 | PRDX1 | Q06830 | 2.58 | 4.30E-02 |
| Protein-L-isoaspartate(D-aspartate) O-methyltransferase | PCMT1 | P22061 | 2.55 | 4.45E-02 |

*LRP1 ligands already reported

Secreted proteins

Transmembrane proteins
