## Supplementary table III for "Uncovering the ligandome of low-density lipoprotein receptor-related protein 1 in cartilage: a top-down approach to identify therapeutic targets"

**Suppl Table III: Molecules decreased in medium of human chondrocytes after 24-h incubation with sLRP1-II**

| Description | Gene name | Accession | Ratio (control vs sLRP1-II) | p-value |
|---|---|---|---|---|
| **Proteins identified only without sLRP1-II** | | | | |
| Vitamin K-dependent protein S | PROS1 | P07225 | 16.88 | 8.31E-05 |
| Prostate-associated microseminoprotein | MSMP | Q1L6U9 | 16.01 | 1.48E-03 |
| Soluble scavenger receptor cysteine-rich domain-containing protein SSC5D | SSC5D | A1L4H1 | 15.27 | 2.21E-05 |
| Carboxypeptidase E | CPE | P16870 | 14.46 | 4.17E-04 |
| *Transforming growth factor beta-2 proprotein | TGFB2 | P61812 | 7.05 | 4.24E-03 |
| WNT1-inducible-signaling pathway protein 2 | WISP2 | O76076 | 5.97 | 4.16E-03 |
| **Proteins >2.5 fold deceased with sLRP1-II** | | | | |
| ADM | ADM | P35318 | 22.98 | 1.25E-03 |
| C-type lectin domain family 11 member A | CLEC11A | Q9Y240 | 21.85 | 3.04E-03 |
| Olfactomedin-like protein 2B | OLFML2B | Q68BL8 | 20.90 | 1.13E-03 |
| Integrin beta-like protein 1 | ITGBL1 | O95965 | 18.74 | 8.02E-04 |
| Spondin-2 | SPON2 | Q9BUD6 | 16.11 | 3.24E-03 |
| Complement factor H | CFH | P08603 | 15.91 | 2.30E-03 |
| Laminin subunit beta-1 | LAMB1 | P07942 | 12.19 | 1.39E-05 |
| Insulin-like growth factor-binding protein 6 | IGFBP6 | P24592 | 12.17 | 1.42E-04 |
| Collagen alpha-1(XV) chain | COL15A1 | P39059 | 11.12 | 3.43E-03 |
| EGF-containing fibulin-like extracellular matrix protein 2 | EFEMP2 | O95967 | 10.99 | 4.19E-03 |
| *Thrombospondin-2 | THBS2 | P35442 | 10.89 | 2.82E-02 |
| Pigment epithelium-derived factor | SERPINF1 | P36955 | 10.60 | 2.52E-04 |
| Extracellular superoxide dismutase [Cu-Zn] | SOD3 | P08294 | 10.48 | 2.17E-04 |
| Thymidine kinase 2, mitochondrial | TK2 | O00142 | 10.17 | 1.82E-05 |
| Mimecan | OGN | P20774 | 9.12 | 1.45E-04 |
| Collagen alpha-2(V) chain | COL5A2 | P05997 | 9.09 | 9.33E-06 |
| Chitinase-3-like protein 2 | CHI3L2 | Q15782 | 8.84 | 2.96E-03 |
| Cartilage oligomeric matrix protein | COMP | P49747 | 8.77 | 5.60E-04 |
| Adipocyte enhancer-binding protein 1 | AEBP1 | Q8IUX7 | 8.19 | 1.79E-04 |
| *Complement C1s subcomponent | C1S | P09871 | 8.10 | 4.38E-04 |
| Extracellular matrix protein 1 | ECM1 | Q16610 | 7.89 | 1.76E-05 |
| *Complement C1r subcomponent | C1R | P00736 | 7.80 | 1.73E-04 |
| Laminin subunit gamma-1 | LAMC1 | P11047 | 7.76 | 2.86E-04 |
| *Metalloproteinase inhibitor 1 | TIMP1 | P01033 | 7.74 | 1.99E-05 |
| Fibrillin-1 | FBN1 | P35555 | 7.48 | 4.52E-04 |
| Collagen alpha-2(I) chain | COL1A2 | P08123 | 7.19 | 6.37E-06 |
| Cystatin-C | CST3 | P01034 | 7.05 | 1.03E-03 |
| *Thrombospondin-3 | THBS3 | P49746 | 7.03 | 4.21E-03 |
| Olfactomedin-like protein 3 | OLFML3 | Q9NRN5 | 7.01 | 7.71E-03 |
| Follistatin-related protein 1 | FSTL1 | Q12841 | 6.64 | 3.39E-05 |
| Insulin-like growth factor-binding protein 4 | IGFBP4 | P22692 | 6.52 | 1.84E-05 |
| *Alpha-1-antichymotrypsin | SERPINA3 | P01011 | 6.45 | 1.06E-02 |
| Ribonuclease 4 | RNASE4 | P34096 | 6.42 | 5.22E-05 |
| Phospholipid transfer protein | PLTP | P55058 | 6.40 | 8.21E-02 |
| Lumican | LUM | P51884 | 6.19 | 2.64E-04 |
| Laminin subunit alpha-4 | LAMA4 | Q16363 | 6.12 | 1.89E-04 |
| Sulfhydryl oxidase 1 | QSOX1 | O00391 | 5.87 | 4.43E-05 |
| Beta-2-microglobulin | B2M | P61769 | 5.76 | 3.98E-03 |
| EGF-containing fibulin-like extracellular matrix protein 1 | EFEMP1 | Q12805 | 5.65 | 2.56E-04 |
| *Metalloproteinase inhibitor 2 | TIMP2 | P16035 | 5.58 | 1.05E-03 |
| Chitinase-3-like protein 1 | CHI3L1 | P36222 | 5.56 | 8.69E-05 |
| Fibronectin type III domain-containing protein 1 | FNDC1 | Q4ZHG4 | 5.56 | 3.63E-02 |
| Fibromodulin | FMOD | Q06828 | 5.40 | 2.13E-03 |
| Cartilage intermediate layer protein 1 | CILP | O75339 | 5.39 | 2.40E-02 |
| Biglycan | BGN | P21810 | 5.26 | 1.54E-04 |
| Collagen alpha-2(IV) chain | COL4A2 | P08572 | 5.08 | 7.37E-03 |
| Collagen alpha-1(XII) chain | COL12A1 | Q99715 | 5.07 | 2.14E-03 |
| *72 kDa type IV collagenase | MMP2 | P08253 | 5.06 | 5.01E-03 |
| *Decorin | DCN | P07585 | 5.04 | 4.75E-04 |
| Collagen and calcium-binding EGF domain-containing protein 1 | CCBE1 | Q6UXH8 | 4.90 | 4.52E-04 |
| *SPARC | SPARC | P09486 | 4.87 | 1.55E-03 |
| Procollagen C-endopeptidase enhancer 2 | PCOLCE2 | Q9UKZ9 | 4.79 | 5.68E-04 |
| Proteoglycan 4 | PRG4 | Q92954 | 4.47 | 3.61E-03 |
| Tetranectin | CLEC3B | P05452 | 4.41 | 6.11E-05 |
| Testican-1 | SPOCK1 | Q08629 | 4.27 | 1.29E-03 |
| Clusterin | CLU | P10909 | 4.19 | 3.74E-04 |

| | | | | |
|---|---|---|---|---|
| Collagen alpha-2(VI) chain | COL6A2 | P12110 | 4.16 | 1.84E-04 |
| Collagen alpha-1(VI) chain | COL6A1 | P12109 | 4.06 | 1.84E-04 |
| Golgi membrane protein 1 | GOLM1 | Q8NBJ4 | 4.03 | 1.04E-04 |
| Prostaglandin-H2 D-isomerase | PTGDS | P41222 | 3.94 | 7.75E-03 |
| Aggrecan core protein | ACAN | P16112 | 3.89 | 6.43E-04 |
| Polypeptide N-acetylgalactosaminyltransferase 2 | GALNT2 | Q10471 | 3.83 | 2.32E-03 |
| Ectonucleotide pyrophosphatase/phosphodiesterase family member 2 | ENPP2 | Q13822 | 3.82 | 2.43E-03 |
| Dystroglycan | DAG1 | Q14118 | 3.77 | 3.76E-03 |
| Insulin-like growth factor-binding protein 7 | IGFBP7 | Q16270 | 3.68 | 4.05E-04 |
| Procollagen C-endopeptidase enhancer 1 | PCOLCE | Q15113 | 3.69 | 6.88E-04 |
| Inactive carboxypeptidase-like protein X2 | CPXM2 | Q8N436 | 3.68 | 6.90E-03 |
| Lysyl oxidase homolog 3 | LOXL3 | P58215 | 3.65 | 6.66E-03 |
| Nidogen-2 | NID2 | Q14112 | 3.59 | 2.82E-05 |
| Eukaryotic translation initiation factor 3 subunit D | EIF3D | O15371 | 3.56 | 3.54E-02 |
| Latent-transforming growth factor beta-binding protein 2 | LTBP2 | Q14767 | 3.40 | 6.84E-04 |
| *Serine protease HTRA1 | HTRA1 | Q92743 | 3.38 | 1.47E-03 |
| Collagen alpha-1(III) chain | COL3A1 | P02461 | 3.36 | 3.00E-04 |
| Collagen alpha-1(V) chain | COL5A1 | P20908 | 3.34 | 2.78E-02 |
| Collagen alpha-3(VI) chain | COL6A3 | P12111 | 3.33 | 1.71E-03 |
| Nucleobindin-1 | NUCB1 | Q02818 | 3.30 | 7.89E-05 |
| Lactadherin | MFGE8 | Q08431 | 3.21 | 5.13E-03 |
| Plasma protease C1 inhibitor | SERPING1 | P05155 | 3.20 | 5.37E-04 |
| *Fibronectin | FN1 | P02751 | 3.17 | 2.07E-05 |
| EMILIN-1 | EMILIN1 | Q9Y6C2 | 3.05 | 2.16E-04 |
| Osteomodulin | OMD | Q99983 | 2.97 | 2.10E-04 |
| Cadherin-13 | CDH13 | P55290 | 2.86 | 5.86E-03 |
| Tryptophan--tRNA ligase | WARS | P23381 | 2.79 | 2.12E-02 |
| Collagen alpha-1(I) chain | COL1A1 | P02452 | 2.78 | 1.08E-04 |
| Nuclear fragile X mental retardation-interacting protein 2 | NUFIP2 | Q7Z417 | 2.76 | 1.95E-03 |
| Actin-related protein 3 | ACTR3 | P61158 | 2.74 | 3.27E-03 |
| Mannosyl-oligosaccharide 1,2-alpha-mannosidase IA | MAN1A1 | P33908 | 2.74 | 3.10E-03 |
| Versican core protein | VCAN | P13611 | 2.74 | 3.64E-03 |
| Collagen alpha-1(XIV) chain | COL14A1 | Q05707 | 2.55 | 4.14E-03 |
| EGF-like repeat and discoidin I-like domain-containing protein 3 | EDIL3 | O43854 | 2.52 | 3.42E-03 |

*LRP1 ligands already reported

Intracellular proteins

Transmembrane proteins
