## Supplementary table IV for "Uncovering the ligandome of low-density lipoprotein receptor-related protein 1 in cartilage: a top-down approach to identify therapeutic targets"

Suppl Table IV: Molecules co-immunoprecipitated with sLRP1-II from medium of human chondrocytes

| Description | Gene name | Accession | Ratio (sLRP1-II vs control) | p-value |
|---|---|---|---|---|
| **Proteins identified only with sLRP1-II** | | | | |
| WD repeat-containing protein 5 | WDR5 | P61964 | | |
| Cartilage intermediate layer protein 2 | CILP2 | Q8IUL8 | | |
| Signal recognition particle 9 kDa protein | SRP9 | P49458 | | |
| 60S ribosomal protein L36a | RPL36A | P83881 | | |
| 60S ribosomal protein L3 | RPL3 | P39023 | | |
| 60S ribosomal protein L21 | RPL21 | P46778 | | |
| Hepatocyte growth factor activator | HGFAC | Q04756 | | |
| Protein Wnt-11 | WNT11 | O96014 | | |
| Gamma-taxilin | TXLNG | Q9NUQ3 | | |
| High mobility group protein B1 | HMGB1 | P09429 | | |
| 60S ribosomal protein L23 | RPL23 | P62829 | | |
| Poly [ADP-ribose] polymerase 1 | PARP1 | P09874 | | |
| LIM and senescent cell antigen-like-containing domain protein 1 | LIMS1 | P48059 | | |
| 60S ribosomal protein L27a | RPL27A | P46776 | | |
| 60S ribosomal protein L17 | RPL17 | P18621 | | |
| Splicing factor 3B subunit 3 | SF3B3 | Q15393 | | |
| 60S ribosomal protein L18a | RPL18A | Q02543 | | |
| Nucleoside diphosphate kinase B | NME2 | P22392 | | |
| Structural maintenance of chromosomes protein 1A | SMC1A | Q14683 | | |
| Multifunctional protein ADE2 | PAICS | P22234 | | |
| Alpha-taxilin | TXLNA | P40222 | | |
| 60S ribosomal protein L32 | RPL32 | P62910 | | |
| Lysine--tRNA ligase | KARS | Q15046 | | |
| Stromal cell-derived factor 2-like protein 1 | SDF2L1 | Q9HCN8 | | |
| Adenylosuccinate lyase | ADSL | P30566 | | |
| AMP deaminase 2 | AMPD2 | Q01433 | | |
| Proprotein convertase subtilisin/kexin type 6 | PCSK6 | P29122 | | |
| FACT complex subunit SPT16 | SUPT16H | Q9Y5B9 | | |
| 60S ribosomal protein L7a | RPL7A | P62424 | | |
| 60S ribosomal protein L36 | RPL36 | Q9Y3U8 | | |
| High mobility group protein B2 | HMGB2 | P26583 | | |
| Eukaryotic translation initiation factor 3 subunit B | EIF3B | P55884 | | |
| T-complex protein 1 subunit beta | CCT2 | P78371 | | |
| Asparagine--tRNA ligase, cytoplasmic | NARS | O43776 | | |
| Coatomer subunit alpha | COPA | P53621 | | |
| Protein disulfide-isomerase A3 | PDIA3 | P30101 | | |
| Selenoprotein P | SELENOP | P49908 | | |
| Nucleoprotein TPR | TPR | P12270 | | |
| Dipeptidyl peptidase 9 | DPP9 | Q86TI2 | | |
| Nucleophosmin | NPM1 | P06748 | | |
| Splicing factor 3B subunit 1 | SF3B1 | O75533 | | |
| Creatine kinase B-type | CKB | P12277 | | |
| UPF0687 protein C20orf27 | C20orf27 | Q9GZN8 | | |
| Protein kinase C and casein kinase substrate in neurons protein 2 | PACSIN2 | Q9UNF0 | | |
| Interleukin enhancer-binding factor 2 | ILF2 | Q12905 | | |
| 60S ribosomal protein L35a | RPL35A | P18077 | | |
| Ubiquitin-like modifier-activating enzyme 1 | UBA1 | P22314 | | |
| Protein RCC2 | RCC2 | Q9P258 | | |
| Copine-3 | CPNE3 | O75131 | | |
| Serine--tRNA ligase, cytoplasmic | SARS | P49591 | | |
| Nascent polypeptide-associated complex subunit alpha, muscle-specific form | NACA | E9PAV3 | | |
| WD repeat-containing protein 1 | WDR1 | O75083 | | |
| Arginine--tRNA ligase, cytoplasmic | RARS | P54136 | | |
| Protein NDNF | NDNF | Q8TB73 | | |
| Heterogeneous nuclear ribonucleoprotein Q | SYNCRIP | O60506 | | |
| Protein DEK | DEK | P35659 | | |
| PHD finger-like domain-containing protein 5A | PHF5A | Q7RTV0 | | |
| Centrosomal protein of 135 kDa | CEP135 | Q66GS9 | | |
| Cysteine and glycine-rich protein 2 | CSRP2 | Q16527 | | |
| Splicing factor 3B subunit 2 | SF3B2 | Q13435 | | |
| F-BAR domain only protein 2 | FCHO2 | Q0JRZ9 | | |
| Cytokine receptor-like factor 1 | CRLF1 | O75462 | | |
| 60S ribosomal protein L37a | RPL37A | P61513 | | |
| Leucine--tRNA ligase, cytoplasmic | LARS | Q9P2J5 | | |
| Cytoplasmic FMR1-interacting protein 2 | CYFIP2 | Q96F07 | | |

| | | | | |
|---|---|---|---|---|
| Ribosomal L1 domain-containing protein 1 | RSL1D1 | O76021 | | |
| Carbonic anhydrase 2 | CA2 | P00918 | | |
| Aflatoxin B1 aldehyde reductase member 2 | AKR7A2 | O43488 | | |
| Zinc finger protein 768 | ZNF768 | Q9H5H4 | | |
| Ubiquitin carboxyl-terminal hydrolase 7 | USP7 | Q93009 | | |
| *Tissue factor pathway inhibitor | TFPI | P10646 | | |
| Eukaryotic translation initiation factor 3 subunit A | EIF3A | Q14152 | | |
| Eukaryotic translation initiation factor 3 subunit C-like protein | EIF3CL | B5ME19 | | |
| Dynein light chain 1, cytoplasmic | DYNLL1 | P63167 | | |
| Aspartate--tRNA ligase, cytoplasmic | DARS | P14868 | | |
| Thioredoxin domain-containing protein 5 | TXNDC5 | Q8NBS9 | | |
| S-formylglutathione hydrolase | ESD | P10768 | | |
| NAD(P)H dehydrogenase [quinone] 1 | NQO1 | P15559 | | |
| Deoxyribonuclease-2-alpha | DNASE2 | O00115 | | |
| Microtubule-associated protein 1A | MAP1A | P78559 | | |
| Epidermal growth factor-like protein 7 | EGFL7 | Q9UHF1 | | |
| 60S ribosomal protein L4 | RPL4 | P36578 | | |
| RNA-binding protein 25 | RBM25 | P49756 | | |
| Zinc finger CCCH-type antiviral protein 1-like | ZC3HAV1L | Q96H79 | | |
| Peroxiredoxin-2 | PRDX2 | P32119 | | |
| Eukaryotic translation initiation factor 2 subunit 1 | EIF2S1 | P05198 | | |
| Growth arrest-specific protein 6 | GAS6 | Q14393 | | |
| Platelet-activating factor acetylhydrolase IB subunit alpha | PAFAH1B1 | P43034 | | |
| 60S acidic ribosomal protein P2 | RPLP2 | P05387 | | |
| Neutral alpha-glucosidase AB | GANAB | Q14697 | | |
| Alpha-crystallin B chain | CRYAB | P02511 | | |
| Unconventional myosin-VI | MYO6 | Q9UM54 | | |
| S-methyl-5'-thioadenosine phosphorylase | MTAP | Q13126 | | |
| Adenosylhomocysteinase | AHCY | P23526 | | |
| Tubulin alpha-1A chain | TUBA1A | Q71U36 | | |
| Rho-associated protein kinase 2 | ROCK2 | O75116 | | |
| Protein disulfide-isomerase A4 | PDIA4 | P13667 | | |
| Septin-2 | SEPT2 | Q15019 | | |
| Serine/threonine-protein kinase TAO1 | TAOK1 | Q7L7X3 | | |
| Latent-transforming growth factor beta-binding protein 3 | LTBP3 | Q9NS15 | | |
| Ras-related C3 botulinum toxin substrate 1 | RAC1 | P63000 | | |
| ATP-dependent DNA helicase Q1 | RECQL | P46063 | | |
| T-complex protein 1 subunit epsilon | CCT5 | P48643 | | |
| Cdc42-interacting protein 4 | TRIP10 | Q15642 | | |
| Tubulin beta-2A chain | TUBB2A | Q13885 | | |
| SLIT-ROBO Rho GTPase-activating protein 2 | SRGAP2 | O75044 | | |
| Glycine--tRNA ligase | GARS | P41250 | | |
| ATP-binding cassette sub-family E member 1 | ABCE1 | P61221 | | |
| Cysteine-rich protein 2 | CRIP2 | P52943 | | |
| Testin | TES | Q9UGI8 | | |
| ELKS/Rab6-interacting/CAST family member 1 | ERC1 | Q8IUD2 | | |
| Heterogeneous nuclear ribonucleoprotein U | HNRNPU | Q00839 | | |
| Inner centromere protein | INCENP | Q9NQS7 | | |
| Signal recognition particle subunit SRP68 | SRP68 | Q9UHB9 | | |
| Putative RNA-binding protein Luc7-like 2 | LUC7L2 | Q9Y383 | | |
| **Proteins >2.5 fold increased with sLRP1-II** | | | | |
| 40S ribosomal protein S2 | RPS2 | P15880 | 1669.87 | 8.70E-08 |
| *Metalloproteinase inhibitor 3 | TIMP3 | P35625 | 1301.54 | 7.16E-06 |
| Signal recognition particle 14 kDa protein | SRP14 | P37108 | 1247.89 | 1.02E-10 |
| 40S ribosomal protein S16 | RPS16 | P62249 | 691.11 | 5.31E-06 |
| Golgi apparatus protein 1 | GLG1 | Q92896 | 661.65 | 8.26E-07 |
| Nucleoside diphosphate kinase A | NME1 | P15531 | 648.80 | 7.17E-06 |
| 60S ribosomal protein L5 | RPL5 | P46777 | 477.88 | 4.80E-06 |
| *Tissue-type plasminogen activator | PLAT | P00750 | 457.60 | 1.84E-08 |
| Four and a half LIM domains protein 1 | FHL1 | Q13642 | 444.09 | 6.55E-08 |
| Eukaryotic translation initiation factor 2 subunit 3 | EIF2S3 | P41091 | 342.33 | 1.36E-07 |
| Insulin-like growth factor-binding protein 7 | IGFBP7 | Q16270 | 293.50 | 1.04E-06 |
| *Connective tissue growth factor | CTGF | P29279 | 292.52 | 7.76E-06 |
| 60S ribosomal protein L10 | RPL10 | P27635 | 262.74 | 7.05E-08 |
| Cell migration-inducing and hyaluronan-binding protein | CEMIP | Q8WUJ3 | 236.61 | 9.10E-08 |
| 60S ribosomal protein L11 | RPL11 | P62913 | 225.13 | 1.47E-04 |
| A disintegrin and metalloproteinase with thrombospondin motifs 1 | ADAMTS1 | Q9UHI8 | 220.73 | 4.48E-08 |
| Myosin-9 | MYH9 | P35579 | 219.40 | 8.15E-05 |
| 60S ribosomal protein L8 | RPL8 | P62917 | 193.59 | 7.04E-08 |
| 40S ribosomal protein S26 | RPS26 | P62854 | 183.48 | 1.42E-04 |
| 40S ribosomal protein S3a | RPS3A | P61247 | 175.19 | 1.94E-08 |
| Microfibrillar-associated protein 2 | MFAP2 | P55001 | 174.38 | 1.09E-06 |
| 40S ribosomal protein S4, X isoform | RPS4X | P62701 | 160.35 | 3.27E-08 |

| | | | | |
|---|---|---|---|---|
| 60S ribosomal protein L14 | RPL14 | P50914 | 157.96 | 4.39E-06 |
| N-alpha-acetyltransferase 15, NatA auxiliary subunit | NAA15 | Q9BXJ9 | 155.70 | 6.91E-06 |
| Peroxiredoxin-1 | PRDX1 | Q06830 | 144.49 | 2.02E-05 |
| 40S ribosomal protein S5 | RPS5 | P46782 | 127.09 | 1.29E-04 |
| Protein disulfide-isomerase A5 | PDIA5 | Q14554 | 126.62 | 6.30E-05 |
| 40S ribosomal protein S3 | RPS3 | P23396 | 120.46 | 3.00E-08 |
| GTP-binding nuclear protein Ran | RAN | P62826 | 115.60 | 2.91E-06 |
| Heterogeneous nuclear ribonucleoprotein D0 | HNRNPD | Q14103 | 113.73 | 4.82E-05 |
| Myosin light polypeptide 6 | MYL6 | P60660 | 99.73 | 4.34E-05 |
| *SPARC | SPARC | P09486 | 96.43 | 5.76E-07 |
| T-complex protein 1 subunit gamma | CCT3 | P49368 | 95.57 | 1.14E-04 |
| Ras GTPase-activating-like protein IQGAP1 | IQGAP1 | P46940 | 92.37 | 2.66E-05 |
| 40S ribosomal protein S25 | RPS25 | P62851 | 92.21 | 1.72E-05 |
| T-complex protein 1 subunit zeta | CCT6A | P40227 | 81.72 | 1.09E-05 |
| 60S ribosomal protein L7 | RPL7 | P18124 | 79.52 | 1.16E-05 |
| 40S ribosomal protein S8 | RPS8 | P62241 | 79.47 | 2.16E-07 |
| 40S ribosomal protein S23 | RPS23 | P62266 | 79.26 | 3.46E-06 |
| AP-2 complex subunit mu | AP2M1 | Q96CW1 | 69.53 | 1.73E-04 |
| *Plasminogen activator inhibitor 1 | SERPINE1 | P05121 | 69.21 | 5.77E-07 |
| 60S ribosomal protein L6 | RPL6 | Q02878 | 66.93 | 1.25E-05 |
| T-complex protein 1 subunit theta | CCT8 | P50990 | 66.63 | 3.07E-06 |
| T-complex protein 1 subunit delta | CCT4 | P50991 | 64.09 | 1.11E-05 |
| Ubiquitin-40S ribosomal protein S27a | RPS27A | P62979 | 63.31 | 2.01E-05 |
| Splicing factor, proline- and glutamine-rich | SFPQ | P23246 | 60.00 | 3.08E-07 |
| *Protein CYR61 | CYR61 | O00622 | 59.25 | 6.92E-04 |
| Netrin-4 | NTN4 | Q9HB63 | 58.71 | 3.60E-06 |
| 40S ribosomal protein S11 | RPS11 | P62280 | 57.38 | 1.21E-08 |
| 60S ribosomal protein L10a | RPL10A | P62906 | 54.31 | 6.49E-07 |
| Thioredoxin | TXN | P10599 | 53.13 | 1.96E-05 |
| Golgin subfamily A member 3 | GOLGA3 | Q08378 | 53.03 | 1.49E-05 |
| Peroxiredoxin-4 | PRDX4 | Q13162 | 52.70 | 1.03E-04 |
| 60S ribosomal protein L9 | RPL9 | P32969 | 51.44 | 2.67E-05 |
| 60S ribosomal protein L24 | RPL24 | P83731 | 46.68 | 3.83E-04 |
| Receptor of activated protein C kinase 1 | RACK1 | P63244 | 44.59 | 5.88E-06 |
| T-complex protein 1 subunit alpha | TCP1 | P17987 | 42.20 | 1.94E-03 |
| Pre-mRNA-processing factor 40 homolog A | PRPF40A | O75400 | 40.62 | 2.18E-07 |
| Talin-1 | TLN1 | Q9Y490 | 40.29 | 5.08E-05 |
| Isoleucine--tRNA ligase, cytoplasmic | IARS | P41252 | 39.66 | 7.46E-06 |
| 40S ribosomal protein S14 | RPS14 | P62263 | 38.56 | 2.37E-03 |
| Guanosine-3',5'-bis(diphosphate) 3'-pyrophosphohydrolase MESH1 | HDDC3 | Q8N4P3 | 37.84 | 4.73E-04 |
| 60S ribosomal protein L19 | RPL19 | P84098 | 36.20 | 2.93E-06 |
| Heterogeneous nuclear ribonucleoprotein L | HNRNPL | P14866 | 36.00 | 1.09E-03 |
| Endoplasmin | HSP90B1 | P14625 | 34.98 | 2.78E-04 |
| Sushi repeat-containing protein SRPX | SRPX | P78539 | 34.42 | 4.13E-06 |
| Coiled-coil domain-containing protein 80 | CCDC80 | Q76M96 | 34.00 | 4.86E-03 |
| 60S ribosomal protein L28 | RPL28 | P46779 | 33.69 | 5.88E-06 |
| 60S ribosomal protein L35 | RPL35 | P42766 | 33.34 | 4.64E-04 |
| Kinesin-1 heavy chain | KIF5B | P33176 | 32.51 | 3.91E-05 |
| Non-POU domain-containing octamer-binding protein | NONO | Q15233 | 32.13 | 2.74E-05 |
| 40S ribosomal protein S15a | RPS15A | P62244 | 31.63 | 7.50E-06 |
| Cysteine and glycine-rich protein 1 | CSRP1 | P21291 | 30.63 | 3.23E-04 |
| Fascin | FSCN1 | Q16658 | 27.37 | 2.69E-05 |
| Slit homolog 2 protein | SLIT2 | O94813 | 27.11 | 9.80E-08 |
| 60S ribosomal protein L29 | RPL29 | P47914 | 26.85 | 9.39E-04 |
| 60S ribosomal protein L13 | RPL13 | P26373 | 26.35 | 4.10E-05 |
| 40S ribosomal protein S20 | RPS20 | P60866 | 24.93 | 2.01E-07 |
| 40S ribosomal protein S27 | RPS27 | P42677 | 24.33 | 2.63E-06 |
| T-complex protein 1 subunit eta | CCT7 | Q99832 | 22.35 | 5.19E-05 |
| *Alpha-2-macroglobulin | A2M | P01023 | 22.02 | 5.28E-06 |
| Endoplasmic reticulum chaperone BiP | HSPA5 | P11021 | 19.81 | 9.74E-08 |
| 60S ribosomal protein L22 | RPL22 | P35268 | 19.20 | 8.54E-04 |
| 60S ribosomal protein L30 | RPL30 | P62888 | 19.12 | 1.25E-05 |
| Fructose-bisphosphate aldolase A | ALDOA | P04075 | 17.95 | 4.98E-04 |
| Signal recognition particle subunit SRP72 | SRP72 | O76094 | 17.30 | 1.02E-03 |
| *Heat shock protein beta-1 | HSPB1 | P04792 | 16.63 | 3.56E-02 |
| Endoplasmic reticulum resident protein 44 | ERP44 | Q9BS26 | 16.58 | 1.01E-02 |
| 60S ribosomal protein L23a | RPL23A | P62750 | 16.56 | 5.91E-04 |
| 60S ribosomal protein L31 | RPL31 | P62899 | 16.35 | 1.99E-04 |
| Tumor necrosis factor-inducible gene 6 protein | TNFAIP6 | P98066 | 16.24 | 1.22E-03 |
| 60S ribosomal protein L38 | RPL38 | P63173 | 15.96 | 5.19E-04 |
| *Heat shock protein HSP 90-alpha | HSP90AA1 | P07900 | 15.61 | 1.04E-02 |
| AP-2 complex subunit alpha-1 | AP2A1 | O95782 | 15.21 | 2.99E-04 |
| AP-3 complex subunit beta-1 | AP3B1 | O00203 | 15.14 | 8.31E-05 |

| | | | | |
|---|---|---|---|---|
| Myosin regulatory light chain 12B | MYL12B | O14950 | 14.86 | 4.83E-04 |
| 40S ribosomal protein S6 | RPS6 | P62753 | 14.61 | 7.79E-06 |
| 60S ribosomal protein L18 | RPL18 | Q07020 | 14.58 | 1.74E-07 |
| Aldo-keto reductase family 1 member C2 | AKR1C2 | P52895 | 14.28 | 2.71E-05 |
| AP-2 complex subunit beta | AP2B1 | P63010 | 13.58 | 3.51E-04 |
| Tubulin beta chain | TUBB | P07437 | 13.46 | 6.39E-04 |
| 40S ribosomal protein S15 | RPS15 | P62841 | 13.34 | 4.39E-04 |
| Clathrin heavy chain 1 | CLTC | Q00610 | 13.19 | 6.42E-03 |
| LIM and SH3 domain protein 1 | LASP1 | Q14847 | 13.18 | 7.21E-06 |
| Glyceraldehyde-3-phosphate dehydrogenase | GAPDH | P04406 | 13.14 | 1.66E-04 |
| Amidophosphoribosyltransferase | PPAT | Q06203 | 12.50 | 1.36E-03 |
| Beta-2-microglobulin | B2M | P61769 | 12.49 | 1.32E-02 |
| 60S ribosomal protein L27 | RPL27 | P61353 | 11.97 | 1.90E-06 |
| UDP-glucose 6-dehydrogenase | UGDH | O60701 | 11.80 | 1.80E-03 |
| Filamin-A | FLNA | P21333 | 11.55 | 9.30E-04 |
| Anosmin-1 | ANOS1 | P23352 | 11.40 | 4.43E-08 |
| Nidogen-1 | NID1 | P14543 | 11.30 | 2.23E-03 |
| *Pregnancy zone protein | PZP | P20742 | 11.00 | 2.35E-06 |
| Tubulin beta-4B chain | TUBB4B | P68371 | 10.98 | 5.89E-04 |
| F-actin-capping protein subunit alpha-2 | CAPZA2 | P47755 | 10.47 | 1.65E-04 |
| Elongation factor 1-gamma | EEF1G | P26641 | 10.28 | 8.50E-05 |
| 40S ribosomal protein S7 | RPS7 | P62081 | 9.69 | 5.09E-03 |
| *Metalloproteinase inhibitor 1 | TIMP1 | P01033 | 9.51 | 1.47E-03 |
| 40S ribosomal protein S9 | RPS9 | P46781 | 9.38 | 5.79E-06 |
| Tubulin alpha-1B chain | TUBA1B | P68363 | 9.27 | 1.07E-04 |
| Inter-alpha-trypsin inhibitor heavy chain H2 | ITIH2 | P19823 | 9.20 | 6.03E-03 |
| L-lactate dehydrogenase A chain | LDHA | P00338 | 9.16 | 3.84E-07 |
| Elongation factor 2 | EEF2 | P13639 | 8.77 | 2.25E-06 |
| 40S ribosomal protein SA | RPSA | P08865 | 8.60 | 1.38E-02 |
| 40S ribosomal protein S21 | RPS21 | P63220 | 8.35 | 1.45E-02 |
| Pyruvate kinase PKM | PKM | P14618 | 8.33 | 3.50E-04 |
| Elastin | ELN | P15502 | 8.28 | 3.55E-02 |
| Legumain | LGMN | Q99538 | 8.27 | 6.18E-04 |
| Progranulin | GRN | P28799 | 8.24 | 3.41E-05 |
| Actin, alpha cardiac muscle 1 | ACTC1 | P68032 | 8.17 | 3.61E-03 |
| Heat shock 70 kDa protein 1A | HSPA1A | P0DMV8 | 8.01 | 3.06E-06 |
| 40S ribosomal protein S13 | RPS13 | P62277 | 7.91 | 8.68E-04 |
| Alpha-enolase | ENO1 | P06733 | 7.79 | 8.46E-05 |
| Collagen alpha-1(XIV) chain | COL14A1 | Q05707 | 7.56 | 1.09E-03 |
| Spliceosome RNA helicase DDX39B | DDX39B | Q13838 | 7.38 | 1.04E-04 |
| Elongation factor 1-alpha 1 | EEF1A1 | P68104 | 7.11 | 1.52E-04 |
| 40S ribosomal protein S24 | RPS24 | P62847 | 7.08 | 3.08E-03 |
| *Heat shock protein HSP 90-beta | HSP90AB1 | P08238 | 6.64 | 4.41E-06 |
| Pre-mRNA-splicing factor ATP-dependent RNA helicase DHX15 | DHX15 | O43143 | 6.34 | 1.24E-02 |
| Stanniocalcin-2 | STC2 | O76061 | 6.05 | 5.65E-05 |
| Hemoglobin subunit alpha | HBA1 | P69905 | 5.85 | 2.07E-05 |
| 40S ribosomal protein S30 | FAU | P62861 | 5.73 | 8.21E-03 |
| Actin, cytoplasmic 1 | ACTB | P60709 | 5.43 | 2.72E-03 |
| Phosphoglycerate kinase 1 | PGK1 | P00558 | 5.33 | 5.42E-03 |
| Fibulin-1 | FBLN1 | P23142 | 5.25 | 3.37E-03 |
| 40S ribosomal protein S18 | RPS18 | P62269 | 5.17 | 2.38E-04 |
| Gremlin-1 | GREM1 | O60565 | 5.12 | 1.10E-02 |
| 26S proteasome non-ATPase regulatory subunit 2 | PSMD2 | Q13200 | 5.01 | 1.23E-02 |
| 14-3-3 protein epsilon | YWHAE | P62258 | 4.98 | 2.64E-06 |
| Junction plakoglobin | JUP | P14923 | 4.57 | 5.72E-03 |
| Peptidyl-prolyl cis-trans isomerase A | PPIA | P62937 | 4.55 | 1.43E-03 |
| 60S acidic ribosomal protein P0 | RPLP0 | P05388 | 4.43 | 4.83E-02 |
| Plectin | PLEC | Q15149 | 4.33 | 1.01E-01 |
| Heterogeneous nuclear ribonucleoprotein K | HNRNPK | P61978 | 4.05 | 9.70E-04 |
| SPARC-related modular calcium-binding protein 2 | SMOC2 | Q9H3U7 | 4.03 | 7.43E-03 |
| *Complement C3 | C3 | P01024 | 3.89 | 1.30E-04 |
| Hemoglobin subunit delta | HBD | P02042 | 3.49 | 2.41E-06 |
| Phospholipase A2, membrane associated | PLA2G2A | P14555 | 3.35 | 2.38E-02 |
| Growth/differentiation factor 15 | GDF15 | Q99988 | 2.79 | 7.70E-03 |
| Lysyl oxidase homolog 2 | LOXL2 | Q9Y4K0 | 2.79 | 4.64E-04 |
| Protein Wnt-5a | WNT5A | P41221 | 2.73 | 3.08E-02 |

*LRP1 ligands already reported

Secreted proteins
